# Dissecting the functional heterogeneity of glutamatergic synapses with high-throughput optical physiology

**DOI:** 10.1101/2024.12.23.629904

**Authors:** Samuel T. Barlow, Aaron D. Levy, Minerva Contreras, Michael C. Anderson, Thomas A. Blanpied

**Affiliations:** Department of Physiology, University of Maryland School of Medicine, Baltimore, MD 21201, USA; UM-MIND, University of Maryland School of Medicine, Baltimore, MD 21201, USA; Program in Neuroscience, University of Maryland School of Medicine, Baltimore, MD 21201, USA; Institute for Cell Engineering, Johns Hopkins University School of Medicine, Baltimore, MD, USA; Department of Neurology, Johns Hopkins University School of Medicine, Baltimore, MD, USA

## Abstract

Fluorescent reporters for neurotransmitter release and Ca^2+^ signaling are essential tools for quantifying synapse function across neurons and circuits. However, leveraging these reporters requires scalable experimental frameworks. Here, we devised a high-throughput approach to efficiently collect and analyze hundreds of optical recordings of glutamate release activity using iGluSnFR3 at synapses in cultured rat hippocampal neurons. Our approach facilitated deep analysis of single-synapse properties following single action potentials, paired stimuli, or spontaneous release, and together with mean-variance analysis, we mapped synaptic functional diversity across the dendritic arbor. We then developed a novel all-optical assay of pre- and postsynaptic glutamatergic synapse function, deploying iGluSnFR3 along with a red-shifted, postsynaptically-targeted Ca^2+^ sensor to enable direct imaging and analysis of both NMDA receptor and AMPA receptor-mediated synaptic transmission at single dendritic spines. Direct observation of the flow of information at single synapses should speed detailed investigations of synaptic mechanistic heterogeneity.

## Introduction

Synapses are the fundamental information processing modules in the brain, performing computations that dictate how electrical activity propagates across neural circuits^1,2^. Thus, a major goal for neuroscience is to identify the basic functional properties of individual synapses which define their computational output, such as vesicle release probability (*P_r_*)^3^, the magnitude and variance of receptor activation, and short-term plasticity behavior^4^. However, the enormous diversity that exists among synapses is a significant barrier to achieving a quantitative understanding of synaptic function. The distinct transcriptomic identities of pre- and postsynaptic neurons drive expansive proteomic diversity among synapses^5^, and synapses are also plastic, with further speciation emanating from each synapse’s unique history of activity^6^. Synapse functional diversity is reflective of this deep proteomic diversity, with *P_r_* varying widely between synapse types (0.05 to 1)^3^. *P_r_* can fluctuate across stimulus frequencies as new vesicle populations^7^ or short-term plasticity mechanisms are engaged^4^, properties which also exhibit a dependence on synapse identity^7,8^. How these permutations of presynaptic properties impact the activation probability of postsynaptic receptors will define which patterns of activity lead to synapse strengthening or weakening^9^, constituting another axis of synaptic heterogeneity with implications for neural circuit development.

Due to their expansive diversity, a quantitative understanding of synaptic communication across single neurons and circuits will only be achieved through a synapse-by-synapse readout of synaptic function. Optical methods are well-positioned to meet this need, and direct measurement of synaptic functional properties has been demonstrated using a variety of fluorescent indicators^10,11,12,13,14^. However, several barriers must be overcome to leverage these tools at scale. First, detailed dissection of synaptic function requires a variety of stimulation protocols, chemical conditions, and imaging modalities, resulting in complex experimental paradigms. To acquire these data efficiently and reproducibly, it is desirable to fully automate microscopy, electrical stimulators, and fluidics. Second, to process large datasets, automated segmentation methods that can extract and analyze the same synapses across hundreds of video recordings are essential. Third, intensity-time recordings from individual synapses must be baseline-corrected and normalized to ΔF/F before fluorescence signals can be extracted and analyzed. Finally, fluorescence data must be converted to interpretable statistics for insight into synaptic functional properties. Indeed, major software packages have been developed to accelerate segmentation and fluorescence signal extraction for calcium (e.g. CaImAn^15^, FIOLA^16^) and voltage imaging (e.g. VolPy^17^) *in vivo*. More recent work has demonstrated high-throughput analysis of calcium transients at single-synapses^18^, but new software approaches are needed to comprehensively analyze both pre- and postsynaptic function with single-synapse resolution.

Here, we set out to develop all-optical methods to measure the functional heterogeneity of single synapses in cultured rat hippocampal neurons. We first focused on the third-generation intensity-based glutamate sensing fluorescent reporter (iGluSnFR3)^19^, which allows robust detection of glutamate release from single presynapses to provide quantitative access to basic functional properties such as basal release probability^20^,short-term plasticity dynamics^21^, or changes in glutamate release following plasticity induction^22^. Existing software packages^15,16,17,18^ possess solutions for drift correction, segmentation, and intensity-time trace analysis, but none were specialized for analysis of iGluSnFR3 recordings, and large segments of the code base were either unnecessary or challenging to customize. We therefore developed a modular approach, enabling end-to-end, high-throughput collection and analysis of hundreds of iGluSnFR3 recordings, through a combination of hardware automation, batch segmentation, and automatic analysis of iGluSnFR3 fluorescence transients. Our scalable, versatile approach enabled deep functional profiling of presynaptic functional heterogeneity (e.g. number of quanta released, *P_r_*, paired-pulse ratio) across hundreds of boutons, with boutons segregating into multiple functional classes according to their iGluSnFR3 responses. We further demonstrated how presynaptic physiology obtained from iGluSnFR3 recordings correlated with expression of Munc13, a protein marker for presynaptic release sites^14^, using post-hoc immunocytochemical analysis of the same synapses. Finally, we extended our approach across the synaptic cleft by combining iGluSnFR3 with a red-shifted, postsynaptically-targeted Ca^2+^ reporter to simultaneously image the activation of postsynaptic AMPA receptors (AMPARs) or NMDA receptors (NMDARs) by single glutamatergic vesicles at single dendritic spines. Our approach allowed pharmacological discrimination of synaptic transmission phenotypes across the dendritic arbor. Directly imaging the flow of information during synaptic transmission at single dendritic spines will enable detailed interrogation of synaptic function, synaptic clustering, and dendritic integration, while providing direct access to the patterns of glutamatergic activity which favor plasticity induction. These innovations lay the groundwork for synapse-by-synapse structure-function analyses to untangle synapse heterogeneity across single neurons and circuits.

## Results

### A high-throughput framework for analyzing glutamate release from single boutons with iGluSnFR3

We sought to leverage iGluSnFR3^19^ to characterize the heterogeneous glutamate release properties of individual presynaptic boutons in cultured rat hippocampal neurons. To accelerate experimental throughput and improve reproducibility, we devised an automated framework for efficiently collecting large datasets (**Figure 1**). We used microscopy software from Andor (Fusion) and their *Python*-accessible REST API to communicate with our microscope (**Figure 1A**). With the REST API, we wrote imaging protocols that captured stimulus-evoked or spontaneous iGluSnFR3 activity along axonal arbors for an arbitrary number of imaging positions using widefield imaging at 200 Hz followed by high-resolution, confocal z-stacks. Custom *Python* scripts allowed flexible, automated control of both electrical stimulation and solution exchange, enabling the design of versatile experiments to probe the physiological properties of presynaptic boutons using iGluSnFR3.

**Figure 1.**
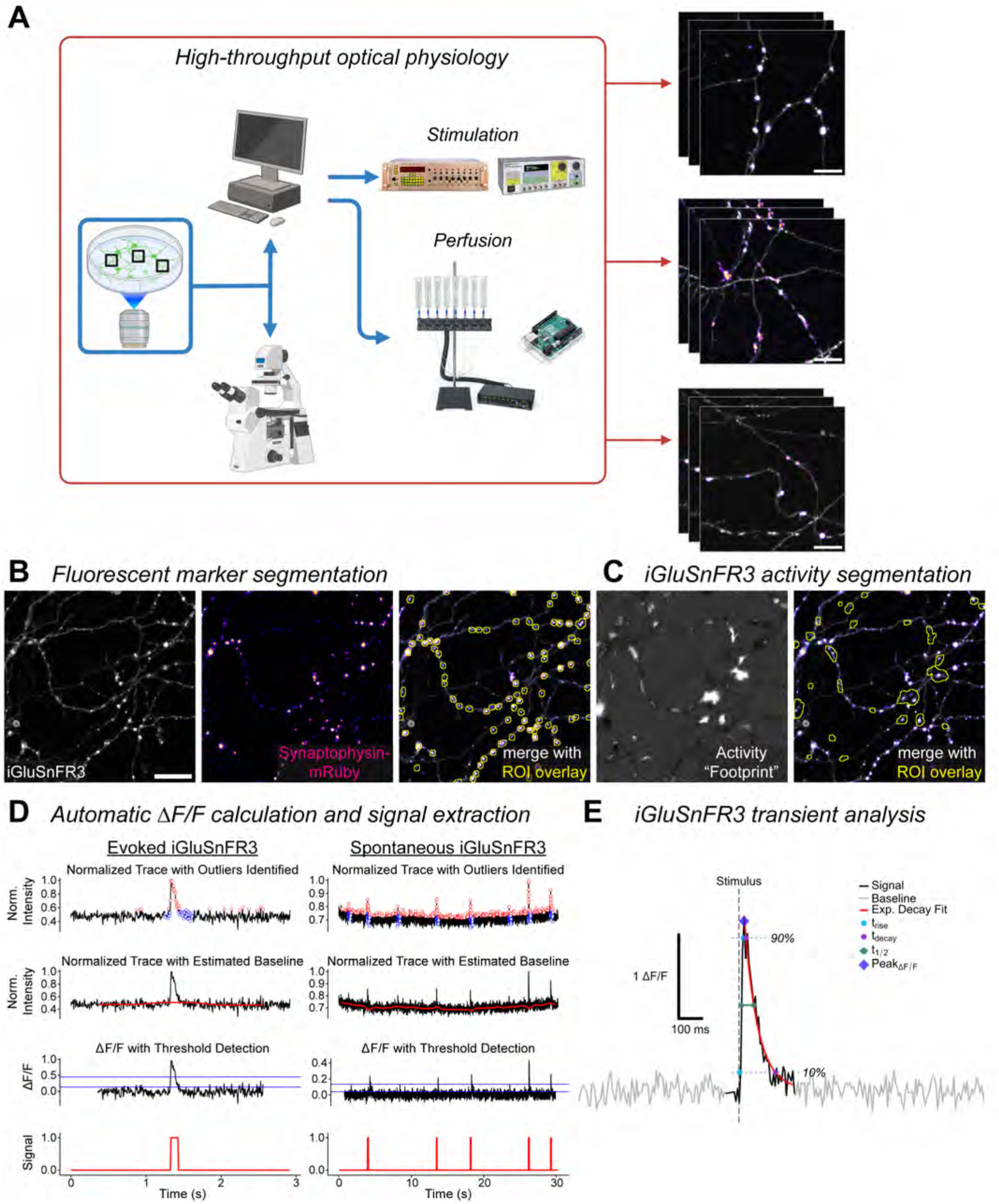
An end-to-end framework enables high-throughput measurement of presynaptic function with iGluSnFR3. **A)** An automated imaging approach using custom Python scripts enables high-throughput optical physiology of presynaptic boutons. Python routines control the microscope, triggering imaging protocols and controlling stage positions via a REST API (Andor, Oxford Instruments). Python sub-routines configure a programmable stimulus generator. Whenever an imaging protocol triggers a TTL pulse to the programmable stimulus generator, the pre-configured stimulus paradigm is sent to an electrical stimulator which stimulates neurons via parallel platinum wire electrodes. Other Python sub-routines toggle pinch valves on a gravity flow perfusion apparatus simultaneously with an Arduino-controlled vacuum to permit fluid exchange. The result is a versatile, automated approach to experimental design that dramatically improves reproducibility and throughput for assays of synapse physiology with iGluSnFR3. At right are imaging regions of axons expressing iGluSnFR3 (gray LUT) and Synaptophysin-mRuby (fire LUT), a marker for the presynaptic bouton. Scale bar = 5 µm. **B,C)** Time-lapse recordings of iGluSnFR3 activity along axons can be batch segmented based on a fluorescent marker (**B**) or iGluSnFR3 activity (**C**). Scale bar = 10 µm. **D)** A custom R algorithm automatically identifies the baseline, calculates ΔF/F, and extracts putative iGluSnFR3 signals. The algorithm’s outputs for two exemplar regions-of-interest (ROIs) are shown. Left, stimulus-evoked iGluSnFR3 activity; right, spontaneous, AP-independent iGluSnFR3 transients. **E)** Extracted iGluSnFR3 transients can be analyzed for several intrinsic peak parameters: peak ΔF/F, exponential decay time constant (τ_decay_), full-width at half-maximum (t_1/2_), 10-90% rise time (t_rise_), 90-10% decay time (t_decay_), and for stimulus-evoked transients, the time interval between peak and stimulus onset (Δt).

To capture iGluSnFR3 activity localized to presynaptic boutons, we transfected hippocampal neurons with iGluSnFR3 and Synaptophysin-mRuby, a fluorescent marker which labels the synaptic vesicle (SV) pool at boutons^23^. We tracked single boutons across many imaging rounds by batch segmenting Synaptophysin-mRuby puncta along iGluSnFR3+ axonal arbors using a custom ImageJ macro and the AI-based ImageJ plugin SynQuant^24^. Alternatively, we could batch segment videos using iGluSnFR3 activity itself with minor modifications of a published approach^25^ (**Figure 1C**). Activity-based segmentation was useful in situations where a synapse marker could not be co-expressed. Where possible, we chose to use a fluorescent marker to track synapses despite its requirement for more image processing, as it allowed analysis of quiescent synapses which could not be identified with activity-based segmentation.

We frequently identified 20 or more ROIs along axonal arbors in single imaging regions and in some experiments, we recorded the same imaging region more than 40 times, resulting in >800 intensity-time traces of iGluSnFR3 activity to be analyzed per region. Thus, with multiple imaging regions in a single experiment, we required a scalable solution for extracting relevant iGluSnFR3 activity for analysis. For this, we developed a custom algorithm in *R* based on a median filter, which automatically corrects baseline fluctuations due to noise and identifies iGluSnFR3 signals caused by stimulus-evoked and spontaneous glutamate release at single boutons (**Figure 1D**, see *Materials and Methods*). **Figure 1E** shows an exemplar stimulus-evoked iGluSnFR3 transient identified by our algorithm, annotated for the intrinsic peak parameters we analyzed: peak amplitude (peak ΔF/F), fitted exponential decay time constant (τ_decay_), the full-width at half-maximum (t_1/2_), 10-90% rise time (t_rise_), 90-10% decay time (t_decay_), and the time-to-peak following electrical stimulation (Δt).

### Batch segmentation methods extract quantal, spontaneous iGluSnFR3 activity localized to boutons

Action potential (AP)-independent, spontaneous glutamate release events should localize to presynaptic boutons marked by Synatophysin-mRuby, providing an intuitive way to benchmark our pipeline. We therefore imaged spontaneous iGluSnFR3 activity at axonal arbors co-expressing iGluSnFR3 and Synaptophysin-mRuby in buffer containing 1 µM tetrodotoxin (TTX) and either 0.5, 1, 2, or 4 mM Ca^2+^ ([Ca^2+^]_bath_), segmenting recordings according to Synaptophysin-mRuby expression or iGluSnFR3 activity. **Figure 2A** shows an exemplar imaging region with many putative boutons. **Figure 2B** shows the boutons (blue) collected by our fluorescent marker segmentation compared to activity-based segmentation (red) for a single imaging trial. Notably, while activity segmentation appeared to merge the responses of multiple boutons in some instances, the strong concordance of activity ROIs with boutons expressing Synaptophysin-mRuby suggests both methods capture spontaneous iGluSnFR3 activity localized to synapses.

**Figure 2.**
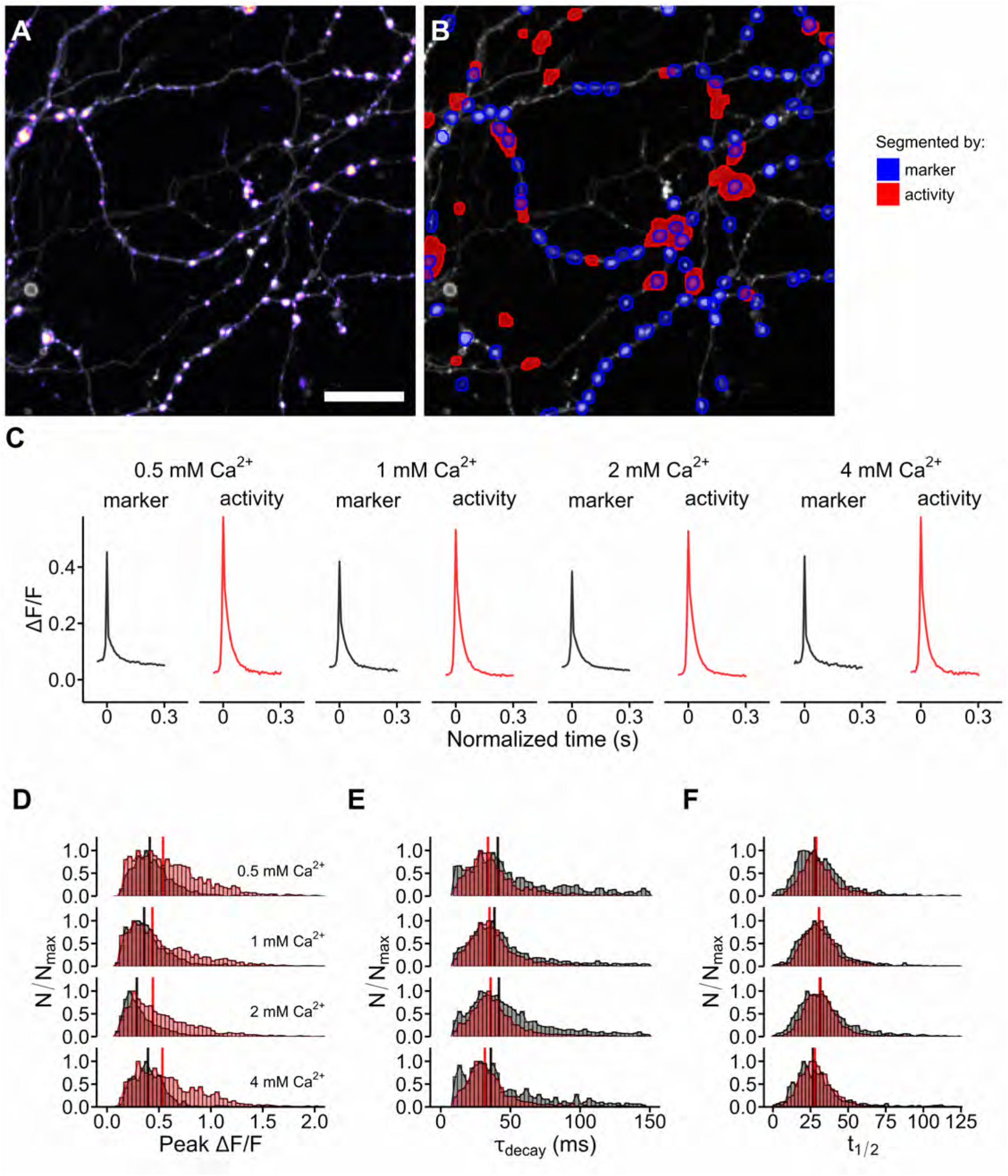
Spontaneous, quantal iGluSnFR3 activity at single boutons is captured by marker- and activity-based segmentation. **A)** An exemplar imaging region (51.2 x 51.2 µm) with an axon co-expressing iGluSnFR3 (grey LUT) and Synaptophysin-mRuby3 (fire LUT). Scale bar = 10 µm. **B)** Spatial map of batch segmented ROIs according to Synaptophysin-mRuby3 expression (blue) or detected iGluSnFR3 activity (red) at [Ca^2+^]_bath_ = 2 mM for one trial. The segmentation comparison indicates that several boutons marked by Synaptophysin-mRuby3 encompass true glutamate release sites, though many more boutons tended to be silent during the 30 s recording period. **C)** The average AP-independent, spontaneous iGluSnFR3 transients at each [Ca^2+^]_bath_ for either segmentation method. For both methods, the shape and amplitude of the average iGluSnFR3 transient was similar across all [Ca^2+^]_bath_, supporting the notion that spontaneous iGluSnFR3 activity represents single vesicle release. **D)** Normalized histogram of peak ΔF/F collected via marker or activity segmentation across [Ca^2+^]_bath_. **E)** Normalized histogram of τ_decay_. **F)** Normalized histogram of t_1/2_. For **D**, **E**, and **F**, vertical lines indicate the medians of the distributions. Detected iGluSnFR3 transients were persistently larger in amplitude with faster τ_decay_ when extracted via activity segmentation.

The average iGluSnFR3 response collected by our high-throughput approach showed little variation in iGluSnFR3 transient shape across [Ca^2+^]_bath_ when compared within segmentation method (**Figure 2C**), reflecting that spontaneous iGluSnFR3 transients are attributable to glutamate release from single SVs at all [Ca^2+^]_bath_. Activity-based segmentation always resulted in a larger average iGluSnFR3 response, likely because activity segmentation captures iGluSnFR3 hotspots directly. By contrast, Synaptophysin-mRuby segmentation captures the centroid of the SV pool at boutons, which will not always perfectly align with the active zone. **Figure 2D-F** show the normalized histograms for selected iGluSnFR3 event parameters across [Ca^2+^]_bath_ and segmentation method. None of the distributions of peak parameters varied substantially in magnitude across [Ca^2+^]_bath_ when compared within segmentation method. Thus, while activity segmentation produces iGluSnFR3 events with larger amplitudes and smaller τ_decay_, Synaptophysin-mRuby segmentation also reliably captures quantal activity at putative single boutons and allowed straightforward tracking of boutons across imaging sessions.

### Estimating the number of synaptic vesicles released during evoked transmission

The number of SVs released following an AP is a critical determinant of synaptic transmission strength^26^. This can be assessed at single synapses using SV-targeted pHluorins to directly count released SVs^13^ and measure the *P_r_* of SV release sites at single boutons following an AP^27,28^. Conversely, the total number of fusion-competent SVs available to be released during an AP, called the readily releasable pool (RRP)^29^, can be measured using mean-variance analysis (MVA)^30^. MVA is typically applied to electrophysiology recordings to calculate the size of the RRP through the relationship *I* = *N_sites_* × *P_r_* × *Q*, where *I* is the mean amplitude of the postsynaptic receptor current, *N_sites_* is the total number of release sites in the RRP, *P_r_* is the uniform release probability for the SVs in the RRP, and *Q* is the current associated with a single SV. However, electrophysiology measures the responses of ensembles of synapses, whereas single-synapse resolution requires fortuitous circuit architecture^31,32^ or advanced experimental paradigms. By contrast, optical sensors which monitor glutamate release via changes in fluorescence intensity (e.g. iGluSnFR3) directly access both the relative magnitude of glutamate release during a single AP and the total size of the RRP through MVA^14^ with single-synapse resolution. Since iGluSnFR3 is sensitive to putative single SV release events, we reasoned that our high throughput approach should be able to determine both how many SVs are released per AP and the overall size of the RRP, identifying the RRP fraction mobilized during each AP at single bouton resolution. To achieve this, we imaged the iGluSnFR3 response evoked by a single AP (10 trials, 20 s between trials) at boutons labeled by Synaptophysin-mRuby (*n*=244 boutons) across different [Ca^2+^]_bath_ (0.5, 1, 2, and 4 mM). **Figure 3A** compares the average evoked iGluSnFR3 response at each [Ca^2+^]_bath_ to the average spontaneous response (spontaneous data corresponds to marker-based segmentation data from **Figure 2**). The amplitude of evoked iGluSnFR3 transients increased substantially with increasing [Ca^2+^]_bath_ as more SVs were released during each AP, in contrast to spontaneous iGluSnFR3 transients (**Figure 3B**). From the normalized histograms of peak ΔF/F and τ_decay_, (**Figure 3C-D**), we observed that the statistics for evoked iGluSnFR3 events at [Ca^2+^]_bath_ = 0.5 mM exhibited substantial overlap with their spontaneous counterparts, suggesting boutons only released a single SV per AP at this [Ca^2+^]_bath_. Most boutons transitioned to multivesicular release as [Ca^2+^]_bath_ increased (**Figure 3C**).

**Figure 3.**
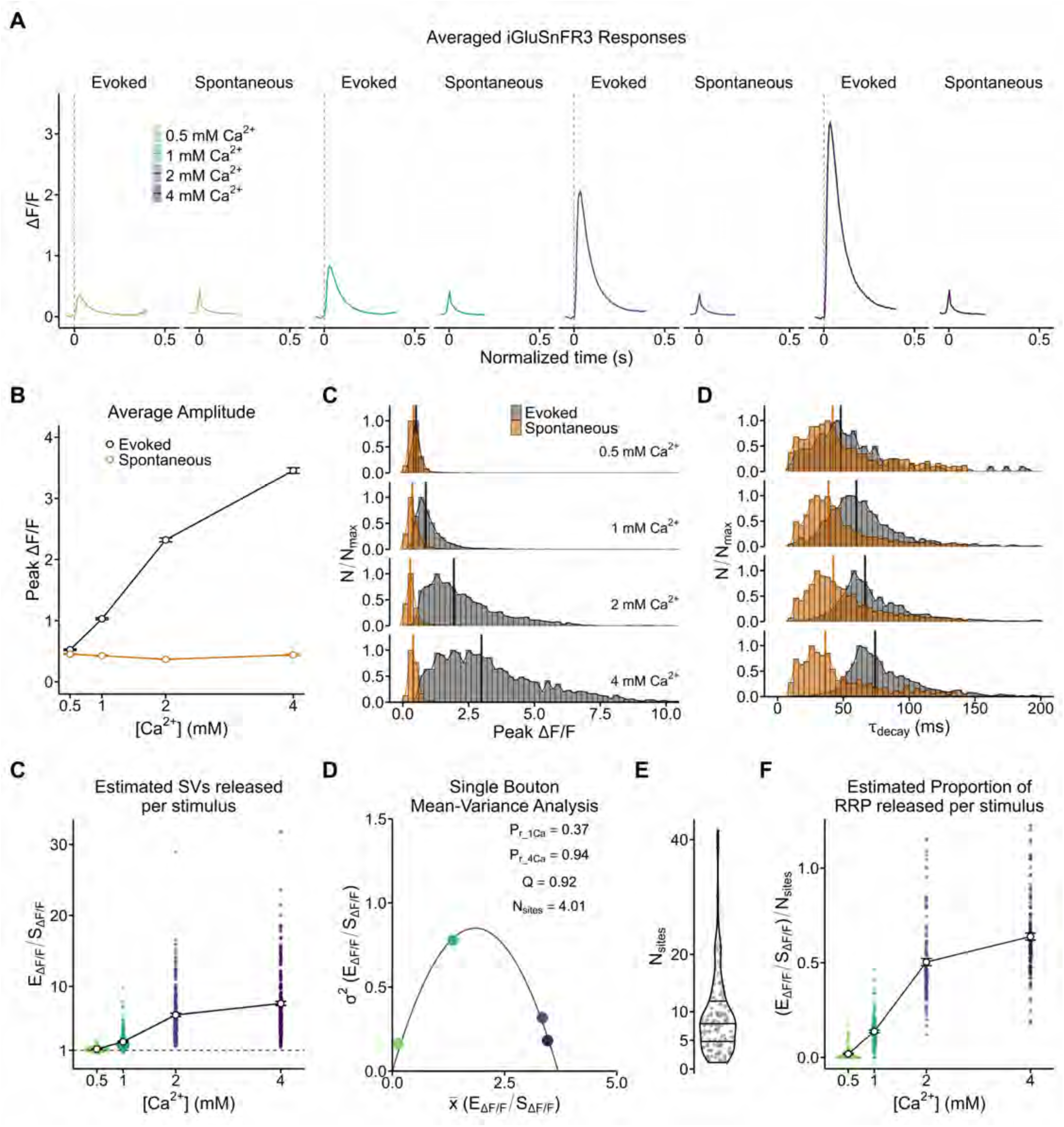
Estimating the number of vesicles released during evoked SV release. **A)** The averaged iGluSnFR3 response for stimulus-evoked and spontaneous glutamate release across [Ca^2+^]_bath_ = 0.5, 1, 2, or 4 mM, incorporating the marker-based segmentation data from Figure 2. Dashed vertical lines indicate stimulus. **B)** Scatterplot of average peak amplitude (peak ΔF/F) vs. [Ca^2+^]_bath_ for stimulus-evoked and spontaneous iGluSnFR3 transients. **C)** Normalized histogram of peak ΔF/F comparing identified spontaneous (orange) with stimulus-evoked (black) iGluSnFR3 transients at each [Ca^2+^]_bath_. At 0.5 mM Ca^2+^, the distribution of spontaneous and stimulus-evoked iGluSnFR3 transients largely overlap, suggesting that at this low [Ca^2+^]_bath_, putative boutons only release a single vesicle in response to stimulus. At 2 and 4 mM Ca^2+^, boutons transition to multivesicular release. **D)** Normalized histogram of τ_decay_. In **C, D,** black and orange vertical lines indicate median values of the distributions for stimulus-evoked and spontaneous iGluSnFR3 transients, respectively. **E)** Scatterplot of the E_ΔF/F_/S_ΔF/F_ vs. [Ca^2+^]_bath_. E_ΔF/F_/S_ΔF/F_ is the peak ΔF/F of stimulus-evoked iGluSnFR3 transients (E_ΔF/F_) normalized to the population average of the peak ΔF/F of spontaneous iGluSnFR3 transients (S_ΔF/F_) at each respective [Ca^2+^]_bath_. Colored dots indicate the mean value of E_ΔF/F_/S_ΔF/F_ at individual boutons. The responses at 0.5 mM Ca^2+^ cluster around Evoked_ΔF/F_/Spont_ΔF/F_ = 1 (horizontal dashed line), indicating univesicular release at this [Ca^2+^]_bath_. **F)** Scatterplot of variance (σ^2^) of E_ΔF/F_/S_ΔF/F_ vs. mean (x̄) E_ΔF/F_/S_ΔF/F_ for an exemplar bouton. Mean-variance distributions were fit with a binomial of the form σ^2^ = *Q*x̄ - x̄^2^/*N_sites_*, where *Q* is the single vesicle iGluSnFR3 response and *N_sites_* is the total number of release sites at each bouton (i.e. the readily releasable pool, RRP). From the fitted binomial, we extracted the uniform release probability at each release site, *P_r_*, for each [Ca^2+^]_bath_. The *P_r_* for 1 and 4 mM Ca^2+^, as well as the solved values for *Q* and *N_sites_* are shown on the plot. **G)** A violin plot of the distribution of solved *N_sites_* per bouton. The behavior of 122/244 boutons in our dataset could be well-described by a binomial fit. **H)** Scatterplot of (E_ΔF/F_/S_ΔF/F_)/*N_sites_* vs. [Ca^2+^]_bath_. As the value of E_ΔF/F_/S_ΔF/F_ was a proxy for the total number of vesicles released per stimulus, the ratio of E_ΔF/F_/S_ΔF/F_ to *N_sites_* permitted us to estimate the proportion of the RRP mobilized per stimulus at each bouton across [Ca^2+^]_bath_.

To count the SVs released per AP, we normalized the peak ΔF/F of each <u>e</u>voked iGluSnFR3 response (E_ΔF/F_) to the population average of the <u>s</u>pontaneous iGluSnFR3 responses (S_ΔF/F_). **Figure 3E** shows the average value of E_ΔF/F_/S_ΔF/F_ vs. [Ca^2+^]_bath_ at single boutons. At 0.5 mM Ca^2+^, the few boutons that were active released approximately 1 SV (E_ΔF/F_/S_ΔF/F_ = 1.12 ± 0.01), while at 4 mM Ca^2+^, boutons released approximately 8.5 vesicles per AP (E_ΔF/F_/S_ΔF/F_ = 8.48 ± 0.32). Ultrastructural observations of boutons in cultured mouse hippocampal neurons indicate that there are 10.1 ± 4.3 (mean ± s.d.) docked SVs per active zone^33^, suggesting that at 4 mM Ca^2+^, most boutons in our sample release >80% of the SVs in the RRP. However, some boutons released > 20 putative SVs per AP at 4 mM Ca^2+^, implying that a subset of boutons in our sample either experienced synaptic crosstalk or possessed multiple active zones.

To better understand what fraction of the RRP was mobilized per AP at individual boutons, we fit the mean (*x̅*) and variance (σ^2^) of the E_ΔF/F_/S_ΔF/F_ values for each bouton with the uniform-probability binomial model^30^, σ^2^ = *Qx̅ - x̅^2^/N_sites_* (**Figure 3F**). Of the 244 boutons in our sample, 122 possessed mean-variance behavior that was well described by the model, yielding an average RRP size of *N_sites_* = 9.05 ± 0.63 (**Figure 3G**), in good agreement with previous ultrastructural measurements and within the range of *N_sites_* (1-10) observed by other glutamate imaging methods^14,20^. Of these 122 boutons, most were quiescent at 0.5 mM Ca^2+^ (E_ΔF/F_/S_ΔF/F_ = 0.101 ± 0.013), but at 1, 2, and 4 mM Ca^2+^ boutons released 1.11 ± 0.08, 3.89 ± 0.22, and 4.98 ± 0.25 SVs, respectively. When we compared estimated SV release (i.e. mean E_ΔF/F_/S_ΔF/F_) to RRP size at each bouton, we found that at 1, 2, and 4 mM Ca^2+^, boutons mobilized 13.7%, 50.4%, and 63.8% of their total RRP, respectively (**Figure 3H**). Taken together, our method extends previous approaches by directly comparing the RRP size calculated from MVA with the approximate number of SVs released per AP at single boutons, permitting a direct assessment of presynaptic efficacy. This approach could provide further access to the time constants associated with vesicle recycling and release site replenishment at individual boutons by measuring how the fraction of RRP released per AP changes with stimulus frequency^29^.

### Tracking the glutamate release behavior of single boutons reveals three functional classes

The high information content of our iGluSnFR3 recordings suggested that boutons could be classified according to their stimulated iGluSnFR3 response behavior. We examined a set of bouton features for each of the 244 boutons in our sample to characterize their iGluSnFR3 behavior in response to a single AP. First, we calculated the probability of observing stimulus-evoked iGluSnFR3 activity (P_iGlu_) for each bouton across [Ca^2+^]_bath_ (**Figure 4A**). At [Ca^2+^]_bath_ = 0.5 mM, most boutons were quiescent (P_iGlu_ = 0.16 ± 0.01), but P_iGlu_ increased sharply at [Ca^2+^]_bath_ = 1 mM (P_iGlu_ = 0.68 ± 0.02). At [Ca^2+^]_bath_ = 4 mM, almost all boutons exhibited iGluSnFR3 activity (P_iGlu_ = 0.96 ± 0.01), suggesting nearly all boutons in our sample were glutamatergic or near enough to a glutamatergic synapse to experience glutamate spillover during periods of strong glutamate release. We also calculated the coefficient of variation (CV) across trials for intrinsic peak parameters, including peak ΔF/F (**Figure 4B**), τ_decay_ (**Figure 4C**), and Δt (**Figure 4D**). For Δt and τ_decay_, the interquartile range (IQR) of the CV distributions decreased as [Ca^2+^]_bath_ increased (**Fig. 4C-D**), indicating that the onset and decay kinetics of evoked iGluSnFR3 responses between trials became more uniform with increasing Ca^2+^ availability for each bouton. The trend for CV of peak ΔF/F (CV_ΔF/F_) was more complex. At 0.5 mM Ca^2+^, the few boutons that were active could only release a single SV following stimulation (**Figure 3F**), biasing CV_ΔF/F_ toward small values. By contrast, at all other [Ca^2+^]_bath_ we observed a broad range of CV_ΔF/F_ values, suggesting the number of SVs released at individual boutons varied substantially between stimuli.

**Figure 4.**
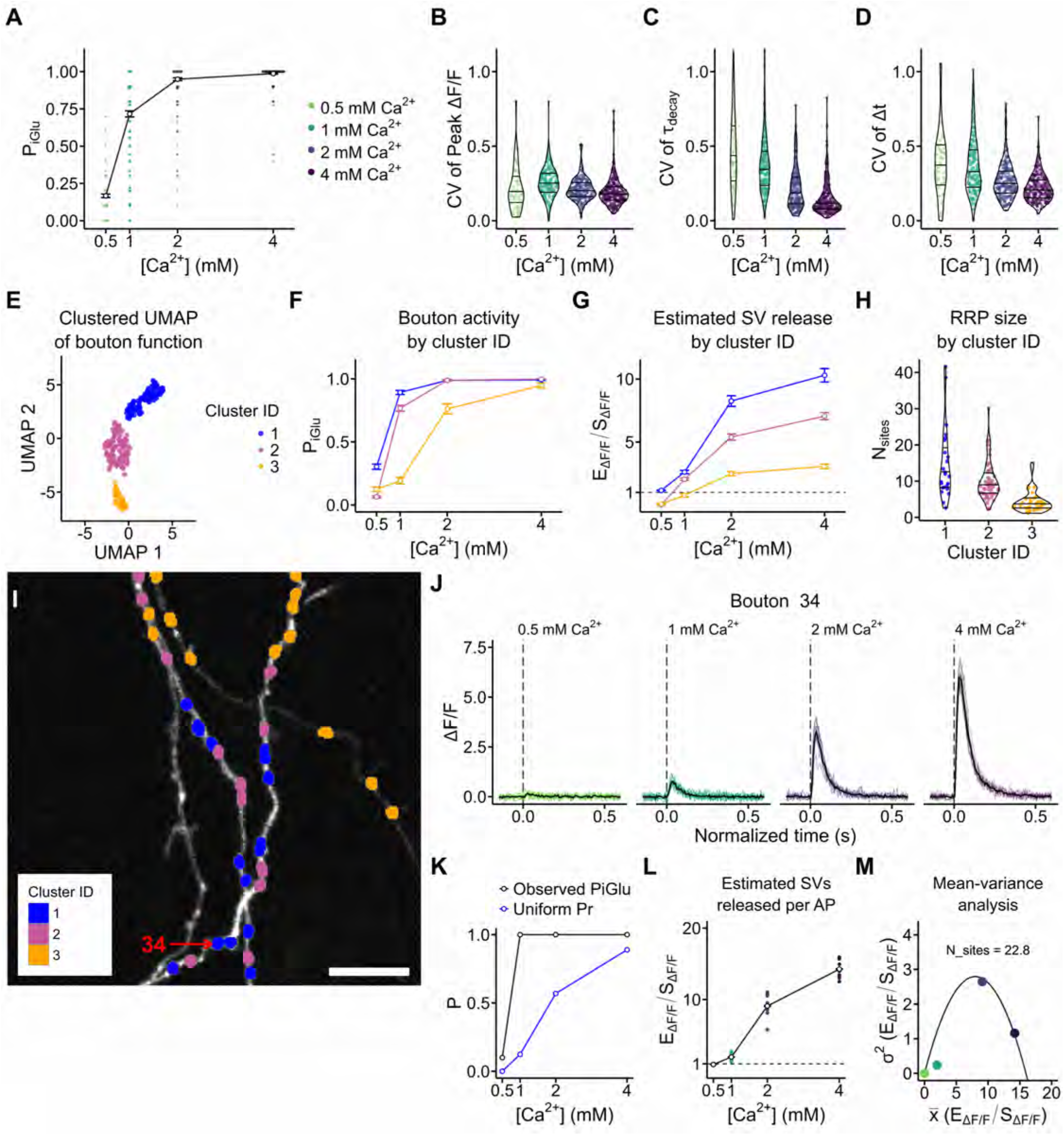
Tracking stimulus-evoked iGluSnFR3 activity at individual boutons captures heterogeneous bouton behavior that can be separated into three functional classes. **A)** The probability of measuring a stimulus-evoked iGluSnFR3 transient (P_iGlu_) was calculated at putative boutons across all [Ca^2+^]_bath_. The trend line indicates the mean ± S.E.M. **B)** The coefficient of variation (CV) of the peak ΔF/F for measured iGluSnFR3 transients. Each colored dot in the violin plots indicates the value of CV_ΔF/F_ for a single bouton. **C)** CV of τ_decay_. **D)** CV of Δt. **E)** UMAP representation of statistics of iGluSnFR3 activity at 244 boutons. DBSCAN of the UMAP indicates that there are three classes of boutons. **F-H)** By grouping the data according to cluster identity, we identified defining features of each bouton grouping. **F)** P_iGlu_ vs. [Ca^2+^]_bath_ grouped by cluster ID. **G)** Estimated number of SVs released (E_ΔF/F_/S_ΔF/F_) vs. [Ca^2+^]_bath_. **H)** Estimated RRP size (*N_sites_*) vs. cluster ID. **I)** An exemplar imaging region with each bouton color coded by cluster ID. Scale bar = 5 µm. **J-M)** Targeted functional analysis of Bouton 34. **J)** Averaged iGluSnFR3 waveform of single AP trials for Bouton 34. The individual trial responses are color-coded by [Ca^2+^]_bath_ with the averaged response in black. **K)** The observed P_iGlu_ (black) and the calculated uniform *P_r_* (blue) vs. [Ca^2+^]_bath_. **L)** The E_ΔF/F_/S_ΔF/F_ curve for Bouton 34 vs. [Ca^2+^]_bath_, with individual trial values color-coded by [Ca^2+^]_bath_. **M)** Scatterplot of variance (σ^2^) of E_ΔF/F_/S_ΔF/F_ vs. mean (x̄) E_ΔF/F_/S_ΔF/F_ for Bouton 34. The colored dots indicate the measured σ^2^ and x̄ at each [Ca^2+^]_bath_, while the black curve indicates the binomial fit.

Between the above statistics and the mean amplitude and variance of the iGluSnFR3 response at each [Ca^2+^]_bath_, we compiled 24 features for each bouton that could be analyzed for their relationship to one another using uniform manifold approximation and projection (UMAP)^34^, which projects multivariate data into two dimensions for cluster analysis. **Figure 4E** shows the UMAP representation of the 244 boutons, which clustered into three different classes of boutons. **Figure 4F** shows the trends in P_iGlu_ according to cluster identity (cluster ID), revealing that Class 1 boutons were almost always active at [Ca^2+^]_bath_ = 1 mM (P_iGlu_ = 0.89 ± 0.02), while Class 3 boutons were only active once every 5 stimuli (P_iGlu_ = 0.19 ± 0.03). When SVs were in their lowest *P_r_* state at 0.5 mM Ca^2+^, active Class 1 boutons reliably released a single SV (E_ΔF/F_/S_ΔF/F_ = 1.16 ± 0.04) while Class 2 and 3 boutons were categorically quiescent (**Figure 4G**). When SVs were in their highest *P_r_* state at 4 mM Ca^2+^, Class 1 boutons released ∼10 SVs, >3-fold more than Class 3 boutons (Class 1: E_ΔF/F_/S_ΔF/F_ = 10.3 ± 0.5, Class 3: E_ΔF/F_/S_ΔF/F_ = 3.1 ± 0.2).

To examine the basis of the observed functional classification, we mapped the cluster IDs onto the bouton parameters extracted from MVA. The distribution of *N_sites_* according to cluster ID suggested that the fidelity and magnitude with which Class 1 boutons responded to stimuli could be explained by their larger RRPs (Class 1: *N_sites_* = 14.7 ± 2.0, **Figure 4H**). These large RRPs imply Class 1 boutons possess multiple active zones, but we also note these boutons possessed faster iGluSnFR3 kinetics (smaller Δt, t_rise_, τ_decay_) with lower variance in their response on all metrics relative to Class 2 and 3 boutons (**Figure S1**). Taken together, these data may imply that Class 1 boutons possess more mature active zones than their Class 3 counterparts.

A key advantage of optical physiology is that functional heterogeneity is spatially resolved. To visualize how bouton physiology varies along the axonal arbor, we mapped cluster IDs to their cellular position (**Figure 4I**). To aid visual comparison across boutons, we generated “Bouton Reports” for each bouton that summarize key features. **Figure 4J-M** shows an example Bouton Report, including the iGluSnFR3 response for all 40 single-stimulus trials (**Figure 4J**), a comparison of P_iGlu_ with the observed uniform *P_r_* from MVA (**Figure 4K**), the estimated SVs released per AP for each trial (**Figure 4L**), and the binomial fit of the mean-variance curve (**Figure 4M**). These tools constitute a data-driven approach to examine single-synapse physiology.

### Pairing immunocytochemistry with high-throughput optical physiology captures developmental changes in presynaptic function correlated with Munc13 expression

We speculated that our cluster analysis had revealed bouton states along a developmental trajectory, in which Class 3 boutons represented nascent presynapses with fewer functional release sites, while Class 1 boutons represented more mature presynapses. However, we could not exclude that some boutons may be “orphans” lacking a postsynaptic partner, and the lack of protein expression data at these synapses meant we could only speculate as to the molecular basis of the observed functional classes (**Figure 4E**). To address these limitations, we designed an experiment to measure both RRP size and the expression of a key presynaptic protein, Munc13-1 (hereafter, Munc13), at excitatory synapses across two neuron culture ages. Munc13 marks presynaptic release sites and its expression scales with RRP size^14,35^, suggesting our experimental design would simultaneously give direct access to how synaptic functional properties fluctuate with developmental age while providing orthogonal validation of our MVA approach. We co-expressed iGluSnFR3 and PSD95.FingR.TagRFP^36^ in hippocampal neurons, allowing segmentation of bona fide glutamatergic synapses, then measured single-synapse RRP size at either 14 (DIV14) or 21 days in vitro (DIV21). Following each live-cell imaging session, neurons were immediately fixed and stained for Munc13. Through a combination of image post-processing and spatial analysis, we were then able to reliably link Munc13 content of synapses to their functional recordings for 148 synapses at DIV14 and 225 synapses at DIV21. **Figure 5A** shows the spatial context of a single-synapse, averaged iGluSnFR3 responses at all [Ca^2+^]_bath_ with corresponding mean-variance analysis, and Munc13 abundance relative to other synapses imaged at either DIV14 or DIV21 (for other examples, see **Figure S3**). We observed similar synapse sampling density per imaging field for each age cohort (**Figure 5B**).

**Figure 5.**
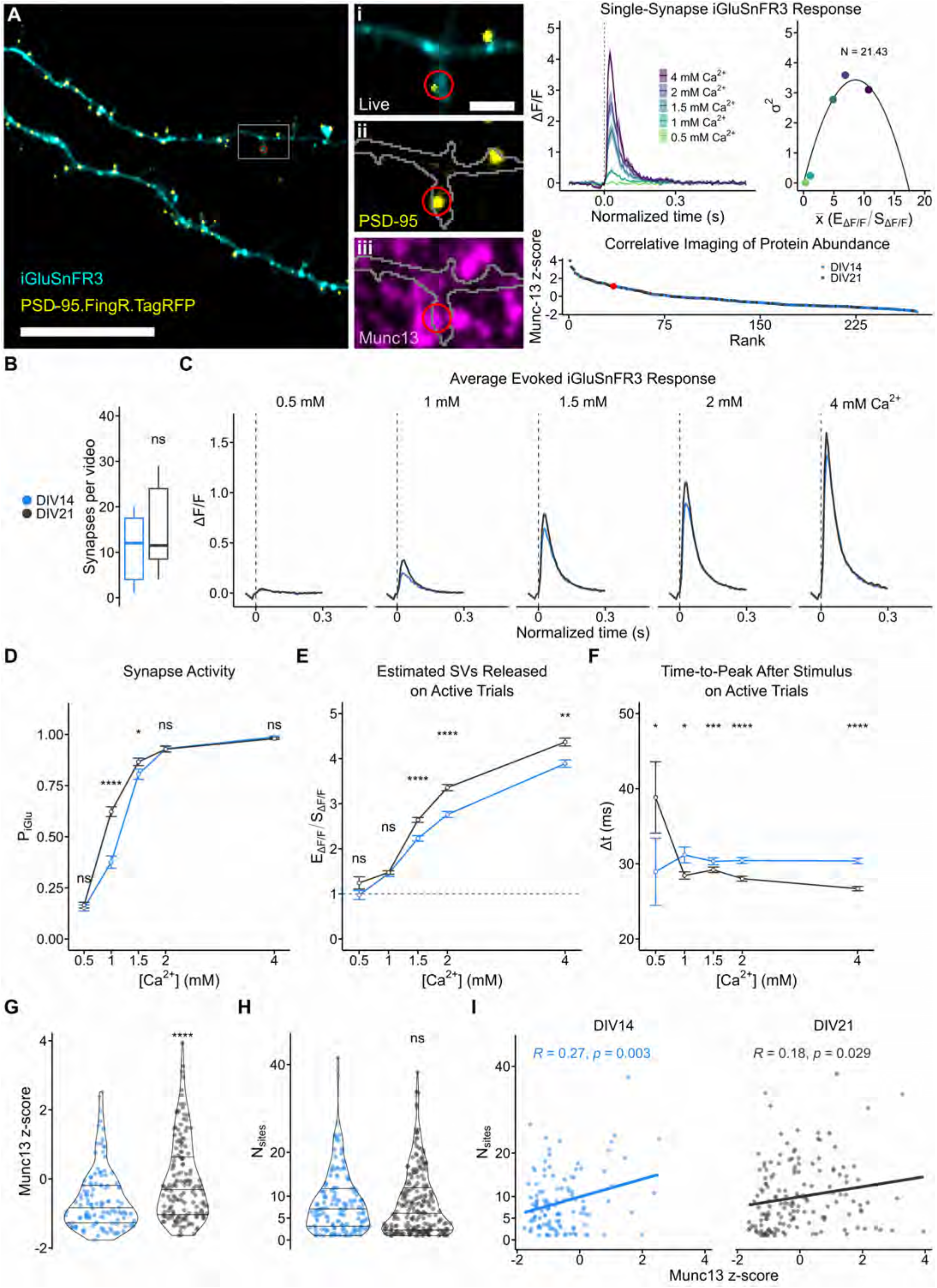
Age-dependent enhancements in the magnitude, kinetics, and reliability of evoked iGluSnFR3 responses correlate with increased Munc-13 expression at single synapses. **A)** An example imaging region of a DIV21 cultured hippocampal neuron dendrite, co-expressing iGluSnFR3 (cyan) and PSD-95.FingR.TagRFP (yellow). Scale bar = 20 µm. **(i)** Inset of the highlighted region in **A**. Scale bar = 2 µm. A synapse-of-interest is highlighted in red. **(ii)** Post-hoc imaging of the same region following immunocytochemistry. PSD-95.FingR.TagRFP signal was imaged at higher laser intensity and exposure time, permitting better resolution of postsynaptic scaffold area. A mask of the live iGluSnFR3 signal is shown in grey. **(iii)** Post-hoc imaging of the same region showing the ICC signal for Munc-13. **Right**, in the order they appear: the Single-Synapse iGluSnFR3 response to a single AP in different [Ca^2+^]_bath_ and the corresponding mean-variance analysis, estimating 21.43 release sites in the RRP. **Bottom**, rank-order plot of the normalized Munc-13 signal (z-score) for each synapse in the dataset, collected from DIV14 and DIV21 neurons. The synapse-of-interest’s position in the rank-order plot is highlighted in red. **B)** The synapses per video which possessed both mean-variance analysis and correlated Munc-13 content. There was no significant difference in synapse sampling density per video between DIV14 (n=148 synapses) and DIV21 (n=225 synapses). **C)** Averaged iGluSnFR3 response to a single AP for DIV14 and DIV21 synapses. **D)** Synapse activity probability (P_iGlu_) for DIV14 and DIV21 synapses. DIV14 synapses were more reluctant to release glutamate at lower [Ca^2+^]_bath_. **E)** Estimated SVs released per AP (E_ΔF/F_/S_ΔF/F_) for DIV14 and DIV21 synapses. **F)** Time-to-Peak (Δt, ms) following stimulus for DIV14 and DIV21 synapses. DIV14 synapses exhibited slower iGluSnFR3 kinetics at all [Ca^2+^]_bath_, except for 0.5 mM, where sampling density was very low due to low release probability. **G)** Normalized Munc-13 expression as measured by immunocytochemistry (z-score). DIV21 synapses accumulated significantly more Munc-13 than DIV14 synapses. **H)** N_sites_ for DIV14 and DIV21 synapses as determined by mean-variance analysis. **I)** The correlation between N_sites_ measured by mean-variance analysis and Munc-13 expression and corresponding linear fit of the data. R and p values from the Pearson correlation shows that in DIV14 synapses, Munc-13 expression more strongly predicts the value of N_sites_. Solid line indicates a linear fit of the data to guide the eye.

**Figure 5C** shows the averaged waveforms of single AP-evoked iGluSnFR3 responses at each [Ca^2+^]_bath_ we tested. Synapses consistently released more glutamate at DIV21 than at DIV14, though the effect was subtle (**Figure 5E**, DIV14, 3.89 ± 0.08 SVs; DIV21, 4.37 ± 0.09 SVs at 4 mM Ca^2+^). P_iGlu_ was significantly smaller at 1 and 1.5 mM Ca^2+^ for DIV14 synapses, suggesting basal release probability increases as synapses develop (**Figure 5D**). DIV21 synapses possessed faster iGluSnFR3 kinetics, with faster t_1/2_, t_rise_, t_decay_, and Δt at all [Ca^2+^]_bath_ (**Figure 5F**, **Figure S4**). Our sampling frequency inherently limited the possible values of Δt, but even with this caveat, DIV21 synapses exhibited a clear bias toward faster Δt values as [Ca^2+^]_bath_ increased, while the effect was more modest for DIV14 synapses (**Figure S4D**).

As expected, Munc13 content was greater in DIV21 synapses than DIV14 synapses, suggesting that either release sites continued to be added as synapses develop or the Munc13 content per release site increased (**Figure 5G**). Consistent with the latter, RRP size did not differ significantly between age cohorts per MVA (**Figure 5H**, DIV14: 8.68 ± 0.58; DIV21: 8.54 ± 0.55). Despite this, RRP size robustly correlated with Munc13 content for both age cohorts, providing molecular validation of our MVA approach (**Figure 5I**). Further, SVs released per AP, variance in SVs released per AP, and P_iGlu_ all correlated strongly with synaptic Munc13 content in the DIV14 cohort, with most correlations becoming statistically insignificant in the DIV21 cohort (**Figure S5**). Together, these data suggest a model in which Munc13 accumulation at synapses increases the number of release sites in the RRP early in development, while at later time points Munc13 content per release site is increased to affect the kinetics of release.

### Single synapses exhibit broad variability in short-term plasticity that exhibits dependence on Munc13 expression

While basal release probability and RRP size constitute the fundamental determinants of presynaptic strength, the presynaptic terminal also contributes to synaptic computations via short-term plasticity, in which pairs of stimuli delivered in short succession can drive increased (facilitation) or decreased (depression) neurotransmitter release on the second pulse^4^. Given the flexibility of our automated imaging approach, we next used it to resolve diversity in short-term plasticity dynamics at single synapses. For this, we recorded from neurons co-expressing iGluSnFR3 and PSD95.FingR.TagRFP, measuring iGluSnFR3 responses to a battery of stimuli, including single APs (test pulse) and paired stimuli separated by a broad range of interstimulus intervals (ISIs) ranging from 60 ms to 500 ms. Neurons were subsequently fixed and stained for Munc13 content, allowing us to measure how short-term plasticity dynamics depend on the accumulation of release sites at individual excitatory synapses.

**Figure 6A** shows the averaged iGluSnFR3 responses for each protocol and [Ca^2+^]_bath_ (1 or 2 mM). The kinetics of iGluSnFR3 result in a convolution of the responses to the 1^st^ and 2^nd^ stimuli at ISIs ≤ 150 ms, obscuring the true amplitude of the 2^nd^ stimulus response. To circumvent this, we subtracted the average response to a single stimulus at each bouton from the responses for each paired pulse trial, allowing measurement of the 2^nd^ stimulus response at all ISIs (**Figure S6B**), and subsequently, calculation of the paired-pulse ratio (PPR), which summarizes the short-term plasticity dynamics for each synapse across all ISIs. **Figure 6B** shows the average measured PPRs at all ISIs and [Ca^2+^]_bath_. Consistent with expectations, the elevated initial release probability at 2 mM Ca^2+^ drove most synapses toward depression or neutrality at all ISIs (**Figure 6B, Figure S6C**). By contrast, at [Ca^2+^]_bath_ = 1 mM we observed that synapses could facilitate, depress, or exhibit no change in their response at the second stimulus for all ISIs, and that these responses appeared to favor specific ISIs ≤ 100 ms (**Figure S7**). Indeed, in separate experiments at single boutons, we also observed that synapses facilitated only at specific ISIs (**Figure S6**). At a minimum, the broad heterogeneity in synaptic PPRs we observed across individual dendrites (**Figure 6C**) implies vesicle trafficking dynamics differ between synapses, though this may be less regulated in cultured hippocampal neurons than *in vivo*.

**Figure 6.**
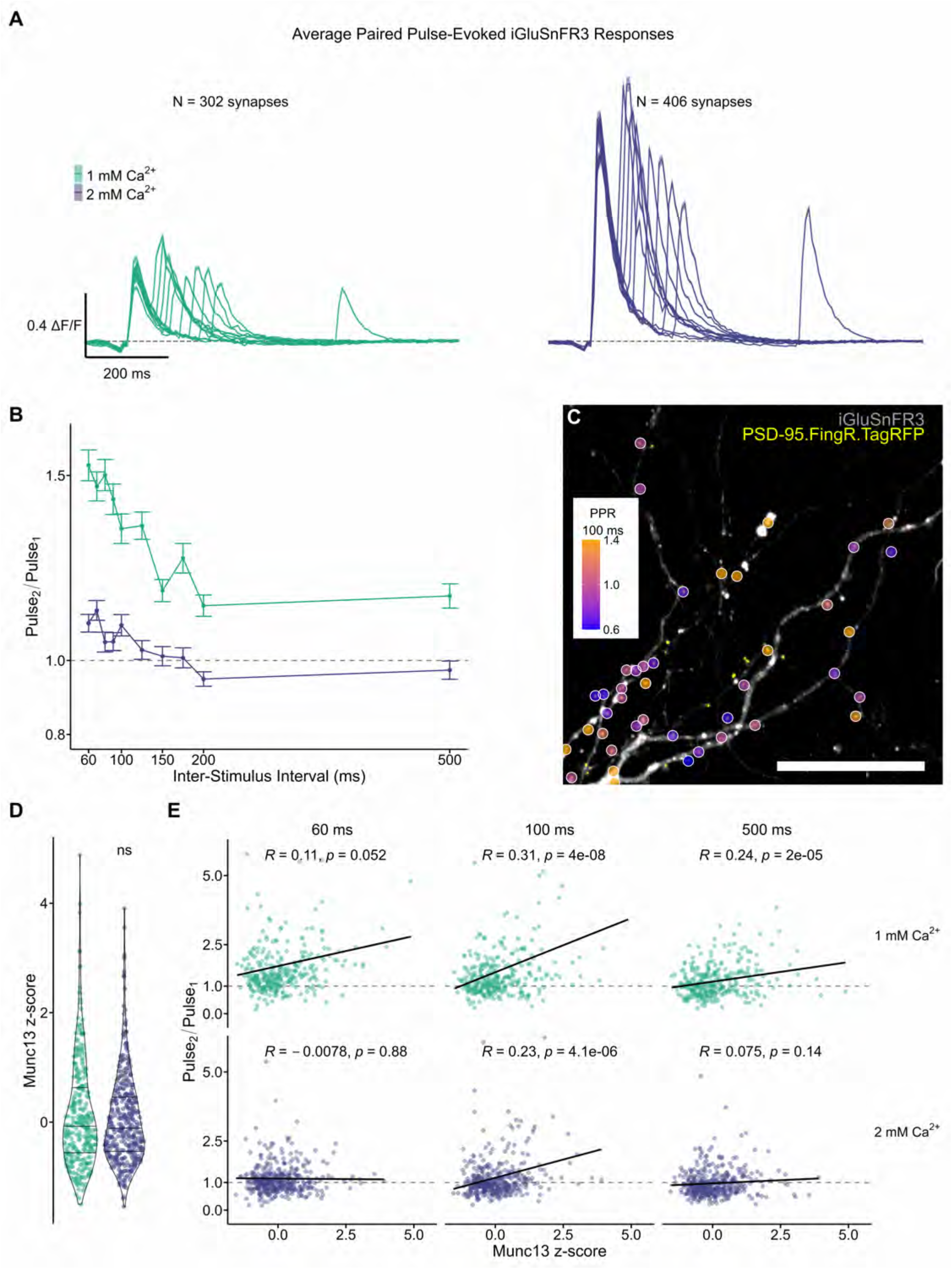
Munc-13 content at individual excitatory synapses predicts presynaptic short-term plasticity behavior at physiological Ca^2+^. **A)** Average iGluSnFR3 response to paired-pulse stimulus across a range of interstimulus intervals (ISIs) and two different [Ca^2+^]_bath_ conditions. Listed are the number of excitatory synapses collected for each condition with correlated post-hoc immunocytochemistry (ICC) measurements of Munc-13. Note that 0.4 ΔF/F corresponds to the average iGluSnFR3 response to a spontaneously released single SV. **B)** Paired-pulse ratio (PPR) across ISIs and [Ca^2+^]_bath_ conditions. Synapses largely depress or remain neutral at [Ca^2+^]_bath_ = 2 mM. **C)** Spatial organization of PPRs at 100 ms across a single dendrite from the dataset at [Ca^2+^]_bath_ = 1 mM. Synapses exhibit a wide range of short-term plasticity behavior. Scale bar = 20 µm. **D)** Violin plot of normalized Munc-13 expression (z-score) from ICC. At the level of Munc-13 expression, synapses sampled at each [Ca^2+^]_bath_ were not statistically different from one another, permitting direct comparisons of how Munc-13 expression interacts with synaptic physiology across [Ca^2+^]_bath_. **E)** Scatter plots showing the correlation of PPR with Munc-13 expression for selected ISIs (see *Supporting Information* for all ISIs). Each point is a synapse, with the black lines showing the linear fit of the distribution to guide the eye. R and p values correspond to the strength and statistical significance of the Pearson correlation, respectively. At 1 mM Ca^2+^, PPRs positively correlate with Munc-13 expression, while at 2 mM Ca^2+^, PPR appears unrelated to Munc-13 expression except at select ISIs.

To better understand the molecular underpinnings of these diverse dynamics, we measured the correlation between PPR and Munc13 expression across our two [Ca^2+^]_bath_. Importantly, there was no statistical difference in Munc13 expression between the two populations of synapses (**Figure 6D**), permitting direct comparisons of how extracellular Ca^2+^ and Munc13 expression interact to produce short-term facilitation and depression. Increasing Munc13 expression correlated with increasing PPR (i.e. biased towards facilitation) across all ISIs at 1 mM Ca^2+^, whereas PPR only correlated with Munc13 expression for specific ISIs at 2 mM Ca^2+^ (**Figure 6E**, **Figure S8**). Consistent with our observations at DIV21 synapses in **Figure 5**, Munc13 expression did not correlate with the amplitude (ΔF/F) of either the 1^st^ or 2^nd^ pulse (**Figure S8**). By contrast, the amplitude (ΔF/F) of both the 1^st^ and 2^nd^ pulse, but not PPRs, correlated with PSD95.FingR.TagRFP expression (**Figure S9**). Thus, while basal glutamate release was higher at synapses expressing more postsynaptic scaffolds, synaptic facilitation only correlated with Munc13 expression.

### Simultaneous imaging of iGluSnFR3 and a far-red Ca^2+^ reporter enables direct, all-optical measurement of NMDAR-mediated synaptic transmission at single dendritic spines

In postsynaptic whole-cell patch-clamp recordings, measures of presynaptic function are indirect and postsynaptic measures lack spatial specificity. Dual optical assays of pre- and postsynaptic function will therefore be required to untangle the basic functional properties of single synapses. Previous work introduced spine-jRGECO1a^37^, a fusion construct that expresses jRGECO1a in actin-rich compartments of the cell like the dendritic spine. However, 488 nm light causes jRGECO1a to photoswitch^38^, dramatically reducing SNR and rendering spine-jRGECO1a incompatible with iGluSnFR3 in imaging paradigms. To avoid this, we replaced the coding sequence for jRGECO1a with that of HaloTag^39^, which we labeled with a far-red chemigenetic Ca^2+^ sensor, JF_646_-BAPTA-HaloTagLigand-AM^40^ (hereafter, JF_646_-BAPTA). Co-expression of iGluSnFR3 with spine-HaloTag thus enabled simultaneous access to glutamate and Ca^2+^ dynamics at dendritic spines (**Figure 7A, B**). We initially focused on resolving NMDAR activation by spontaneously released glutamate, imaging hippocampal neurons co-expressing iGluSnFR3 and spine-HaloTag in buffer with 4 mM Ca^2+^, 0 mM Mg^2+^, and 1 µM TTX^41^. These recordings were segmented via iGluSnFR3 activity hotspots (**Figure S10**), allowing observation of iGluSnFR3 transients localized to dendritic spines that were often, but not always, followed by a JF_646_-BAPTA transient (**Figure 7B**). We used the iGluSnFR3 peaks detected by our algorithm to guide identification of correlated JF_646_-BAPTA peaks (shaded regions **Figure 7B**, bottom), and considered these event pairs to be putative synaptic transmission events if the JF_646_-BAPTA amplitude exceeded a threshold of 5 times the standard deviation of the JF_646_-BAPTA baseline (**Figure 7B, C**).

**Figure 7.**
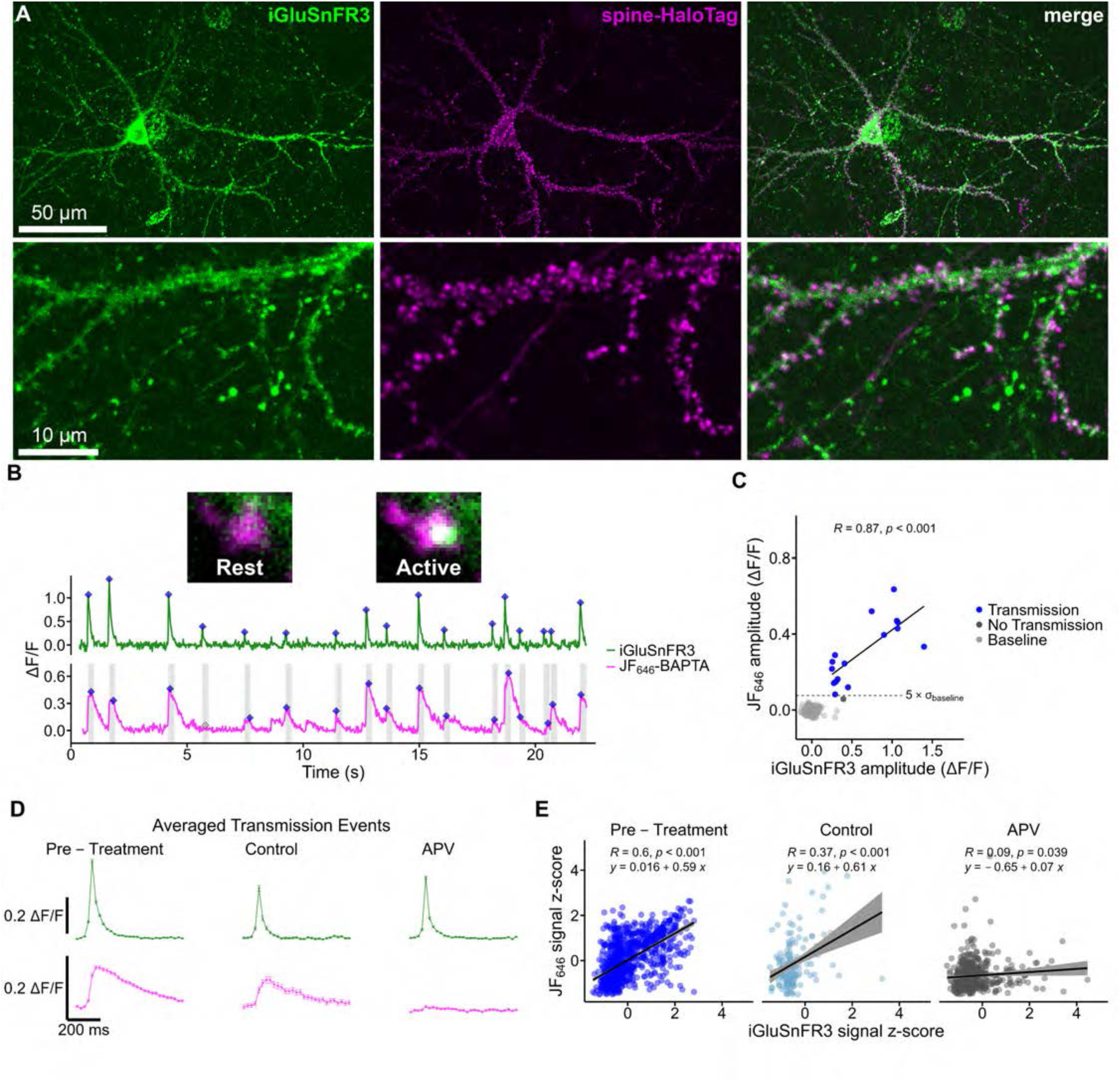
Directly imaging NMDAR activation by spontaneous glutamate release at single dendritic spines using a dual fluorescent reporter approach. **A)** An exemplar imaging region with a neuron expressing both iGluSnFR3 and spine-HaloTag dyed with JF_646_-BAPTA-HTL-AM. spine-HaloTag accumulates in actin-rich compartments of the cell, like the dendritic spine head. Insets show a zoom-in of a stretch of dendrite. **B)** ΔF/F vs. time trace with spontaneous activity collected from iGluSnFR3 (green) and JF_646_-BAPTA-AM (magenta) from a single dendritic spine head. In the green trace, our peak finding algorithm identified iGluSnFR3 signals and their peak maxima (blue diamonds). In the magenta trace, grey shaded regions in the magenta trace indicate regions around iGluSnFR3 signals. To identify the peaks of JF_646_-BAPTA signals, we searched for local maxima in these grey shaded regions (blue diamonds). Insets show the corresponding dendritic spine at rest and during synaptic transmission. **C)** Scatterplot of JF_646_ peak amplitude vs. the iGluSnFR3 peak amplitude for the spine in **B**. Our peak finding algorithm identified putative baseline indices as part of its baseline adjustment routine. For each dendritic spine, we calculated the standard deviation (σ) of the baseline noise. We set a threshold of 5σ, and JF_646_-BAPTA signals above this threshold were considered putative synaptic transmission events (blue dots). **D)** We collected all transmission events from spines before and after exposure to a control solution or a solution containing 100 µM APV. The averaged waveforms show that APV treatment abolished JF_646_-BAPTA signals, but not iGluSnFR3 signals, indicating that JF_646_-BAPTA signals represent NMDAR-mediated Ca^2+^ flux. **E)** We normalized transmission events by spine (z-score) and plotted the population correlations between iGluSnFR3 and JF_646_-BAPTA signals. The slope of the correlation is largely intact following treatment of spines with a control solution, while treatment with 100 µM APV abolished the correlation. The Pearson correlation coefficient (R) and statistical significance are indicated on each subplot.

To determine whether these JF_646_-BAPTA signals represent NMDAR activation by glutamate, we imaged neurons before and after perfusion of an NMDAR antagonist APV or a control solution. Indeed, the time-aligned, averaged transmission events indicated that APV treatment abolished JF_646_-BAPTA events but not iGluSnFR3 events, indicating that this Ca^2+^ activity originated from NMDAR activation (**Figure 7D**). To confirm these observations quantitatively, we normalized event amplitudes for spine-to-spine variation by z-scoring. APV abolished the strong positive correlation between normalized JF_646_-BAPTA and iGluSnFR3 signals seen before and after perfusion with a control solution (**Figure 7E**).

Our dual fluorescence reporter approach clearly captures information transfer from pre- to postsynaptic compartments. However, consistent experimental success was limited by co-expression challenges with the two fluorescent sensors. Even when sensor co-expression was successful, GPI-anchored iGluSnFR3^19^ was often outside the spine head, resulting in scenarios where segmentation of iGluSnFR3 activity often captured multiple synapses or regions of membrane which did not feature correlated Ca^2+^ signaling dynamics (**Figure S10A**). To address these challenges, we developed a bicistronic construct which expressed spine-HaloTag and iGluSnFR3 linked to a modified cytosolic fragment of Stargazin (SGZ)^19^ from a single plasmid. Transfection of hippocampal neurons with spine-HaloTag-P2A-iGluSnFR3.SGZ dramatically improved sensor co-expression and iGluSnFR3 localization at synapses, permitting reliable segmentation of correlated glutamate and Ca^2+^ activity at single spines (**Figure S10B**).

### Direct synaptic transmission imaging captures diverse Ca^2+^ signaling phenotypes at single spines with single-vesicle resolution

Elevation of postsynaptic intracellular Ca^2+^ triggered by transmission at excitatory synapses is critical in numerous aspects of neuronal signaling^42^. Activation of NMDARs underlies induction of synapse-specific plasticity in principal neurons, but AMPARs lacking the GluA2 subunit are also Ca^2+^-permeable and all AMPARs can initiate Ca^2+^ rises through a number of means including activation of voltage-dependent Ca^2+^ channels or subsequent calcium-induced Ca^2+^ release. The receptor source of Ca^2+^ is critical for the nature of plasticity^43,44^, but there is little information regarding how spontaneous transmission may drive downstream calcium signaling or whether this varies substantially between synapses. Given that our technique readily resolves glutamate-dependent Ca^2+^ elevation, we reasoned that we should be able to compare NMDAR- or AMPAR-driven Ca^2+^ increases across synapse populations through sequential pharmacological antagonism of these receptors. **Figure 8A** illustrates the experiment via the response from an example synapse: First, AMPAR-mediated spontaneous synaptic transmission can be seen while AMPARs are open but NMDARs are closed (blue trace, +APV/-DNQX); Next, NMDARs are unblocked and AMPARs are blocked as APV is washed out and DNQX is washed in (red trace, -APV/+DNQX); Finally, both receptor populations are blocked as a negative control (+APV/+DNQX). In this manner, we collected AMPAR-mediated and NMDAR-mediated activity driven by single glutamatergic SVs for 500 spines from 19 dendrites.

**Figure 8.**
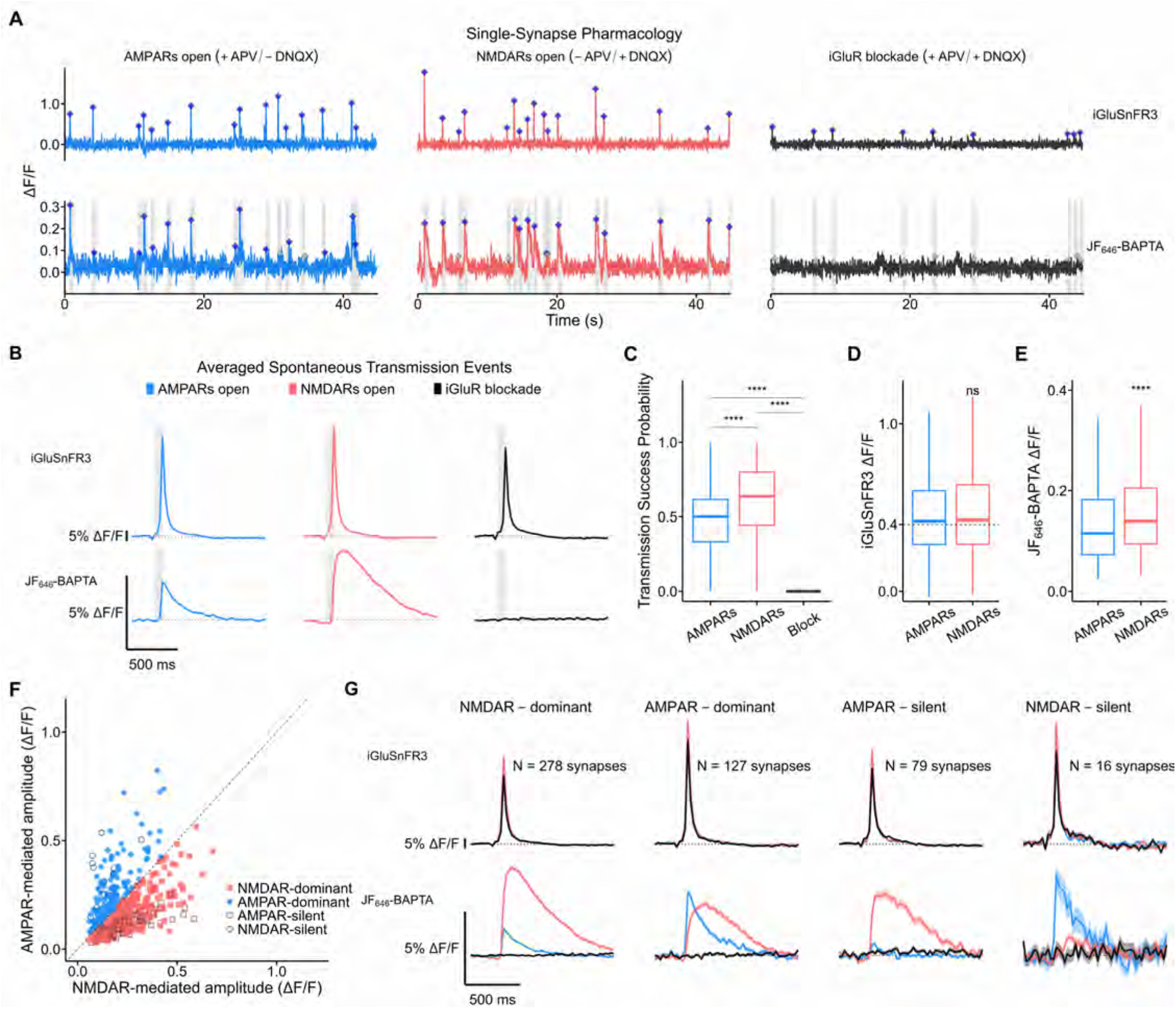
Synaptic transmission imaging enables pharmacological discrimination of synapse types according to iGluR content. **A)** Single-synapse pharmacology using iGluSnFR3 and JF_646_-BAPTA to report on pre- and postsynaptic glutamate and Ca^2+^ dynamics, respectively. Neurons were sequentially bathed in pharmacological blockers to isolate postsynaptic Ca^2+^ responses to spontaneous single-vesicle driven by AMPARs (blue, +APV/-DNQX), NMDARs (red, -APV/+DNQX), or the lack of response during iGluR blockade (black, +APV/+DNQX). Shown is the recording at each phase of the experiment for a single-synapse. Gray shading on the traces indicates the time interval that was scanned for a local maximum in the JF_646_-BAPTA trace. Blue hashes indicate either an identified iGluSnFR3 event (top) or a successful transmission event (bottom). Transmission failures are indicated by grey hashes (see iGluR blockade trace). **B)** Averaged spontaneous synaptic transmission events. Top row, iGluSnFR3 responses, bottom row, correlated Ca^2+^ responses during “AMPARs open” (+APV/-DNQX), “NMDARs open” (-APV/+DNQX) or “iGluR blockade” (+APV/+DNQX). Note that while confidence intervals are present, they are narrow due to the large sample size. Data are averaged from 16,372 spontaneous iGluSnFR3 events collected from 500 spines from 19 dendrites. The gray shaded region of each trace indicates the time interval from the start of the iGluSnFR3 transient to its peak to guide the eye to the kinetic differences in the averaged JF_646_-BAPTA transients between the AMPAR and NMDAR conditions. Note that the trace averages are limited by the sampling frequency of our imaging paradigm (∼40 Hz). Dashed line indicates ΔF/F = 0 to guide the eye. **C)** Observed probability of successful transmission. NMDAR-mediated transients were more consistently activated by single vesicle release events than AMPAR-mediated transients, consistent with the higher affinity of NMDARs for glutamate. iGluR blockade resulted in 99.7% of synaptic transmission attempts (∼4455 events from 500 synapses) being categorized as failures. **D)** Amplitude of iGluSnFR3 signals during AMPAR and NMDAR imaging phases among successful synaptic transmission events. Dashed line indicates the average spontaneous iGluSnFR3 event amplitude collected from presynaptic boutons (see Figure 3), emphasizing the stability of single vesicle iGluSnFR3 amplitudes whether measured pre- or postsynaptically. **E)** Amplitude of JF_646_-BAPTA signals during AMPAR and NMDAR imaging phases. **F)** Relationship between the maximum amplitude of JF_646_-BAPTA signals collected from AMPARs (y-axis) or NMDARs (x-axis) at single synapses. Dashed line indicates equality. Using these data, we categorized synapses according to their receptor tone. **G)** Averaged synaptic transmission responses grouped by synapse category with number of synapses indicated. Dashed line indicates ΔF/F = 0 to guide the eye.

**Figure 8B** shows the averaged spontaneous transmission response from all iGluSnFR3 events we collected in each phase of the experiment (n=16,372 events). Spontaneously released single glutamatergic SVs stimulated NMDAR-mediated responses that peaked more slowly and took longer to decay than their AMPAR-mediated counterparts (**Figure S11**), consistent with the expected biophysics of these two iGluR types^42,45^. By contrast, blocking both iGluR types completely abolished Ca^2+^ signals correlated with iGluSnFR3 activity. In separate experiments, we found that recovery of spine-HaloTag after single-spine photobleaching proceeded with a time constant of several minutes (data not shown) and thus mobility of the reporter did not contribute substantially to the measured responses. To compare the reliability of responses to each receptor at each synapse, we calculated activation probabilities for each spine (**Figure 8C**). We observed that single glutamatergic vesicle release stimulated successful NMDAR-mediated transmission on 63.1% of attempts (5,651 attempts), while successful AMPAR-mediated transmission only occurred on 51% of attempts (6,266 attempts). By contrast, iGluR blockade resulted in 99.7% synaptic transmission failures (4,455 attempts). Among successful synaptic transmission events there was no difference in iGluSnFR3 event amplitude (**Figure 8D**), but NMDAR-mediated events possessed larger JF_646_-BAPTA event amplitudes. Thus, NMDARs activate more readily and trigger greater Ca^2+^ activity in response to single vesicles of glutamate than AMPARs, yet spontaneous transmission is still readily capable of triggering postsynaptic Ca^2+^ increases via AMPAR activation.

Our experimental design allowed us to compare intra-spine AMPAR-mediated and NMDAR-mediated responses, assess these against the bona fide synaptic transmission failures we observed during iGluR blockade, and ultimately classify spines as NMDAR-dominant, AMPAR-dominant, AMPAR-silent or NMDAR-silent (**Figure 8F**). When we sorted events according to their spine classification and averaged them, it was clear that our spine categories successfully separated iGluR-mediated Ca^2+^ signaling phenotypes (**Figure 8G**). Unsurprisingly, most Ca^2+^ signaling at spines was mediated by NMDARs with or without contributions from AMPARs (55.6% NMDAR-dominant, 15.8% AMPAR-silent), but the remaining 28.6% of the synapses assayed possessed substantial AMPAR-mediated activity.

The fast kinetics of the AMPAR-triggered Ca^2+^ responses suggest that calcium-permeable (CP)-AMPARs may contribute to these responses in AMPAR-dominant synapses. To explore this possibility, we looked for the distinctive aspiny morphology of parvalbumin-expressing interneurons, which makes them straightforward to recognize even without cell-type selective expression or molecular contrast^46^ (**Figure 9A**). These neurons should have high CP-AMPAR expression and few NMDARs^47^, in contrast to spiny excitatory neurons (e.g. pyramidal neurons, **Figure 9B**), which highly express NMDARs with few CP-AMPARs^47^. As predicted, most synapses we observed on the dendrites of an aspiny neuron were AMPAR-dominant or NMDAR-silent, consistent with strong CP-AMPAR expression and weak NMDAR expression; by contrast, the synapses onto a spiny neuron from the same imaging session were overwhelmingly NMDAR-dominant, consistent with weak CP-AMPAR expression and strong NMDAR expression (**Figure 9C**-**F**). **Figure 9E** compares the averaged spontaneous transmission responses collected from the aspiny and spiny neurons, supporting that these dendrites possess distinct receptor expression patterns. Thus, we leveraged our high-throughput imaging and analysis pipeline together with a novel dual-reporter approach to capture pre- to postsynaptic information transfer driven by single-vesicle signaling at single spines, offering a new lens through which to view heterogeneity in synaptic signaling and receptor expression across dendritic arbors. We expect our method will prove useful in understanding dendritic integration, synaptic clustering, and how inputs are patterned across dendrites to facilitate neuronal signaling across cell types and brain regions.

**Figure 9.**
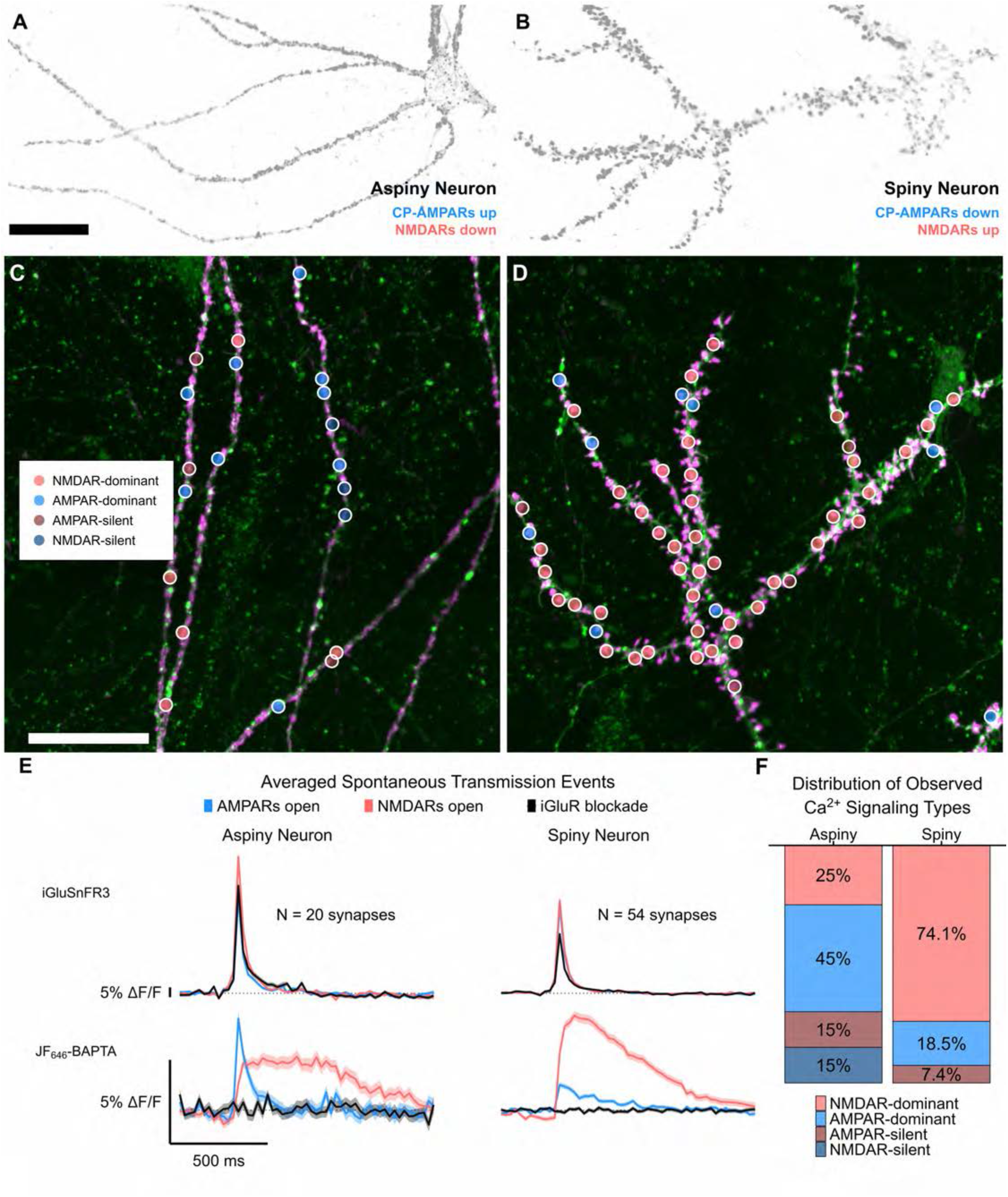
Synaptic transmission imaging captures predicted physiology of single-synapses according to cell-type. **A)** spine-HaloTag expression in an aspiny neuron. Aspiny neurons exhibit periodic nodes of actin along their dendrites which represent putative synaptic contacts. Excitatory synapses onto aspiny neurons are predicted to express higher levels of Ca^2+^-permeable AMPARs and lower levels of NMDARs relative to spiny neurons. Scale bar = 25 µm. **B)** spine-HaloTag expression in a spiny neuron. Excitatory synapses onto spiny neurons are predicted to express lower levels of Ca^2+^-permeable AMPARs and higher levels of NMDARs relative to aspiny neurons. Note that **A, B** have identical contrast, allowing direct comparison of spine-HaloTag expression by eye. **C,D)** Spatial mapping of single-spine receptor signaling properties at dendrites of **C)** the aspiny neuron in **A**, and **D)** the spiny neuron in **B**. Scale bar = 25 µm. iGluSnFR3 expression is shown in green and spine-HaloTag in magenta. **E)** Averaged spontaneous transmission events for aspiny and spiny neuron synapses from **C, D**. From the averaged traces, it is apparent that the aspiny neuron had higher expression of Ca^2+^-permeable AMPARs relative to NMDARs, whereas the spiny neuron had weak expression of Ca^2+^-permeable AMPARs relative to NMDARs. There are also kinetic differences, implying divergent expression of receptor subtypes and engagement of Ca^2+^ release processes secondary to receptor opening. **F)** Distribution of observed Ca^2+^ signaling types for each dendrite. As predicted, synapses in the aspiny neuron were primarily AMPAR-dominant, while the spiny neuron was overwhelmingly NMDAR-dominant.

## Discussion

We developed a high-throughput optical physiology approach which opens new lines of inquiry into synaptic functional heterogeneity across single neurons and circuits. Our approach combined three key technical innovations. First, we developed hardware control schemes in *Python* facilitating complex, versatile experiments. Second, we developed a suite of scripts across *ImageJ* and *MATLAB* for batch drift correction and segmentation of optical recordings. *R*-based file handling routines automatically matched ROIs to recordings of synaptic activity, allowing facile tracking of iGluSnFR3 activity at single synapses across 40+ imaging rounds. Third, our custom *R* algorithm corrected baselines and normalized intensity-time traces, enabling analysis of 10,000+ traces in tens of minutes. Notably, our algorithm should perform baseline correction, trace normalization, and feature extraction for other fluorescent biosensors, extending its usefulness for the neuroscience community. Together, these innovations constituted a scalable, unbiased data collection and analysis framework which enhanced the reproducibility and rigor of our experiments and shortened the time lag between experiments and publication-quality figures from weeks to days. While other sophisticated analysis solutions exist for calcium^15,16,18^ and voltage imaging^17^, we expect that our modular approach and its implementation in popular open-source software (e.g. *ImageJ*) should facilitate adoption of our workflow by the broader optical physiology community. Scalability may be less relevant for in vivo applications where it is common to analyze single neurons, but our approach lends itself to novel experimental designs that emphasize throughput and integration with multimodal analyses (e.g. immunocytochemistry, transcriptomics). Relating synaptic physiology to cellular identity may offer novel insights into basic mechanisms of neural circuit function. Further work could incorporate these methods into studying disease-relevant patient mutations in animal models or patient-derived neurons to help contextualize how high-penetrance mutations disrupt neuronal physiology.

With our high-throughput approach and our novel biosensor spine-HaloTag, we were able to image spontaneous glutamate release at single dendritic spines and subsequent receptor-mediated Ca^2+^ responses, directly visualizing the flow of information across the synaptic cleft with single-spine and single-vesicle resolution. We believe this method offers new avenues for careful biophysical analyses of synaptic transmission, evidenced by our ability to measure the receptor-specific probability of transmission at single spines. With pharmacology, we isolated Ca^2+^ responses mediated by NMDARs or AMPARs, and could categorize each spine based on the receptor type dominating postsynaptic Ca^2+^ elevations. Strikingly, release from single vesicles was sufficient to trigger substantial Ca^2+^ rises in a surprisingly large fraction of spines even with NMDARs blocked, suggesting the contribution of calcium-permeable (CP)-AMPARs or voltage-dependent Ca^2+^ channels, potentially also with subsequent release of Ca^2+^ from intracellular stores. Importantly, in an aspiny neuron where CP-AMPAR expression is known to dominate^47^, AMPAR-mediated Ca^2+^ transients were prominent and indeed dominated over NMDAR-mediated transients at a far higher proportion of synapses than in spiny neurons, demonstrating single-synapse analysis of synaptic transmission via CP-AMPARs. Thus, our synaptic transmission imaging and analysis approach captures single-spine receptor expression across the dendritic arbor by taking advantage of robust pharmacology to separate receptor types. These tools open new possibilities for directly linking the endogenous activity patterns of glutamatergic input to their outcomes in synaptic plasticity and dendritic integration.

The rich information content of our recordings enabled quantification of synapse heterogeneity across several axes of synapse physiology. Using UMAP^34^ and cluster analysis, we achieved data-driven classification of boutons based on their iGluSnFR3 responses to single APs. The three functional classes we observed existed along a gradient, in which boutons with the largest RRPs exhibited more glutamate release per AP, enhanced release probability, and faster iGluSnFR3 response kinetics. These data implied a developmental trajectory of bouton function, in which boutons expand their RRP with concomitant enhancements to presynaptic efficacy and kinetics. We tested this model by repeating these measurements across synapses at distinct culture ages (DIV14 and DIV21) with correlated immunocytochemistry of Munc13, allowing us to link culture age, synapse physiology, and molecular context. We observed that greater Munc13 expression predicted larger RRP sizes. As cultures aged, Munc13 content increased and synapses also exhibited greater presynaptic efficacy and faster kinetics in agreement with the model implied by our UMAP analysis. This feedback loop between unbiased analysis methods and targeted experimental design could prove useful in studying synaptic mechanisms in vivo, where the complexity of endogenous synaptic activity can be more challenging to parse.

Depending on culture age, synapse function was strongly predicted by Munc13 expression. At DIV14, Munc13 expression predicted glutamate release magnitude and variance, and the probability of observing iGluSnFR3 transients at all (P_iGlu_). Surprisingly, most of these correlations were lost at DIV21, though at this age, synapses still exhibited clear enhancements in presynaptic efficacy and iGluSnFR3 kinetics which coexisted with higher Munc13 expression. In both age cohorts, Munc13 content correlated with RRP size, consistent with the role of Munc13 in defining release sites^14^. However, RRP size was not different between DIV14 and DIV21 synapses, suggesting that DIV21 synapses might possess more Munc13 per release site or excess release sites which were not mobilized by single APs. Recent work shows that Munc13 content per release site does vary within the active zones of CA1 excitatory synapses^35^. Super-resolution imaging work from our lab has also observed inter-site variability in Munc13 content, in which release sites aligned with postsynaptic PSD-95 nanoclusters (i.e., a release site within a nanocolumn^28^) possess higher Munc13 density than those that are unaligned with postsynaptic PSD-95^48^. Though more work is needed to understand the consequences of inter-site variation in Munc13 content, the parsimonious interpretation is that not all release sites within an active zone are functionally equal. Interestingly, we analyzed our data using a binomial model with uniform release probability, but only ∼50% of the synapses we measured could be well-fit by this model. Our experimental design here precluded access to synapse-specific values for *Q* and therefore we could not leverage the non-uniform release probability model^30^, but future experiments could expand on this work by resolving the prevalence of synapses with non-uniform *P_r_* release sites and how these relate to Munc13 expression.

Synaptic Munc13 expression also predicted larger synaptic facilitation (PPR) at all ISIs tested at 1 mM Ca^2+^. At 2 mM Ca^2+^, this correlation was abolished, despite there being no difference in synaptic Munc13 expression between the two [Ca^2+^]_bath_ cohorts. These observations imply that higher Munc13 expression increases the number of low *P_r_* release sites which can only be engaged upon a second stimulus (facilitation) at 1 mM Ca^2+^, while the larger Ca^2+^ microdomains and elevated initial *P_r_* at 2 mM Ca^2+^ led to release site depletion (depression)^4^. We speculate that the observed heterogeneity in facilitation properties at synapses across dendritic arbors may be explained through the interplay of release site localization relative to VGCCs and inter-site variability in Munc13 expression, and how these properties vary from synapse-to-synapse. We noted that synapses produced the largest PPRs at ISIs ≤125 ms, consistent with the timing of transient SV docking^33^, in which additional SVs dock to the presynaptic membrane in a Ca^2+^-dependent manner within ∼100 ms of an AP. Though it is outside the scope of the present work, future work could deeply assay single-synapse paired-pulse dynamics using our automated microscopy and analysis approach in-tandem with post-hoc super-resolution imaging to examine how release site localization relative to VGCCs and expression of Munc13 interact with other relevant molecules (e.g. Synaptotagmin-7^8^) to produce heterogeneous short-term plasticity dynamics at single synapses.

As a suite of tools, our high-throughput approach enables observation of the broadly heterogeneous physiology among synapses along axonal and dendritic arbors and linking these properties to molecular expression by integrating live-cell physiology with post-hoc immunocytochemistry of the same synapses. Defining molecular context, its relationship to synapse physiology, and how these relationships are patterned across individual neurons is an important goal for neuroscience, as delineating these properties will aid the further development of biologically realistic models of neurons and how they become impaired in disease-relevant states. The approach demonstrated here enables these efforts and opens new lines of inquiry for synaptic neuroscience.

## Author Contributions

S.T.B. and T.A.B. designed research; S.T.B. performed research, wrote code, and analyzed data; A.D.L., M.C.A., M.C. contributed new reagents; S.T.B., A.D.L., and T.A.B. wrote the paper.

## Acknowledgements

This work was supported by 1R21MH141559-01 to A.D.L., F31MH124283 to M.C.A., and R37 MH080046 and R01 MH119826 to T.A.B. We thank the members of the Blanpied Laboratory for rigorous discussion and critical evaluation of the manuscript.

## Disclosure

Copyright 2024, University of Maryland, Baltimore. All rights reserved. Patent pending.

## Data Availability

The data from this paper to reproduce the figures and an example dataset to run *peakFinder.R* will be deposited in the Open Science Framework at the time of publication. Other data and materials used in this study are available from the corresponding authors by reasonable request. Source code for the analyses has been made available at https://github.com/blanpiedlab/HighThroughputOpticalPhysiology.

## Materials and Methods

### DNA constructs

pAAV.CAG.iGluSnFR3.v857.GPI (Addgene plasmid #178334) and pAAV.CAG.iGluSnFR3.v857.SGZ (Addgene plasmid #178335) were gifts from Kaspar Podgorski. pEF-Synaptophysin-mRuby was a gift from Edwin Chapman (Addgene plasmid #188980). LZF97_hSyn-spine-jRGECO1a (Addgene plasmid #119198) and PSD95.FingR.TagRFP were gifts from Don Arnold. psPAX2 (Addgene plasmid #12260) and pMD2.G (Addgene plasmid #12259) were gifts from Didier Trono. pFW_iGluSnFR3 was made by subcloning the promoter and open reading frame from pAAV.CAG.iGluSnFR3.v857.GPI into pFW using NEB HIFI Assembly. LZF97_hSyn-spine-HaloTag was made by replacing the jRGECO1a-TPR3-ZFBP sequence in LZF97_hSyn-spine-jRGECO1a with HaloTag using NEB HIFI Assembly. spine-HaloTag-P2A-iGluSnFR.SGZ was cloned from LZF97_hSyn-spine-HaloTag by first adding P2A at the end of the spine-HaloTag ORF by ligating a synthetic DNA sequence (Twist Biosciences) into the Esp3I site, then adding iGluSnFR.SGZ after P2A by NEB HIFI Assembly. All sequences were confirmed by whole plasmid sequencing (Plasmidsaurus) using Oxford Nanopore Technology with custom analysis and annotation.

### Lentivirus production in HEK cell culture

Lentivirus was produced in HEK293T cells (ATCC CRL-3216) maintained in DMEM + 10% FBS and penicillin/streptomycin at 37°C and 5% CO_2_. Cells were plated at 5x10^6^ cells/10 cm plate and transfected 12-24h later with 6 µg of either pFW_iGluSnFR3 or LZF97_hSyn-spine-HaloTag + 4 µg psPAX2 + 2 µg pMD2.G using PEI for 4-6 hours. After 48h, the virus-containing media was harvested, debris removed by centrifugation at 1000 RPM for 5 min and 0.45 µm PES filtering, and single use aliquots were frozen at -80°C for long term storage.

### Rat hippocampal neuron culture and transduction

All animal procedures were approved by the University of Maryland Animal Use and Care committee. Dissociated hippocampal cultures were prepared from E20 Sprague-Dawley rats of both sexes and plated on poly-L-lysine-coated coverslips (#1.5, 18 mm, Warner) at a density of 50,000 cells/coverslip, as previously described^41^. For experiments with iGluSnFR3 and Synaptophysin-mRuby/PSD95.FingR.TagRFP, neurons were transfected with 1 µg of pAAV.CAG.iGluSnFR3.v857.GPI and 1 µg pEF-Synaptophysin-mRuby/ PSD95.FingR.TagRFP at DIV14-16 with Lipofectamine 2000 per manufacturer instructions. For experiments with spine-HaloTag-P2A-iGluSnFR3.SGZ, 1 µg of the bicistronic plasmid was transfected at DIV8-12 with Lipofectamine 2000 per manufacturer instructions. Cells were plated at a total density of 50,000 cells/well. Neurons were imaged between DIV17-25.

### Widefield and confocal microscopy

Widefield and confocal images were acquired on a Nikon TI2 inverted microscope equipped with an Andor Dragonfly spinning disk confocal, a Plan Apo λD 60x/1.42 NA oil immersion objective. Excitation light (488/561/640 nm) was supplied by an Andor ILE and reflected to the sample through a 405/488/561/638 nm quadband polychroic (Chroma).

We recorded high-speed (200 Hz framerate) time-lapses of iGluSnFR3 activity in widefield mode, wherein the emission light bypassed the confocal unit to pass through appropriate emission filters (ET525/50, ET600/50, ET700/75 (Chroma)) to a Zyla 4.2+ sCMOS camera (Andor). 25.6 x 25.6 µm imaging regions were imaged at 20% laser power (488 nm, ∼8 W/cm^2^) with 5 ms exposures for experiments with iGluSnFR3 alone. For two-color experiments in **Figure 7** with iGluSnFR3 and spine-HaloTag, we simultaneously illuminated neurons with 488 and 640 nm laser lines, imaging 51.2 x 51.2 µm imaging regions, with 2x2 pixel binning, at 30% laser power (488/640 nm, ∼11-13 W/cm^2^) with 20 ms exposures. In two-color experiments in **Figure 8,9** with spineHaloTag-P2A-iGluSnFR3.SGZ, we simultaneously illuminated neurons with 488 and 640 nm laser lines, imaging 102.4 x 102.4 µm imaging regions, with 4x4 pixel binning, at 20% laser power on the 488 nm (∼7-9 W/cm^2^) and 7.5% laser power on the 640 nm (∼2-4 W/cm^2^) with 20 ms exposures. The emission light was split by a 565 nm long-pass dichroic mirror to split the light from iGluSnFR3 and spine-HaloTag to two Zyla 4.2+ sCMOS cameras (Andor). For high-resolution confocal z-stacks, emission light was passed through the confocal unit to the appropriate emission filters to a Zyla 4.2+ sCMOS camera. Neurons were imaged in confocal mode at 50% laser power (488/561/640 nm, ∼1-2 W/cm^2^) with 200 ms exposures and 10 µm z-stacks (step size = 0.3 µm) were acquired using a piezo-controlled stage (ASI).

### Glutamate imaging

For the experiments shown in **Figures 2-6**, cultured neurons co-expressing iGluSnFR3 and Synaptophysin-mRuby (or PSD95.FingR.TagRFP) were imaged on DIV17-25. For imaging, neurons were transferred to an imaging chamber with parallel platinum electrodes spaced by ∼1 cm and bathed in a modified Tyrode’s buffer containing (in mM): 136.5 NaCl, 3 KCl, 2 MgCl_2_, 1 CaCl_2_, 10 D-glucose, 10 HEPES, at pH 7.4 (adjusted with 1 M NaOH) with 20 µM DNQX and 100 µM DL-AP5 to block recurrent excitation. When [Ca^2+^] was varied, NaCl was iso-osmotically substituted with CaCl_2_ to maintain a nominal osmolarity of 308 mOsm. To help maintain temperature in the bath, the objective was heated to 37 °C with a heating collar (TOKAI HIT USA Inc., USA). In experiments with stimulus-evoked iGluSnFR3 activity, field stimuli (10 V/cm, 1 ms) were delivered by a stimulator box (S88X Square Pulse Stimulator, Grass Instrument Co., USA) triggered externally by a programmable stimulus generator (Master-8, A.M.P. Instruments, Israel). Generally, we selected 25.6 x 25.6 µm imaging regions with abundant iGluSnFR3^+^ axonal processes and Synaptophysin-mRuby puncta; these were subjected to a battery of stimulus protocols with 20 s rest periods in between protocols. Once the protocol sequence was complete, this was repeated at the next imaging region. When we imaged spontaneous iGluSnFR3 activity, the bath solution also contained 1 µM TTX to prevent APs. We repeatedly imaged single regions for 30 s at 200 Hz (5 s rest periods in between, 6-9 trials per region) to capture spontaneously released glutamate at individual boutons.

### Post-hoc immunocytochemistry

For experiments with post-hoc immunocytochemistry (main-text **Figure 5,6**), we devised a simple protocol that allowed robust identification and alignment of the same imaging regions between the live and post-hoc imaging sessions. At the end of each live-cell imaging session, the XY-stage coordinates of each imaging position were saved to a .csv file for later, and a 5x5 tile of the coverslip was taken at 10x magnification, using the 488 nm laser to image iGluSnFR3 expression across many neurons. Neurons were immediately fixed in 2% paraformaldehyde (PFA) / 2% sucrose in 1X phosphate-buffered saline (PBS) for 10 minutes at room temperature and washed three times for 5 minutes in PBS/glycine (PBS + 100 mM glycine). At this point, neurons were either stored at 4°C for 24-48 hours before immunocytochemical labeling or immediately processed.

Immunocytochemistry proceeded using a standard procedure with all steps occurring at room temperature unless otherwise indicated: 1) Permeabilization with 0.3% Triton-X-100 in PBS/glycine for 20 minutes while shaking at 80 rpm; 2) Washed three times for 5 minutes in PBS/glycine while shaking; 3) Incubated coverslips inverted on parafilm with Image-IT (Invitrogen, #136933) for 30 minutes; 4) Washed three times for 5 minutes in PBS/glycine while shaking; 5) Blocking in Blocking Buffer (10% donkey serum, 0.2% Triton-X-100 in PBS/glycine) for 60 minutes shaking; 6) Incubated inverted on parafilm with primary antibody in 1:1 mixture of PBS/Glycine and Blocking Buffer (1:250 Rabbit Anti-Munc13-1, Synaptic Systems, Cat No. 126103) overnight at 4°C; 7) Washed three times for 5 minutes in PBS/glycine while shaking; 8) Incubated inverted on parafilm with secondary antibody (1:500, Donkey Anti-rabbit Alexa Fluor 647, Invitrogen A31573) in PBS/glycine; 9) Washed three times for 5 minutes in PBS/glycine while shaking. Stained cells were then stored at 4°C until they were imaged later the same day.

To align the imaging sessions, we again acquired 5x5 tiles of iGluSnFR3 expression using the 488 nm laser to sample neuron orientation, and identified recognizable neurons present in both imaging sessions. We repeatedly imaged neurons with well-defined morphology that were seen in both sessions, and carefully aligned them in this manner. From here, we used the previously recorded stage positions from the corresponding live-cell session to find the approximate imaging region from the live-cell imaging session, and scanned until the correct imaging region could be identified and aligned.

### Simultaneous glutamate and Ca^2+^ imaging

For the experiments shown in **Figure 6**, cultured neurons co-expressing iGluSnFR3 and spine-HaloTag were prepared for imaging on DIV17-25. We achieved spine-localized Ca^2+^ imaging by labeling spine-HaloTag with JF_646_-BAPTA-HaloTagLigand (HTL)-AM. JF_646_-BAPTA-HTL-AM was a gift from Luke Lavis^40^. We aliquoted JF_646_-BAPTA-HTL-AM as a 1 mM stock in anhydrous dimethyl sulfoxide (DMSO) and stored at -80 °C.

To treat cells with JF_646_-BAPTA-HTL-AM, we followed a protocol similar to what has been previously described by Bradberry et al. 2021^49^. Conditioned media (3x300 µL) was transferred from a well containing a coverslip with cultured neurons to an empty 12-well plate. For each experiment, we made a working stock of JF_646_-BATPA-HTL-AM diluted in a modified Tyrode’s buffer with the following composition (in mM): 136.5 NaCl, 3 KCl, 2 MgCl_2_, 1 CaCl_2_, 10 D-glucose, 10 HEPES and 2 µM JF_646_-BAPTA-HTL-AM. For each coverslip incubation, we added 300 µL of the JF_646_-BAPTA-HTL-AM working stock to a well with 300 µL of conditioned media, for a 1 µM effective dye concentration during incubation.

Coverslips of cultured neurons were transferred to the dye-containing wells and incubated for 30 minutes at 37 °C. Following incubation, coverslips were immediately transferred sequentially to the other two wells with conditioned media to wash off free JF_646_-BAPTA-HTL-AM before being returned to their original 12-well plate for recovery. Neurons were allowed to recover from the dye incubation for 30-60 minutes at 37 °C. For imaging, neurons were transferred to an imaging chamber (Warner Instruments, USA) and bathed in a modified Tyrode’s buffer containing (in mM): 135 NaCl, 3 KCl, 4 CaCl_2_, 0 MgCl_2_, 10 D-glucose, 10 HEPES, 1 Trolox (to preserve fluorescence of JF_646_-BAPTA-HTL-AM), and 1 µM TTX in a climate-controlled chamber (37 °C, 100% humidity, TOKAI HIT USA Inc., USA). Spontaneous glutamate release and subsequent NMDAR-mediated Ca^2+^ flux at single dendritic spines were observed for 20 s at 50 Hz before moving to another imaging region or washing in solution containing 100 µM DL-AP5. After wash-in, imaging of the same regions was repeated.

For single-spine pharmacology experiments (**Figure 8,9**), several imaging regions were recorded with the following sequence: imaging synaptic transmission at baseline, with AMPARs open and NMDARs closed (+100 μM DL-APV, no DNQX) in the imaging buffer described above; wash-in of 50 mL of imaging buffer with no APV, 50 μM DNQX, and 100 μM D-serine to open NMDARs and close AMPARs while maximizing NMDAR co-agonist availability followed by imaging for 1 minute; wash-in of 50 mL of imaging buffer with 100 μM DL-APV, 50 μM DNQX to block NMDARs and AMPARs.

### Hardware automation

As described by **Figure 1A**, we developed a hardware automation scheme to fully automate imaging experiments, improving throughput/reproducibility. A custom *Python* script enabled control of the microscope and stage (i.e. imaging positions), image acquisition, electrical stimulation, and solution exchange.

The microscope and stage piezo were controlled from the script via the Andor (Oxford Instruments) REST API and Fusion software. This approach allowed us to store stage positions of multiple regions in a Python list object and iterate protocols over chosen imaging regions. We developed custom functions to automate updates to key imaging parameters (e.g. camera settings, filenames, focus stabilization) throughout the protocol, enabling more complex and robust automated imaging sequences.

Electrical field stimulation was controlled via the Master-8 and S88x Square Pulse Stimulator. At the beginning of each imaging protocol with electrical stimulus, we first established USB serial communication with the Master-8. To automatically update stimulus paradigms, we used a dictionary data structure to link stimulus paradigm parameters to imaging protocols – this enabled us to pre-configure the Master-8 prior to each imaging protocol to deliver diverse stimulus paradigms (e.g. single stimuli, paired stimuli, stimulus trains). To trigger stimulation during imaging, we predefined the frame in each imaging protocol (Fusion) during which a TTL pulse would be sent via BNC cable to the Master-8. After receiving the TTL from the microscope, the Master-8 triggered field stimulus of neurons by sending its pulse sequence via BNC cable to the S88X Square Pulse Stimulator.

The workflow was similar for solution exchange. Sub-functions established USB serial communication with a ValveLink8.2 Controller (AutoMate Scientific), which allowed us to toggle pinch valves on a gravity flow perfusion apparatus, initiating fluid flow. To achieve fluid exchange, we simultaneously triggered an Arduino microcontroller (Arduino, Italy) which controlled a peristaltic pump, vacuuming excess fluid from the imaging chamber^50^. The gravity flow perfusion apparatus was adjusted to a flow rate of ∼1.5 mL/min.

### Image segmentation and extraction of intensity-time traces

At the end of each iGluSnFR3 imaging session, confocal z-stacks were recorded of the same imaging fields to capture the position of Synaptophysin-mRuby or PSD95.FingR.TagRFP puncta (i.e. boutons or excitatory synapses) along the axonal or dendritic arbor. Most confocal z-stack image processing was performed with custom macros in FIJI as we have previously described^45^. Background subtraction was performed by identifying the lowest 1% of pixel intensity per channel and subtracting this value from its respective image channel. We constructed a mask of the axonal or dendritic arbor by automatically thresholding the iGluSnFR3 channel on the top 5% of pixel intensities. Binary axonal arbor masks were smoothed and small background particles were removed. Putative synaptic puncta were identified in the synaptic marker channel with the plugin SynQuant. Only ROIs within the boundary of the mask of the axonal or dendritic arbor were analyzed. After drift correcting for small stage movements between perfusion rounds at single imaging regions, we distributed the “ROI maps” to folders containing individual imaging trials. A custom FIJI macro iterated through all subdirectories, opening videos and applying their respective ROI maps, populating a separate directory with .csv files of the intensity-time recordings at each ROI, which we then read into *R* for analysis.

For activity-based segmentation, we applied a method described in Mendonca et al. 2022^24^ with some modifications. Briefly, a moving average filter with a 5-point span was used to smooth the temporal profile of the iGluSnFR3 responses. A band-pass Gaussian filter (0.05-200 Hz) was then applied to amplify the iGluSnFR3 signal. At each pixel we subtracted the mean value and divided by the standard deviation, which had the effect of suppressing inactive stretches of the axonal arbor and amplifying active stretches (the “activity footprint” of the axon). From these, we created max intensity projections which were thresholded on the top 3% of all pixels to generate ROI maps of activity along the axonal arbor. We applied these “activity maps” to their respective videos and extracted .csv files of intensity-time recordings for each ROI, which we then read into *R* for analysis.

### Immunocytochemistry analysis and linking regions-of-interest

Post-hoc immunocytochemistry imaging sessions were drift-corrected to align to their corresponding live-cell imaging session using the iGluSnFR3 imaging channel of a max intensity-projected, high-resolution confocal z-stack. A custom ImageJ macro used the plugin SynQuant to identify PSD95.FingR.TagRFP puncta, and measured the intensity of both PSD95.FingR.TagRFP and labeled Munc13 at each ROI. ROIs were then linked to their correlated live-cell imaging session ROI by performing a nearest-neighbor analysis. If the centroid of the identified post-hoc ROI was within 3 µm of its live-cell nearest neighbor, the ROIs were considered to be the same between imaging sessions. Correlational analyses were performed only on synapses with linked post-hoc immunocytochemical quantification.

### Fluorescence signal extraction

To normalize iGluSnFR3 intensity-time traces to ΔF/F, correct baseline fluctuations, and extract fluorescence signals at scale, we wrote a custom algorithm in *R* (*peakFinder.R*). The algorithm proceeds in three stages, iteratively refining its approximation of the trace’s baseline. First, the algorithm flagged outlier indices as those which rise above a threshold of σ (standard deviation) from the rolling median (0.75 s span) of the raw fluorescence intensity (median filter). Once a first approximation of the outliers is known, a more refined approximation of the baseline, F, was made on the raw intensity trace using a rolling median (0.75 s span) which excluded the known outlier indices. The outlier indices were replaced with the last non-NA value (i.e. known intensity at non-signal indices) using a “last <u>o</u>bservation <u>c</u>arried forward” function in *R* (*na.locf*()).Then, traces were adjusted to ΔF/F by dividing the raw intensity by the approximate baseline. Using a Schmitt trigger thresholding approach, we identified putative fluorescence signals when they exceeded an upper threshold of 3.5σ; the signals terminated when they decayed below a lower threshold of 1.5σ; the indices corresponding to these putative fluorescence signals were flagged. To identify the full extent of putative fluorescence signals and properly exclude them from the baseline approximation, additional points before and after putative fluorescence signals were flagged as signal indices according to the known rise and decay kinetics of the fluorescent sensor being imaged. With the original outliers and the putative signal indices flagged, a new baseline (F) was calculated from the rolling average of the raw intensity trace excluding outliers and putative signals, and the final iteration of ΔF/F was calculated. The final detection of fluorescence transients was achieved with a more stringent threshold (lower, 1.5σ, upper, 5σ) and annotated for further analysis.

For the experiments in **Figure 6**, we found that the slower kinetics of JF_646_ and the frequent convolution of JF_646_ transients with one another made it difficult to approximate the baseline and accurately convert traces to ΔF/F using a median filter. Instead, we used a percentile filter approach, wherein the trace was divided into 10 equivalent time bins (e.g. 0-2 s, 2-4 s, etc.), and indices which comprised the bottom 30% of intensity values were identified in each bin. Similar to above, these were flagged as putative baseline indices, and the baseline fluorescence intensity at these indices were interpolated using *na.locf()*. A rolling average of these points with a 0.75 s span was calculated to approximate the baseline. The trace was then adjusted to ΔF/F, and putative signals identified with a Schmitt trigger (lower, 1.5σ, upper, 3.5σ). We then repeated this process with the putative signals being excluded from the percentile filter. The final iteration of ΔF/F was calculated and JF_646_ transients were once again identified with a stringent threshold (lower, 1.5σ, upper, 5σ) and annotated for further analysis.

### Analysis of single stimulus-evoked and spontaneous iGluSnFR3 transients

Annotated iGluSnFR3 transients from stimulus-evoked and spontaneous recordings were analyzed using custom analysis scripts in *R*. We measured the peak ΔF/F of found transients, and for stimulus-evoked transients, we calculated the time delay (Δt) between the stimulus and the peak ΔF/F to measure the speed with which iGluSnFR3 reports stimulus-secretion coupling. To measure the rise time, t_rise_, we established a time cutoff of corresponding to the time of stimulus, prior to the time of peak ΔF/F. We then found the minimum ΔF/F prior to the peak maximum, and solved for the slope of a line between the minimum ΔF/F and the peak ΔF/F. From the peak ΔF/F, we established the 10% and 90% fluorescence levels for each transient and calculated the t_rise_ between these two points based on the found rise slope for each transient. To measure the decay time constant, τ_decay_, for each iGluSnFR3 transient, we used the package *nlstools*^51^ to fit an exponential decay function, f(t) = A*e*^!*t*/$^*^decay^* from the peak ΔF/F to the end of the annotated transient. From the calculated values for the pre-exponential factor, *A*, and τ_decay_, we calculated the decay time from 90% to 10% of the peak ΔF/F, t_decay_. Similarly, to calculate the full-width at half-maximum (t_1/2_) of each transient, we used the 50% ΔF/F value for each peak to solve for the time of 50% ΔF/F on the peak rise (linear fit) and the peak decay (exponential decay fit), and subtracted these values to generate t_1/2_. The procedure was identical for spontaneous iGluSnFR3 transients, except for the calculation of t_rise_. Since there was no stimulus to mark the beginning of an iGluSnFR3 transient, we established our time cutoff as 25 ms prior to the peak ΔF/F (informed by the published rise kinetics of iGluSnFR3, ∼19 ms^19^). Otherwise, peak statistics were measured identically (omitting Δt, which could not be calculated). For both stimulus-evoked and spontaneous transients, signals were filtered out of the dataset if τ_decay_ < 15 ms or τ_decay_ > 300 ms, as we determined these represented spurious, high-frequency noise or non-glutamate release activity (e.g. iGluSnFR3^+^ vesicle trafficking through the axon, which had a characteristically slow time constant uncoupled from the stimulus), respectively.

We calculated several properties of synapses from our stimulus-evoked iGluSnFR3 trials. The probability of observing an iGluSnFR3 transient (P_iGlu_) was defined as a binary for each trial: 1 if a signal was detected, or 0 if there was no signal detected. The sum of these binary values was divided by the total number of trials to define P_iGlu_ for an ROI at a single [Ca^2+^]_bath_. To assess whether the stimulus-evoked glutamate release was mono-or multivesicular, we compared the average peak ΔF/F of stimulus-evoked iGluSnFR3 transients for each ROI (E_ΔF/F_) to the average peak ΔF/F of spontaneous iGluSnFR3 transients at each [Ca^2+^]_bath_ (S_ΔF/F_, see **Figure 3**), which represented putative single vesicle glutamate release events. When the value of the derived metric E_ΔF/F_/S_ΔF/F_ was >1, we interpreted this as the release of multiple vesicles (but see *Discussion*). The coefficients of variation (CVs, mean / σ) were calculated for all statistics at each ROI. For pair-wise comparisons, we calculated statistical significance via Kolmogorov-Smirnov and Mann-Whitney-Wilcoxon tests. To compare statistics across [Ca^2+^]_bath_, we calculated one-way analysis of variance (ANOVA).

### Mean-variance analysis of iGluSnFR3 transients

To perform mean-variance analysis (MVA) of iGluSnFR3 transients, we sampled the single action potential (AP)-evoked iGluSnFR3 activity at individual boutons across multiple [Ca^2+^]_bath_ (sampling multiple *P_r_* states). To accurately measure the mean and the variance of the iGluSnFR3 ΔF/F for each stimulus, we used an approach that was agnostic to the algorithmic definition of an iGluSnFR3 signal. The maximum of the iGluSnFR3 ΔF/F was measured in a time window starting from the stimulus (t = 0) and extending to 250 ms post-stimulus. This enabled measurement of sub-threshold maxima and noise, especially at 0.5 mM Ca^2+^, where most boutons were quiescent. We then converted the measured amplitudes to E_ΔF/F_/S_ΔF/F_ and measured their variance. Each bouton possessed four data points of the form (x, y) corresponding to (mean, variance) at each [Ca^2+^]_bath_, and each bouton was fit using the uniform probability binomial model of the form: σ^2^ = *Qx̅ - x̅^2^/N_sites_^30^*. In fitting the binomial model, *N_sites_* was constrained to positive values < 100 and *Q* was constrained to values between 0.8-1.2, as E_ΔF/F_/S_ΔF/F_ = 1 was expected to represent the iGluSnFR3 response due to a single vesicle, per the analyses in **Figure 3**. In **Figure 5**, we followed this same workflow, but using synapses defined by PSD95.FingR.TagRFP. In addition, we added another [Ca^2+^]_bath_ sample at 1.5 mM.

Boutons/synapses were accepted for analysis if the sum of the squared residuals was ≤ 1 and their maximum *P_r_* > 0.45^14^. From the binomial model, we calculated the following synaptic parameters: the uniform release probability (*P_r_*) at each [Ca^2+^]_bath_, the quantal size of the iGluSnFR3 response (*Q*), and the total number of release sites in the readily releasable pool (*N_sites_*).

### Generating a UMAP representation of bouton physiology

To cluster boutons into functionally distinct classes (**Figure 4**), we calculated a collection of statistics which summarized the intrinsic behavior of boutons during single AP trials. First, only iGluSnFR3 transients which exceeded our threshold of detection (see above, *Fluorescence signal extraction*) were used for calculating summary statistics. The probability of observing an iGluSnFR3 transient, P_iGlu_, was the ratio of the number of transients identified at a given bouton to the number of single stimulus trials administered. Transients were only counted toward P_iGlu_ if a transient maximum occurred within 500 ms of a preceding stimulus. P_iGlu_ was calculated per bouton for each [Ca^2+^]_bath_. We also calculated the mean amplitude and the variance for iGluSnFR3 transients in terms of E_ΔF/F_/S_ΔF/F_ at each [Ca^2+^]_bath_. Because the mean and variance were often unavailable for boutons which possessed no identifiable iGluSnFR3 transients at 0.5 mM Ca^2+^, boutons with *NA* were coerced to 0 for the purposes of the analysis.

For the remaining statistics, we restricted our analysis to [Ca^2+^]_bath_ = 1, 2, or 4 mM, as data was too sparse to calculate coefficients of variation (CVs) at 0.5 mM Ca^2+^. We calculated the mean decay time constant, τ_decay_, as well as the CV of peak ΔF/F, CV of τ_decay_, and CV of Δt. The mean Δt was omitted as it exhibited little variation in magnitude across boutons (unlike mean amplitude and mean τ_decay_).

For all statistics (except for P_iGlu_, which was already normalized between 0 and 1), missing values were coerced to 0 and the values in each column were normalized between 0 and 1, equally weighting all of the statistics in the data matrix. This resulted in 24 data columns (each [Ca^2+^]_bath_ constituted an additional observation for each of the 7 summary statistics). We used the *umap* and *dbscan* libraries in *R* to generate UMAP scores for the data matrix and cluster the UMAP score output (DBSCAN minimum points per cluster = 10). Cluster identities were re-associated with their original bouton IDs, which enabled us to categorize our bouton physiology data according to their UMAP class. Grouping the data in this way allowed us to visualize the defining characteristics of each bouton class (**Figure 4E-H**).

### Analysis of iGluSnFR3 transients evoked by paired stimuli

The protocol for the paired pulse experiment in **Figure 5** was administered with each protocol in triplicate. The protocol sequence began with a Test Pulse (a single stimulus), followed by paired pulses with inter-stimulus intervals (ISIs) of 60, 75, 100, 150, or 500 ms. There was a 20 s recovery period between each trial. This protocol sequence was first administered in a bath solution with 0.5 mM Ca^2+^. In between iterations of the protocol sequences, buffer solution containing 1 mM Ca^2+^ or 2 mM Ca^2+^ was washed in.

We averaged the responses from the three Test Pulse trials to generate the average response to a single stimulus for each ROI and each [Ca^2+^]_bath_ measured. We aligned the average response to the test pulse to the first stimulus in each paired pulse recording and subtracted the test pulse from the paired pulse recording, enabling more accurate measurement of the amplitude of the 2^nd^ iGluSnFR3 transient (see **Figure S6B**). This was especially important in cases where a short ISI resulted in a convolution of the two iGluSnFR3 transients (e.g. ISI = 60, 75, or 100 ms).

We performed this operation for every ROI in the dataset and then divided the traces into two stimulus epochs: the first stimulus epoch began at the first stimulus and ended at the second stimulus; the second stimulus epoch began with the second stimulus and terminated 200 ms later. For each trial, we identified the amplitude of the first pulse (Pulse_1_) as the maximum ΔF/F within the first stimulus epoch of the unmodified trace. To determine the amplitude of the second pulse (Pulse_2_), we identified the maximum ΔF/F within the second stimulus epoch in the subtracted trace.

### Analysis of simultaneous iGluSnFR3 and JF_646_-BAPTA-AM recordings

We segmented simultaneous recordings of iGluSnFR3 and JF_646_-BAPTA-HTL-AM on the activity in the iGluSnFR3 channel as described above. These iGluSnFR3 activity maps were distributed to folders containing the recording for either the iGluSnFR3 or JF_646_ channel and intensity-time traces were extracted as before. Because the ROI maps were identical between channels of the same recording, we were able to pair the traces by ROI in *R*. To characterize putative synaptic transmission events (paired iGluSnFR3/JF_646_ signals), we took advantage of our robust peak discrimination in the iGluSnFR3 channel. Around each iGluSnFR3 signal identified by our peak finding algorithm, we defined a temporal window in which to search for a local maximum in the JF_646_ channel (**Figure 6**). The lower boundary of the temporal window was 10 ms prior to the iGluSnFR3 peak maximum, and the upper boundary of the temporal window was 250 ms after the iGluSnFR3 peak maximum. In this way, we were able to pair the local maxima (which reflected the peak of paired Ca^2+^ transients) in each channel for bona fide glutamate release events, which could then be analyzed. This approach enabled exponential decay analysis and averaging operations using synaptic transmission events.

To discriminate putative transmission events from noise, we took advantage of the output from our peak finding algorithm, which tabulates putative baseline indices as part of the baseline adjustment routine. For each dendritic spine in the dataset, we selected the bottom 50% of all indices for the JF_646_-BAPTA channel and calculated the standard deviation of these points. A signal threshold of 5σ was set for each spine, and JF_646_-BAPTA signals above this threshold were considered transmission.

To discriminate synapse Ca^2+^ signaling phenotypes described in **Figure 8**, **9**, we identified the amplitude of the maximum AMPAR-mediated Ca^2+^ event, the maximum NMDAR-mediated Ca^2+^ event, and the maximum iGluR blockade event. Then, we calculated the following ratios for each synapse: AMPAR/NMDAR, NMDAR/AMPAR, and the corresponding null ratios (blockade/NMDAR, blockade/AMPAR), which represented the putative detection limit for each synapse. Synapses could then be categorized through logical statements. For example, synapses which were NMDAR-dominant needed to have NMDAR transmission > AMPAR transmission but AMPAR transmission also needed to be present (i.e. NMDAR/AMPAR > blockade/AMPAR & AMPAR/NMDAR > blockade/NMDAR), whereas AMPAR-silent synapses had NMDAR transmission > AMPAR transmission but blockade > AMPAR transmission. This approach helped to normalize each synapse’s responses and allowed each synapse to act as its own benchmark for SNR, as the null ratios denoted bona fide synaptic transmission failure according to the noise level of each synapse.

### Data Visualization

All data analysis and visualization was carried out using custom scripts in *R*. In general, we relied on structured data frames with several string variables by which data could be sorted and analyzed. Figures were organized into multi-panel layouts and plotted using a combination of the packages *ggplot2* and *gridExtra*.

For maps of synaptic function, we used a custom workflow and the packages *RImageJROI* and *EBImage*. First, we generated .png files of the max-projected confocal z-stacks for individual imaging regions. These .png files were read into *R* as raster objects. Zip files containing the ROI information were generated for each imaging region in *FIJI*. Using *RImageJROI*, we were able to read the .zip files into *R*. Using *ggplot2*, we layered the ROI polygons from *FIJI* on top of the raster of the imaging region. Statistics could then be assigned to each ROI polygon on the basis of their unique ROI identity. In later experiments with bona fide synaptic localization (**Figure 5,6,9**), we simply distributed ROIs according to their centroids. In **Figure 5** and corresponding supplemental figures, we flattened a binarized mask overlay of the live-cell dendritic arbor onto a .png file of the corresponding post-hoc immunocytochemistry imaging region. Post-hoc images were aligned to the live cell imaging session. In our implementation, insets centered around selected ROIs could be extracted for zoom-ins of smaller regions.

### Statistics

Statistical analyses were performed in *R*. Statistical differences between datasets were tested with pair-wise Mann-Whitney-Wilcoxon and Kolmogorov-Smirnov tests. For datasets with multiple groups, data were also tested with one-way ANOVA. For correlational analyses, we calculated Pearson or Spearman correlation coefficients, depending on the linearity of the distribution.

## Supplementary Figures

**Figure S1.**
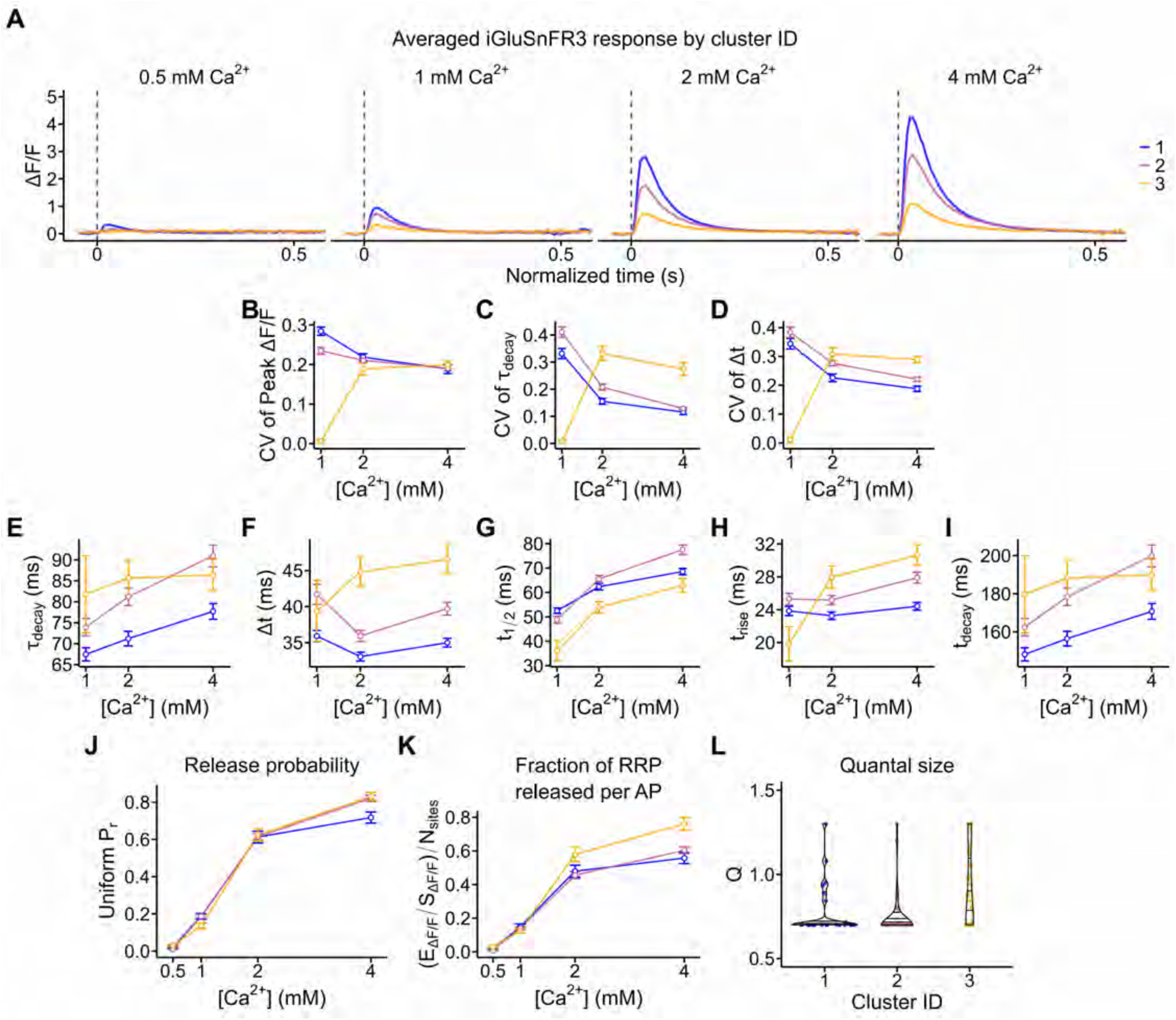
Additional functional properties of the three bouton classes from main-text Figure 4 imply differences in the SV release machinery. **A)** The average iGluSnFR3 response in response to stimulus across [Ca^2+^]_bath_ and cluster ID. For the plots presented in B-G, 0.5 mM Ca^2+^ has been omitted due to the sparse data in cluster 2 and 3. B) Coefficient of variation (CV) of peak ΔF/F vs. [Ca^2+^]_bath_. C) CV of τ_decay_ vs. [Ca^2+^]_bath_. D) CV of Δt vs. [Ca^2+^]_bath_. E) τ_decay_ vs. [Ca^2+^]_bath_. F) Δt vs. [Ca^2+^]_bath_. **G)** t_1/2_ vs. [Ca^2+^]_bath_. **H)** t_rise_ (10-90%) vs. [Ca^2+^]_bath_. **I)** t_decay_ (90-10%) vs. [Ca^2+^]_bath_. **J)** Uniform release probability, *P_r_* vs. [Ca^2+^]_bath_. **K)** RRP fraction released per AP (E_ΔF/F_/S_ΔF/F_)/N_sites_ vs. [Ca^2+^]_bath_. **L)** Binomial model Q vs. cluster ID.

**Figure S2.**
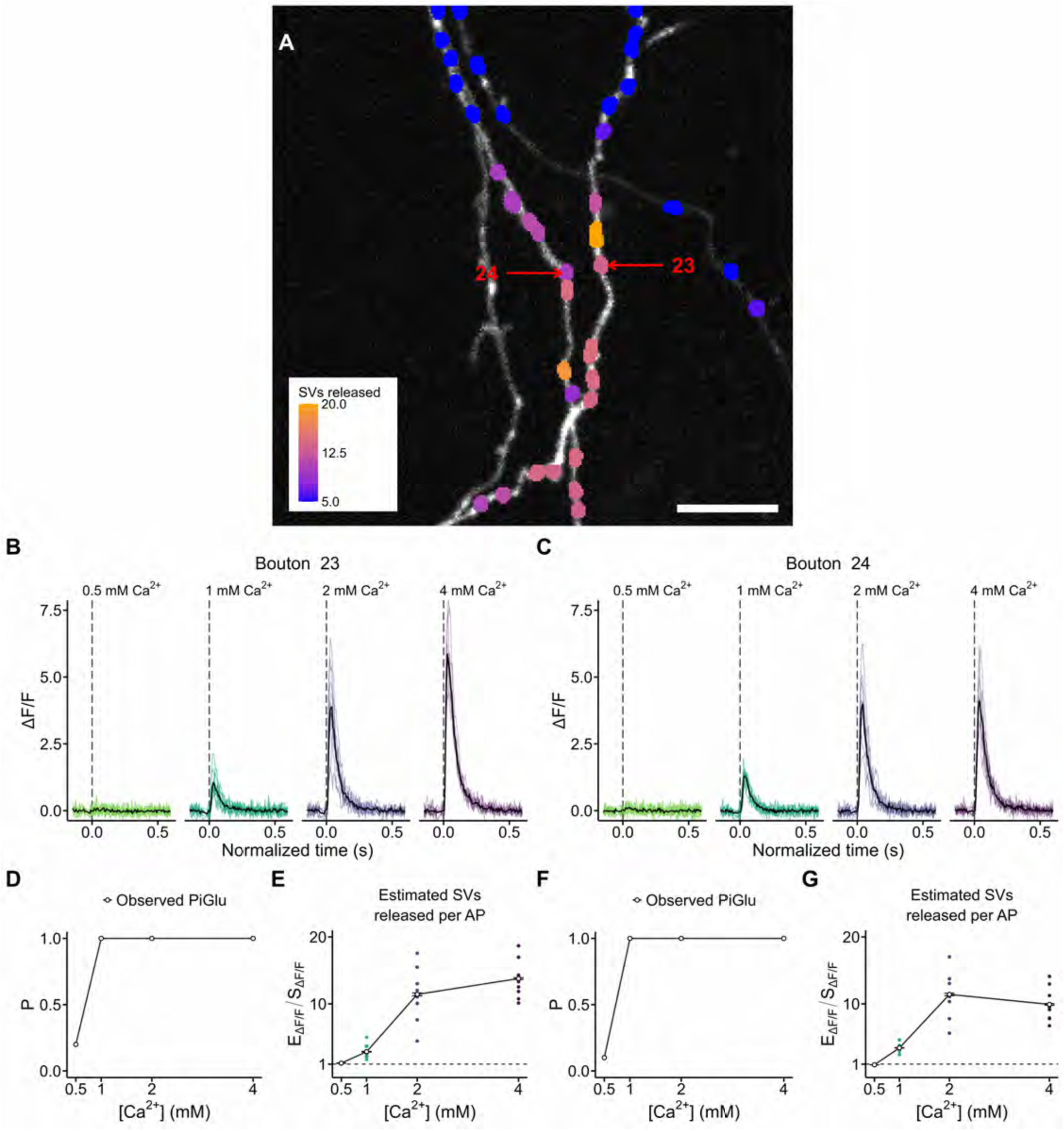
Boutons separated by ∼1.8 µm exhibit divergent iGluSnFR3 behavior. **A)** The imaging region from Figure 4I, color-coded according to number of SVs released at 4 mM Ca^2+^. Boutons 23 and 24 are indicated for further analysis. Scale bar = 5 µm. **B, C)** The iGluSnFR3 responses at all [Ca^2+^]_bath_ tested. Though the boutons are only separated by 1.8 µm on separate branches of the same axon, they exhibit divergent iGluSnFR3 behavior. **D,F)** P_iGlu_ vs. [Ca^2+^]_bath_. **E,G)** Estimated SVs released per AP (E_ΔF/F_/ S_ΔF/F_) vs. [Ca^2+^]_bath_. Note that Bouton 24 appears to reaches its maximal response at 2 mM Ca^2+^, with little change in behavior at 4 mM Ca^2+^.

**Figure S3.**
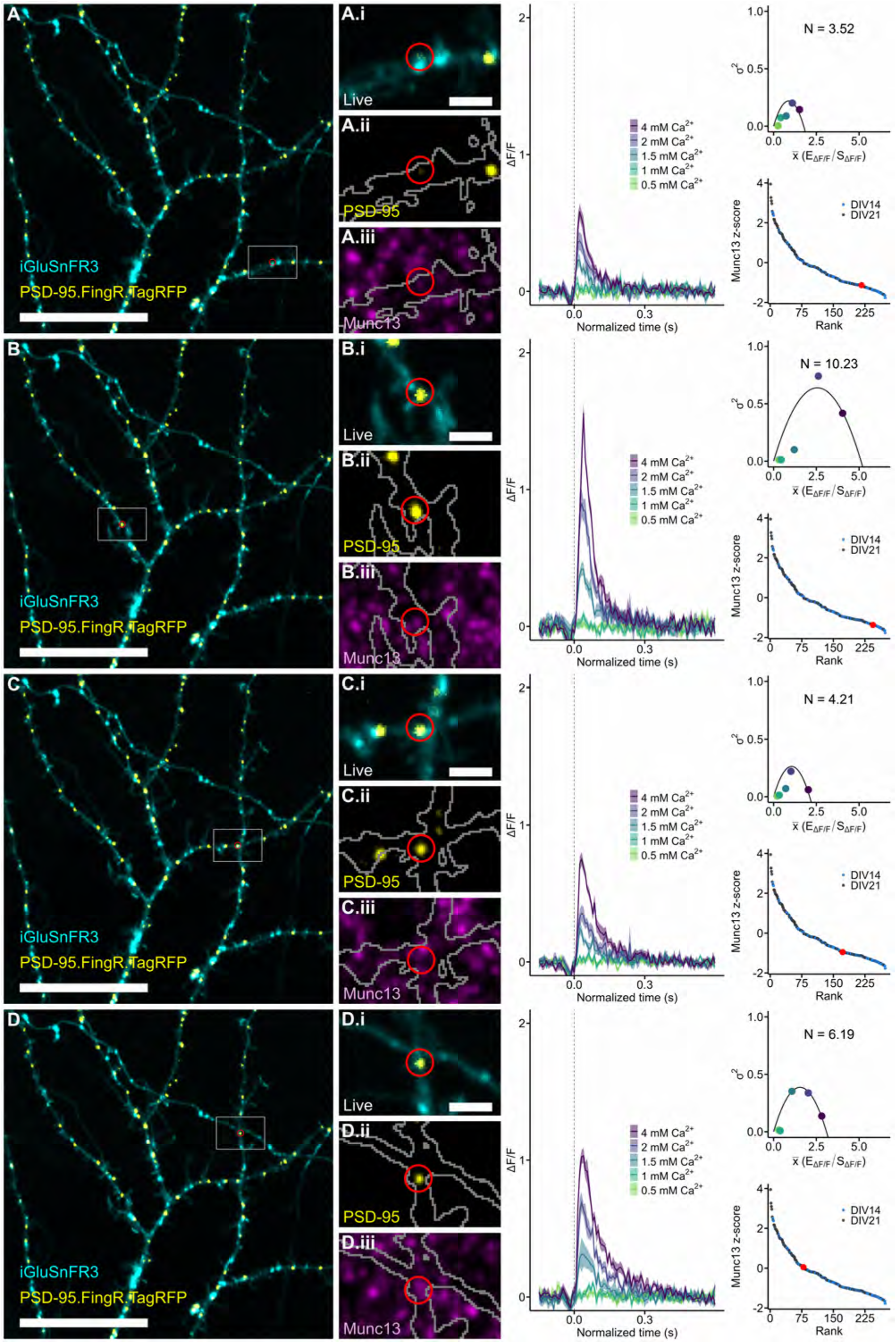
Single-synapse reports for selected DIV14 synapses (main-text Figure 5). **A)** Full imaging region of DIV14 dendrite expressing iGluSnFR3 (cyan) and PSD-95.FingR.TagRFP (yellow). Scale bar = 20 µm. **(i)** Inset of the imaging region shown in **A**. Red circle indicates the synapse selected for analysis. Scale bar = 2 µm. **(ii)** Post-hoc imaging of PSD-95.FingR.TagRFP at higher illumination intensity. Grey outline indicates a mask of the live iGluSnFR3 signal. **(iii)** Post-hoc imaging of Munc13 immunofluorescence. **Right**, the single synapse iGluSnFR3 response to single APs at each [Ca^2+^]_bath_ tested, mean-variance analysis and estimated N_sites_, and Munc13 expression rank. **B-D)** The same as A, but for other synapses in the current imaging region. Note the variability in the goodness-of-fit for the uniform release probability binomial model. The synapse in panel **D** possesses excellent agreement with the binomial model.

**Figure S4.**
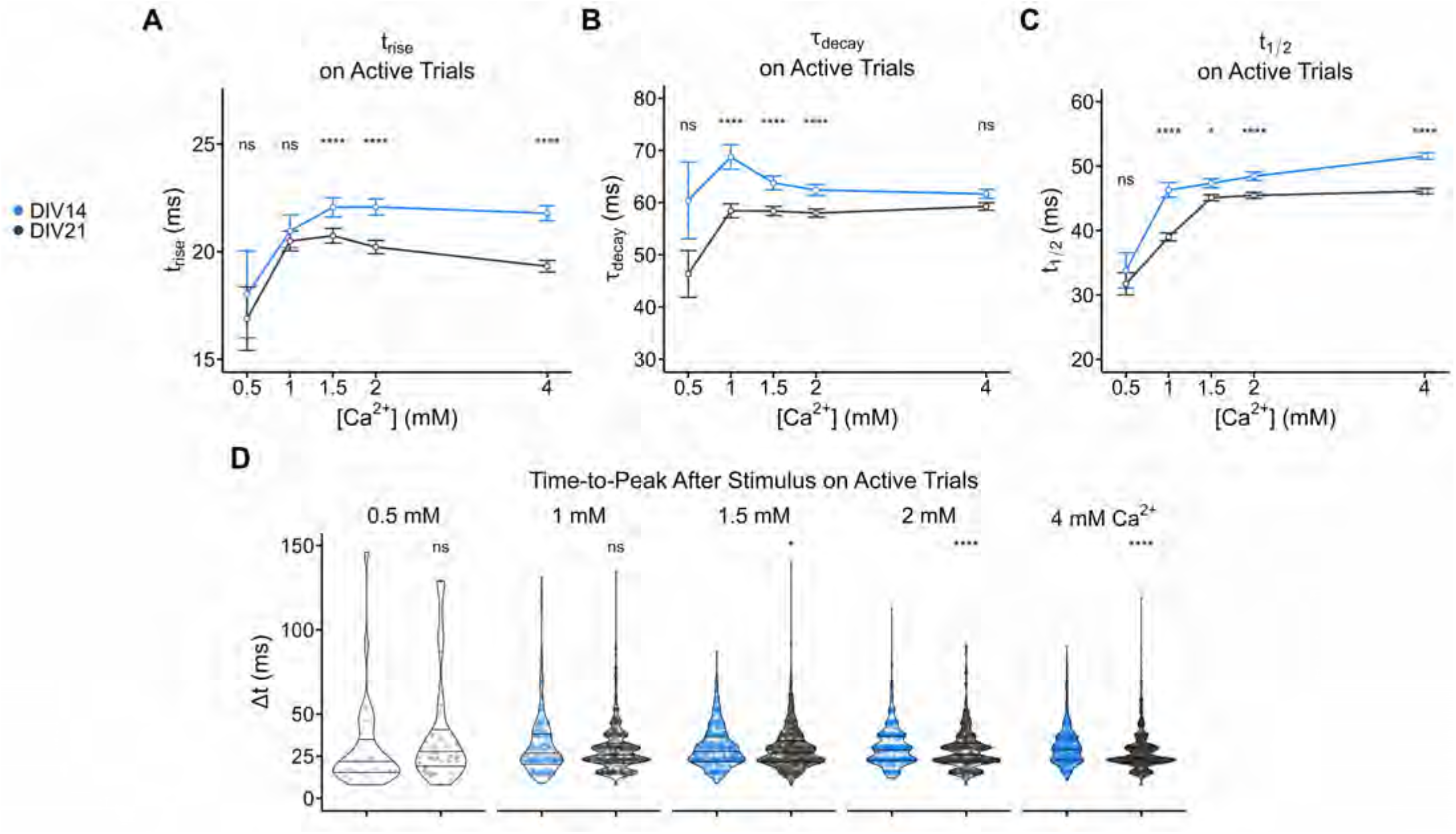
DIV14 synapses exhibit slower glutamate release and clearance kinetics than their DIV21 counterparts. **A)** t_rise_ (ms) on active single AP trials for DIV14 and DIV21 synapses. iGluSnFR3 transients had longer rise times for DIV14 synapses at all [Ca^2+^]_bath_ tested. B) τ_decay_ (ms) on active single AP trials for DIV14 and DIV21 synapses. iGluSnFR3 transients decayed more slowly back to baseline in DIV14 synapses, suggesting that glutamate clearance post-stimulus is more rapid in DIV21 synapses. **C)** Full-width at half-maximum (t_1/2_, ms) on active single AP trials for DIV14 and DIV21 synapses. iGluSnFR3 transients stimulated by single APs were longer in duration at DIV14 synapses than at DIV21 synapses, consistent with slower rise and decay kinetics at DIV14. **D)** Violin plots of time-to-peak after single AP stimulus (Δt, ms) for DIV14 and DIV21 synapses. Note that this data is identical to Figure 5F, but presented here as a violin plot to show the distribution at each [Ca^2+^]_bath_. Here, one can readily observe the quantization of the data driven by our imaging framerate (∼7.8 ms per frame, ∼128 Hz). As such, the distribution of measured Δt has multiple modes. Still, iGluSnFR3 transients more rapidly reached their peak fluorescence in DIV21 synapses, suggesting kinetic differences between DIV14 and DIV21 synapses that are partly explained by differences in Munc13 expression.

**Figure S5.**
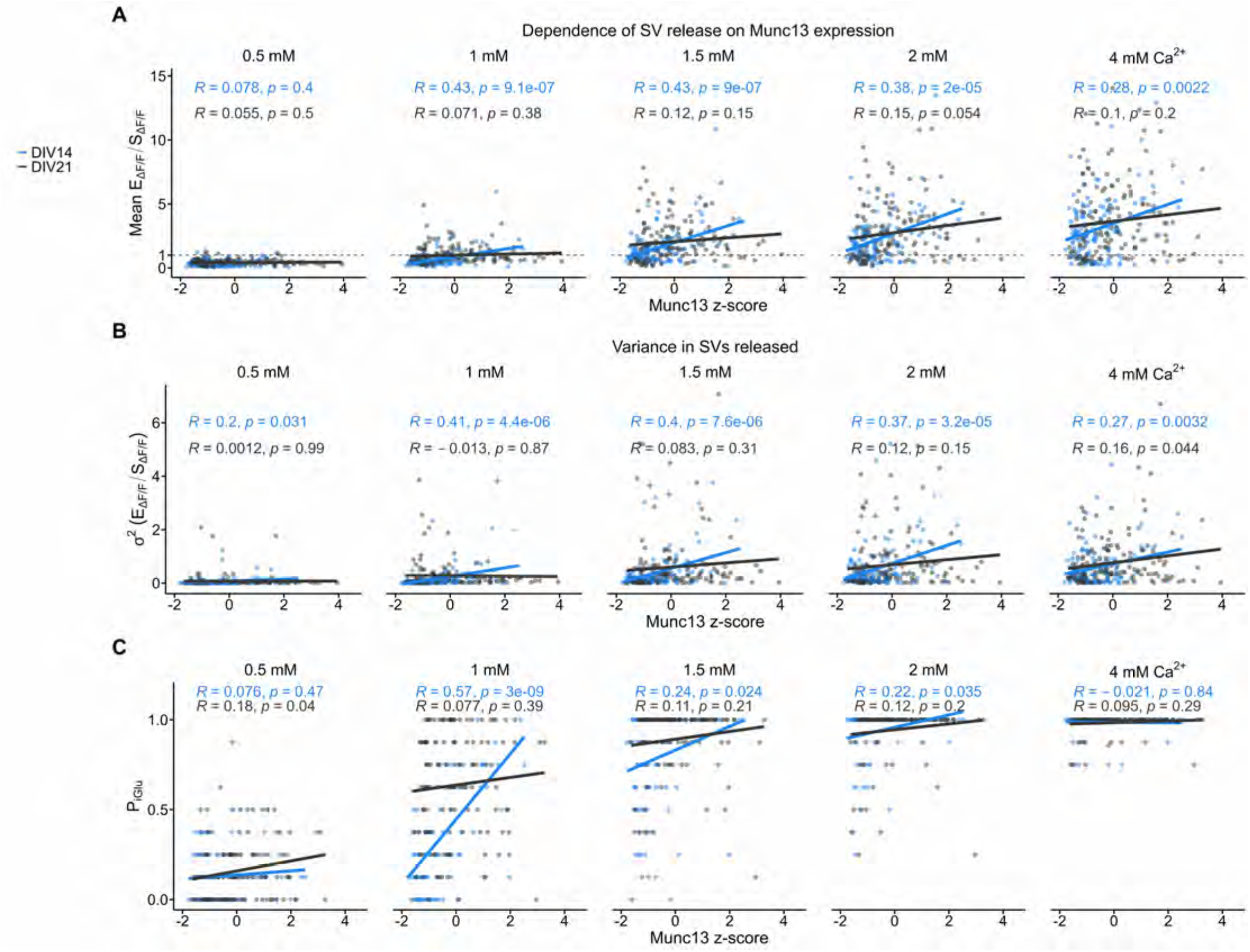
Munc13 expression strongly predicts total SV release per AP, variance in SVs released, and probability of observing a glutamate transient (P_iGlu_) at DIV14 synapses. **A)** Average iGluSnFR3 transient amplitude in terms of (E_ΔF/F_/S_ΔF/F_) to estimate the number of SVs released per AP vs. Munc13 expression (z-score) for DIV14 and DIV21 synapses across [Ca^2+^]_bath_. Each point is a single synapse. Dashed line indicates the iGluSnFR3 response to a single SV. Shown are the R values and statistical significance of the Pearson correlation. **B)** Variance in SVs released (σ^2^, E_ΔF/F_/S_ΔF/F_) vs. Munc13 expression (z-score) for DIV14 and DIV21 synapses across [Ca^2+^]_bath_. Shown are the R values and statistical significance of the Pearson correlation. **C)** Probability of observing an iGluSnFR3 transient per single AP trial vs. Munc13 expression (z-score) for DIV14 and DIV21 synapses across [Ca^2+^]_bath_. Shown are the R values and statistical significance of the Pearson correlation. Note that the data is highly discretized because each synapse possessed 8 trials at each [Ca^2+^]_bath_.

**Figure S6.**
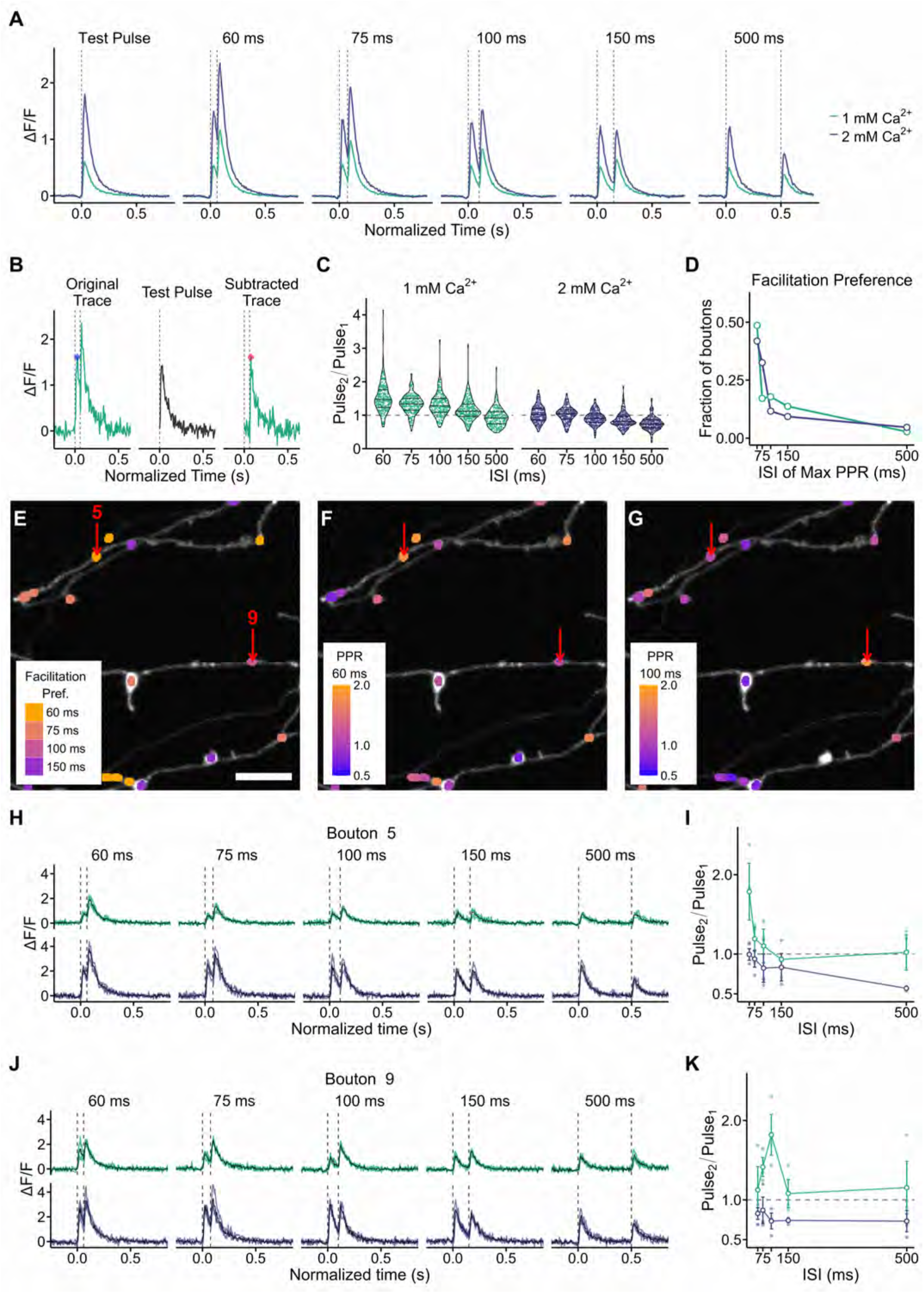
iGluSnFR3 responses evoked by paired stimuli reveal heterogeneous short-term plasticity dynamics along the axonal arbor (Synaptophysin-mRuby segmentation). **A)** The averaged iGluSnFR3 response for each stimulus protocol administered in this experiment across [Ca^2+^]_bath_ = 1, 2 mM. Dashed vertical lines indicate the stimulus. **B)** Subtraction routine for determining paired-pulse ratios (PPRs) at individual boutons. *Left:* The amplitude of Pulse_1_ (blue diamond) was the maximum ΔF/F observed within the first stimulus epoch in the unmodified, Original Trace for each trial. *Middle:* For each bouton, we determined the average iGluSnFR3 response to a single stimulus (Test Pulse) from three trials. *Right:* The amplitude of Pulse_2_ (red diamond) was the maximum ΔF/F observed within the second stimulus epoch after subtracting the Test Pulse from the Original Trace for each trial. **C)** Violin plots showing the value of PPR as a function of [Ca^2+^]_bath_, interstimulus interval (ISI). Each colored dot indicates the mean value of PPR for each bouton. The dashed line indicates PPR = 1, the threshold between facilitation (PPR > 1) and depression (PPR < 1). Boutons largely depressed at all ISIs at [Ca^2+^]_bath_ = 2 mM. **D)** We categorized boutons according to which ISI produced the maximum value of PPR for boutons, the “Facilitation Bias”. Of the facilitating boutons (PPR ≥ 1.1), 48.6% facilitated most strongly at ISI = 60 ms and [Ca^2+^]_bath_ = 1 mM. E) Facilitation bias mapped to an exemplar axonal arbor for [Ca^2+^]_bath_ = 1 mM. In this imaging region, no boutons facilitated most strongly at ISI = 500 ms. Scale bar = 5 µm. F,G) PPR mapped to the same axonal arbor as in **E**. Warmer colors indicate facilitation, while cooler colors indicate depression. **F)** PPRs for ISI = 60 ms, [Ca^2+^]_bath_ = 1 mM. G) PPRs for ISI = 100 ms, [Ca^2+^]_bath_ = 1 mM. In **E,F,G,** red arrows indicate boutons targeted for functional analysis. **H, I)** Bouton report for Bouton #5. H) The individual trial responses are color-coded by [Ca^2+^]_bath_ with the averaged response in black. I) PPR curves across ISIs for 1 and 2 mM Ca^2+^ for Bouton #5. **J,K)** Bouton report for Bouton #9 as in **H,I..**

**Figure S7.**
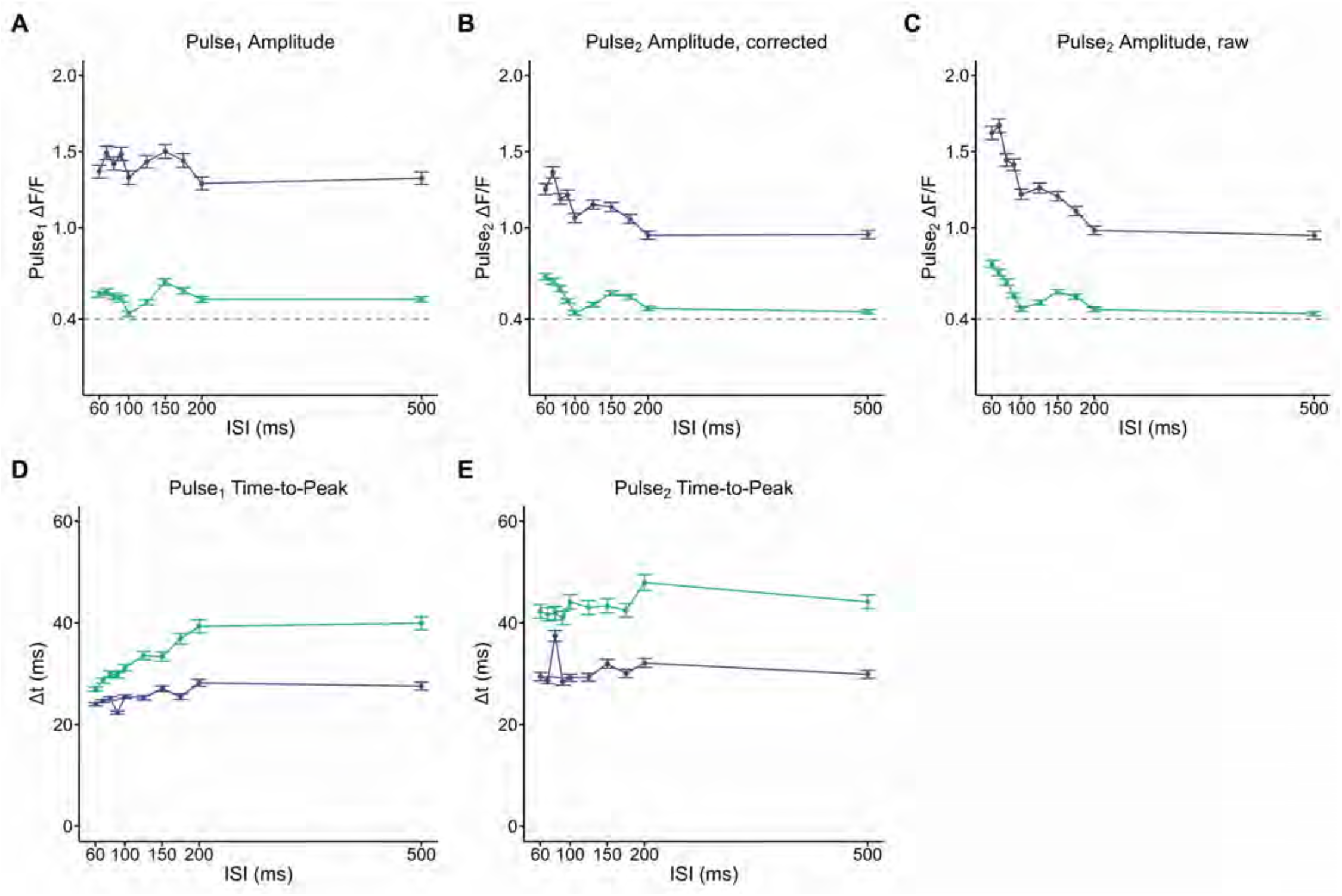
Basic peak properties for paired-pulse experiment from main-text Figure 6. **A)** Pulse_1_ amplitudes as a function of ISI. There is little jitter across ISI, consistent with a stable initial *P_r_* across the experiment. Dashed line indicates the amplitude of the average iGluSnFR3 response to a putative single SV. **B)** Pulse_2_ amplitudes as a function of ISI, corrected by subtracting the average iGluSnFR3 response to a single action potential for each synapse from its corresponding paired-pulse traces. The shape of the Pulse_2_ amplitude curve implies short-term plasticity mechanisms are engaged in a frequency-dependent manner. **C)** Uncorrected Pulse_2_ amplitudes as a function of ISI. Note that compared to *B*, Pulse_2_ amplitudes at longer ISIs are unaffected. **D)** Pulse_1_ time-to-peak. Consistent with observations from main-text Figure 5, iGluSnFR3 transients peak more rapidly following stimulus at 2 mM Ca^2+^ than at 1 mM Ca^2+^. Note that the shorter time-to-peak at short ISIs is likely an artifact of the way iGluSnFR3 were identified (e.g., for ISI = 60 ms, the maximum possible Δt is 60 ms, while for ISI ≥ 200 ms, the maximum possible Δt is 200 ms). **E)** Pulse_2_ time-to-peak. Consistent with observations from main-text Figure 5, iGluSnFR3 transients peak more rapidly following stimulus at 2 mM Ca^2+^ than at 1 mM Ca^2+^.

**Figure S8.**
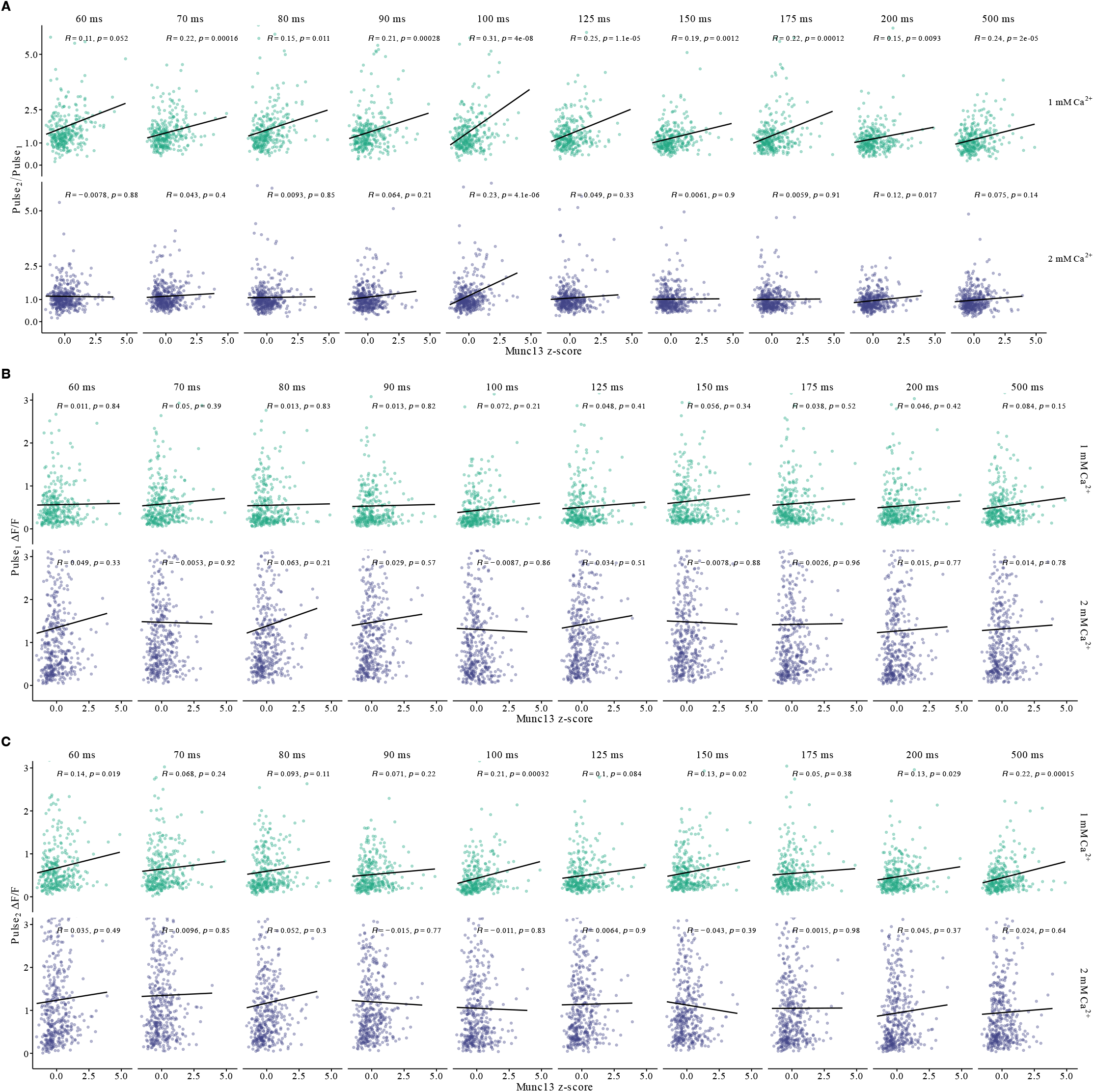
Dependence of short-term plasticity dynamics on synaptic Munc13 expression for all ISIs. **A)** At 1 mM Ca^2+^, PPR positively correlates with Munc13 expression for all ISIs, with strength of Pearson correlation (R value) and statistical significance (p value) varying according to ISI. At 2 mM Ca^2+^, PPR does not correlate with Munc13 expression with ISI = 100 ms as the notable exception. **B)** The amplitude (ΔF/F) of the iGluSnFR3 response to the first stimulus showed little to dependence on Munc13 expression at either [Ca^2+^]_bath_ tested. **C)** The amplitude (ΔF/F) of the iGluSnFR3 response to the 2^nd^ stimulus showed little to no dependence on Munc13 expression at either [Ca^2+^]_bath_ tested.

**Figure S9.**
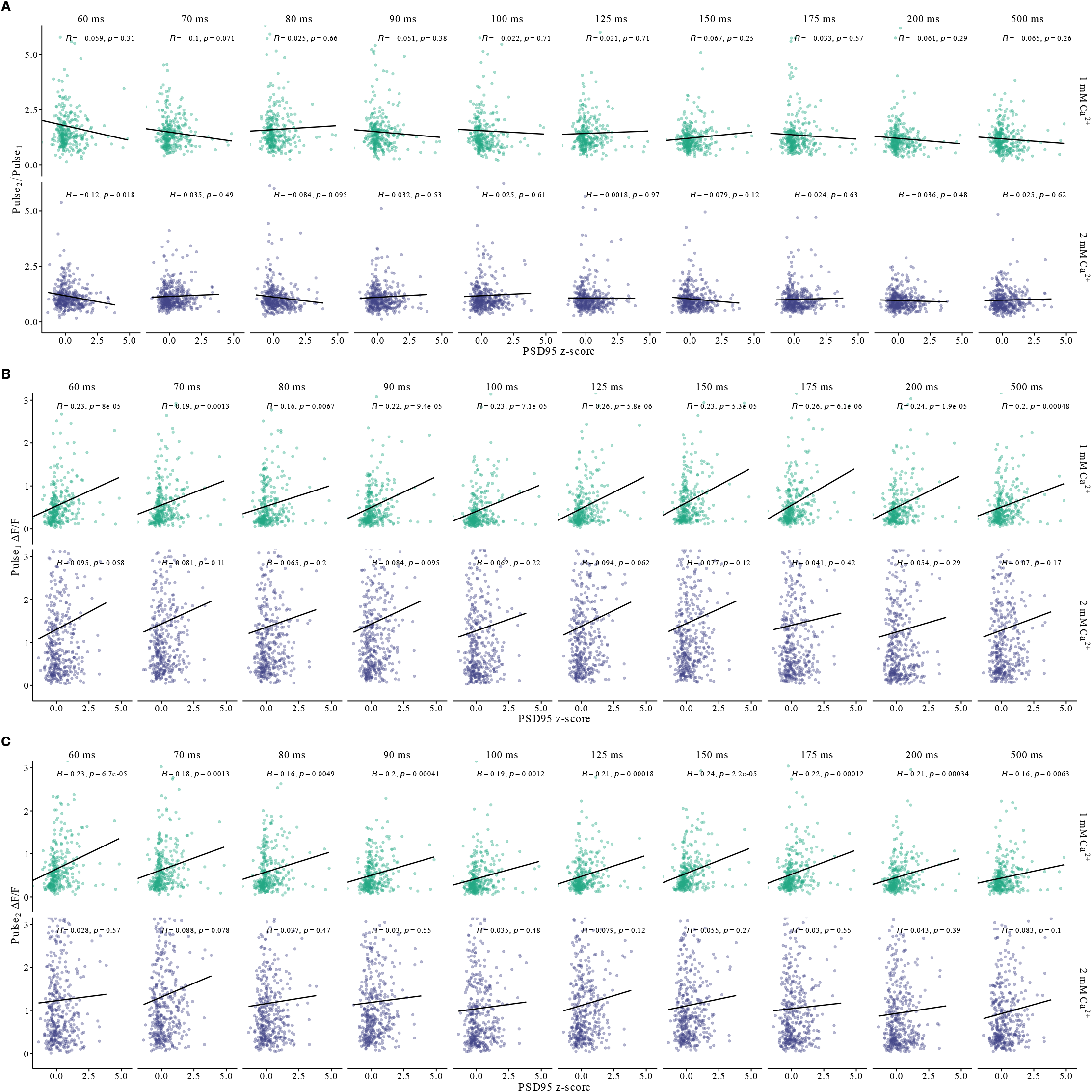
Dependence of short-term plasticity dynamics on PSD95.FingR.TagRFP expression for all ISIs. **A)** PPR did not correlate with PSD95.FingR.TagRFP expression for any ISI at either [Ca^2+^]_bath_ tested, with strength of Pearson correlation (R value) and statistical significance (p value) varying according to ISI. **B)** The amplitude (ΔF/F) of the iGluSnFR3 response to the first stimulus showed a dependence on PSD95.FingR.TagRFP expression at both [Ca^2+^]_bath_ tested. **C)** The amplitude (ΔF/F) of the iGluSnFR3 response to the 2^nd^ stimulus showed substantial dependence on PSD95.FingR.TagRFP expression, but only at 1 mM Ca^2+^.

**Figure S10.**
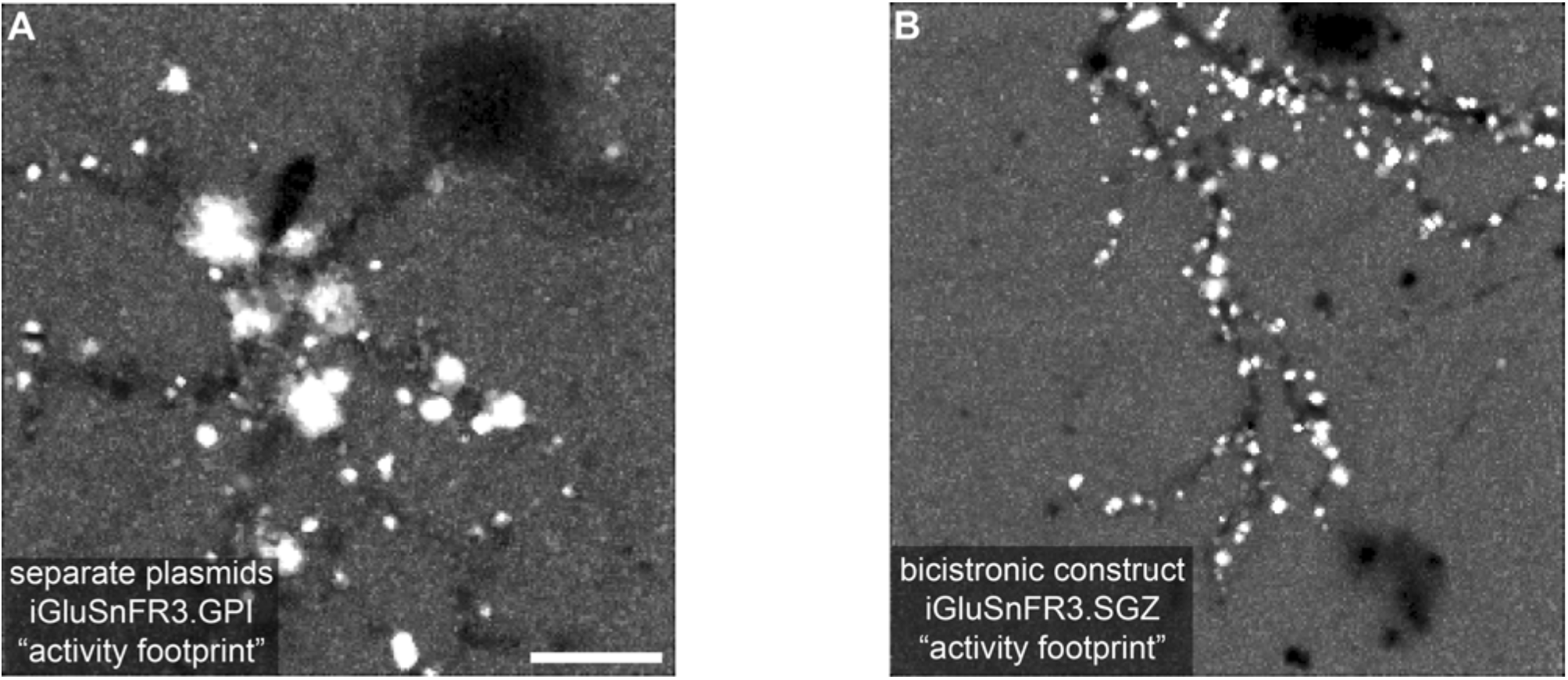
Improved trafficking of iGluSnFR3.SGZ into the synapse improves activity-segmentation of single spines versus iGluSnFR3.GPI. **A)** Activity-based segmentation of iGluSnFR3 signal collected from a neuron expressing spine-HaloTag and the GPI-anchored version of iGluSnFR3 (iGluSnFR3.GPI) from separate plasmids. While some individual spines are extracted for analysis, GPI-anchored iGluSnFR3 trafficks poorly into the synapse, limiting synaptic resolution and decreasing throughput. Scale bar = 20 µm. **B)** Activity-based segmentation of the stargazin(SGZ)-anchored version of iGluSnFR3 signal collected from a neuron expressing a single, bicistronic construct: spine-HaloTag-P2A-iGluSnFR3.SGZ. SGZ-anchored iGluSnFR3 avidly trafficks into the synapse, increasing synaptic resolution and throughput.

**Figure S11.**
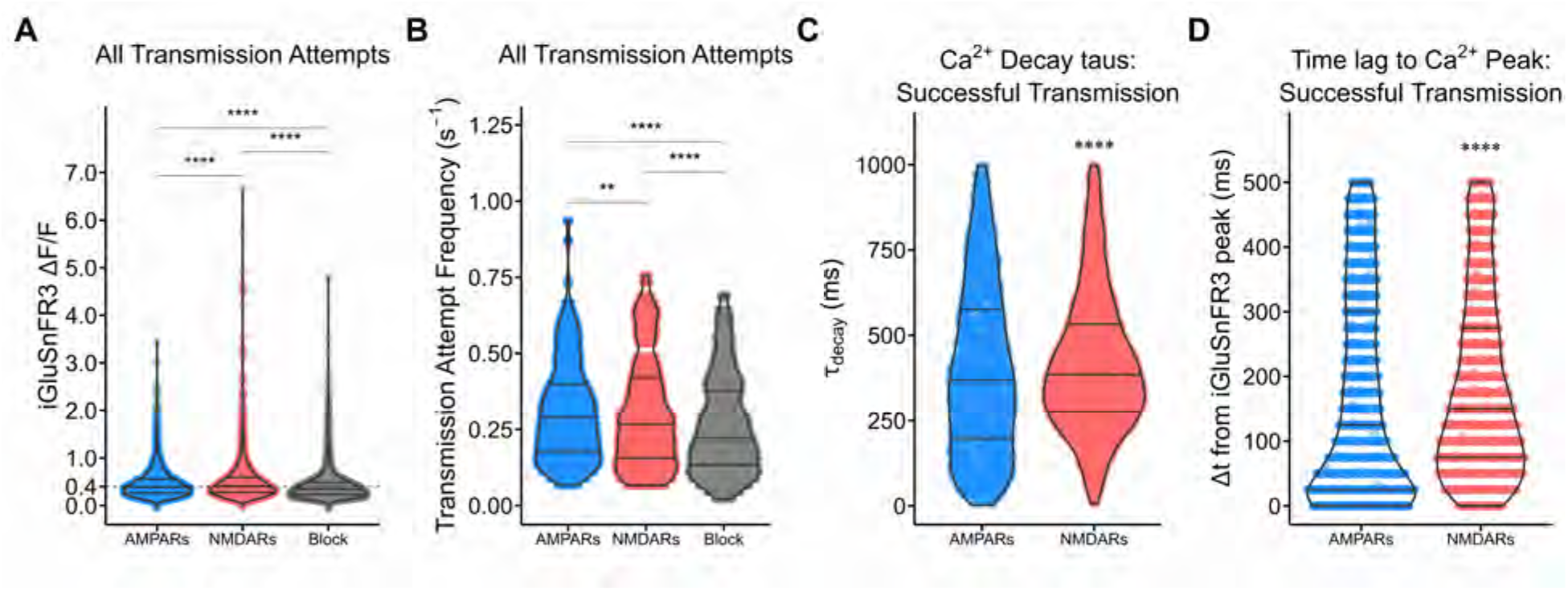
Statistical properties of all-optical synaptic transmission recordings (main-text Figure 8, 9). **A)** iGluSnFR3 amplitude across all transmission attempts. Main-text Figure 8 shows that iGluSnFR3 amplitudes for successful synaptic transmission events were statistically indistinguishable between the “AMPARs open” and “NMDARs open” stages of the experiment. However, when all synaptic transmission attempts are considered (inclusive of failures and successes), we do resolve differences in iGluSnFR3 amplitude. It remains unclear why iGluSnFR3 event amplitude increases slightly following APV washout (note the shift in the top quartile), and then decreases following iGluR blockade. **B)** Transmission attempt frequency by imaging phase. While some synapses appear to increase their transmission frequency following APV washout (red), most synapses experienced a rundown in spontaneous release frequency over the course of the experiment. **C)** Ca^2+^ event τ_decay_ from single-exponential decay fitting among successful transmission events. **D)** Time delay to Ca^2+^ peak from preceding iGluSnFR3 event among successful transmission events. On average, AMPARs quickly reached their peak amplitude following iGluSnFR3 events, indicating fast receptor kinetics upon successful receptor activation. By contrast, NMDARs took longer to peak, possibly due to saturation of JF_646_-BAPTA dye or more permissive receptor activation dynamics due to their higher affinity for glutamate. Note that the quantization of the Δt values is an inherent property of our sampling frequency (∼40 Hz).

## Notes

### Competing Interest Statement

The authors have declared no competing interest.

### Summary of Updates

Substantial new experiments and analysis have been added, including four new main figures and several additional supporting figures. Improvements to the dual-sensor imaging and analysis now allow direct and novel discrimination of AMPAR- and NMDAR-mediated transmission at single synapses mapped across the dendritic arbor. A new complete workflow allows matching post hoc immunocytochemistry to live-imaged synapses for examination of molecular mechanisms underlying synaptic behavior, plasticity, and population-level variance.

