## Supplemental figures for "Dissecting the functional heterogeneity of glutamatergic synapses with high-throughput optical physiology"

Table of Contents:

*Figure S1*. Additional functional properties of the three bouton classes from main-text Figure 4 imply differences in the SV release machinery.

*Figure S2*. Boutons separated by ~1.8 μm exhibit divergent iGluSnFR3 behavior.

*Figure S3*. Single-synapse reports for selected DIV14 synapses (main-text Figure 5).

*Figure S4*. DIV14 synapses exhibit slower glutamate release and clearance kinetics than their DIV21 counterparts.

*Figure S5*. Munc13 expression predicts total SV release per AP, variance in SVs released, and probability of observing a glutamate transient ($P_{iGlu}$) with correlation strength depending on synapse age.

Figure *S6*. iGluSnFR3 responses evoked by paired stimuli reveal heterogeneous short-term plasticity dynamics along the axonal arbor (Synaptophysin-mRuby segmentation).

*Figure S7*. Basic peak properties for paired-pulse experiment from main-text Figure 6.

*Figure S8*. Dependence of short-term plasticity dynamics on synaptic Munc13 expression for all ISIs.

*Figure S9*. Dependence of short-term plasticity dynamics on PSD95.FingR.TagRFP expression for all ISIs.

*Figure S10*. Increased trafficking of iGluSnFR3.SGZ into the synapse improves activity-segmentation of single spines versus iGluSnFR3.GPI.

*Figure S11*. Statistical properties of all-optical synaptic transmission recordings (main-text Figure 8,9).

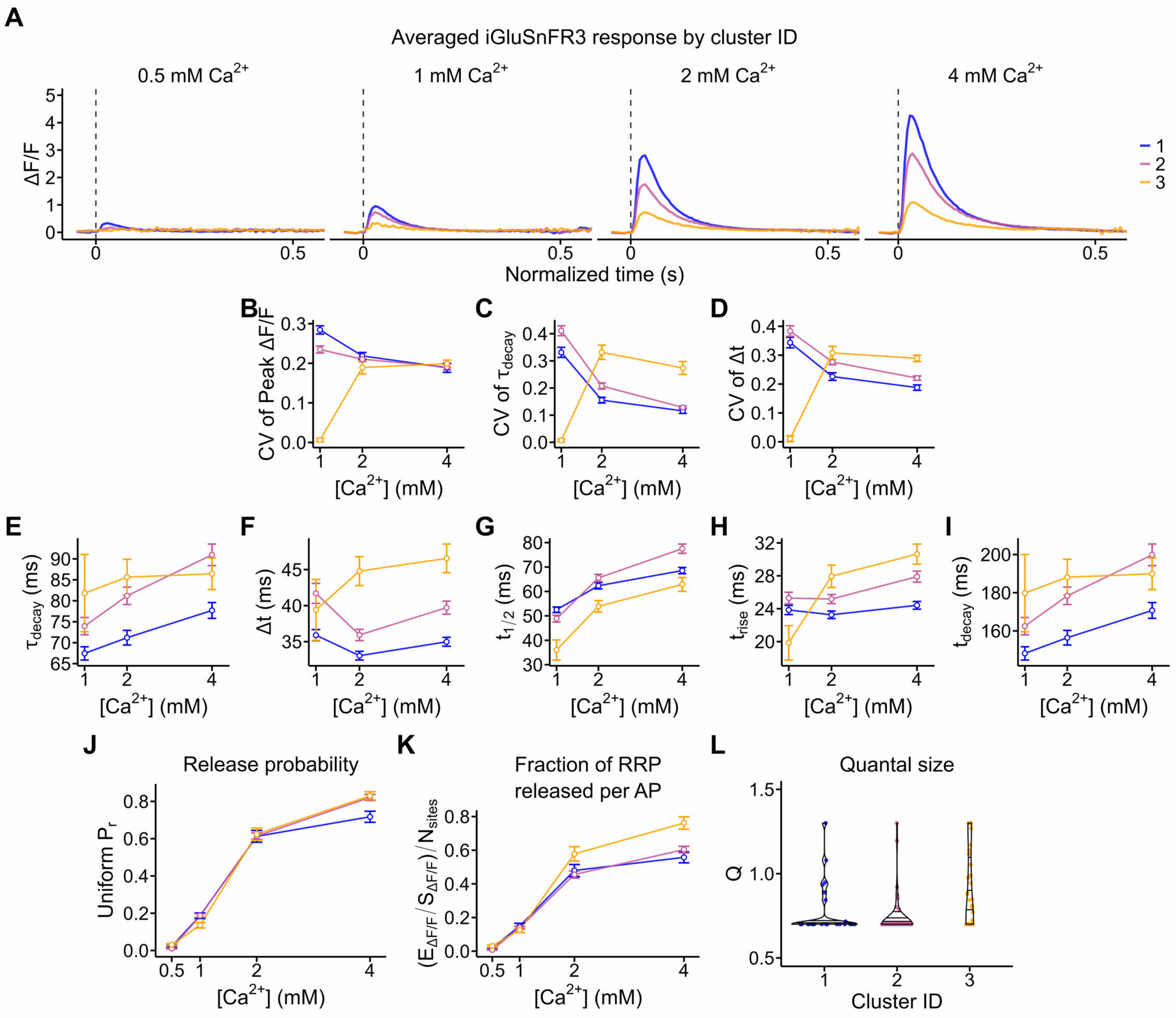


**Figure S1. Additional functional properties of the three bouton classes from main-text Figure 4 imply differences in the SV release machinery. A)** The average iGluSnFR3 response in response to stimulus across $[Ca^{2+}]_{bath}$ and cluster ID. For the plots presented in **B-G**, 0.5 mM $Ca^{2+}$ has been omitted due to the sparse data in cluster 2 and 3. **B)** Coefficient of variation (CV) of peak ΔF/F vs. $[Ca^{2+}]_{bath}$. **C)** CV of $\tau_{decay}$ vs. $[Ca^{2+}]_{bath}$. **D)** CV of Δt vs. $[Ca^{2+}]_{bath}$. **E)** $\tau_{decay}$ vs. $[Ca^{2+}]_{bath}$. **F)** Δt vs. $[Ca^{2+}]_{bath}$. **G)** $t_{1/2}$ vs. $[Ca^{2+}]_{bath}$. **H)** $t_{rise}$ (10-90%) vs. $[Ca^{2+}]_{bath}$. **I)** $t_{decay}$ (90-10%) vs. $[Ca^{2+}]_{bath}$. **J)** Uniform release probability, $P_r$ vs. $[Ca^{2+}]_{bath}$. **K)** RRP fraction released per AP $(E_{\Delta F/F}/S_{\Delta F/F})/N_{sites}$ vs. $[Ca^{2+}]_{bath}$. **L)** Binomial model Q vs. cluster ID.

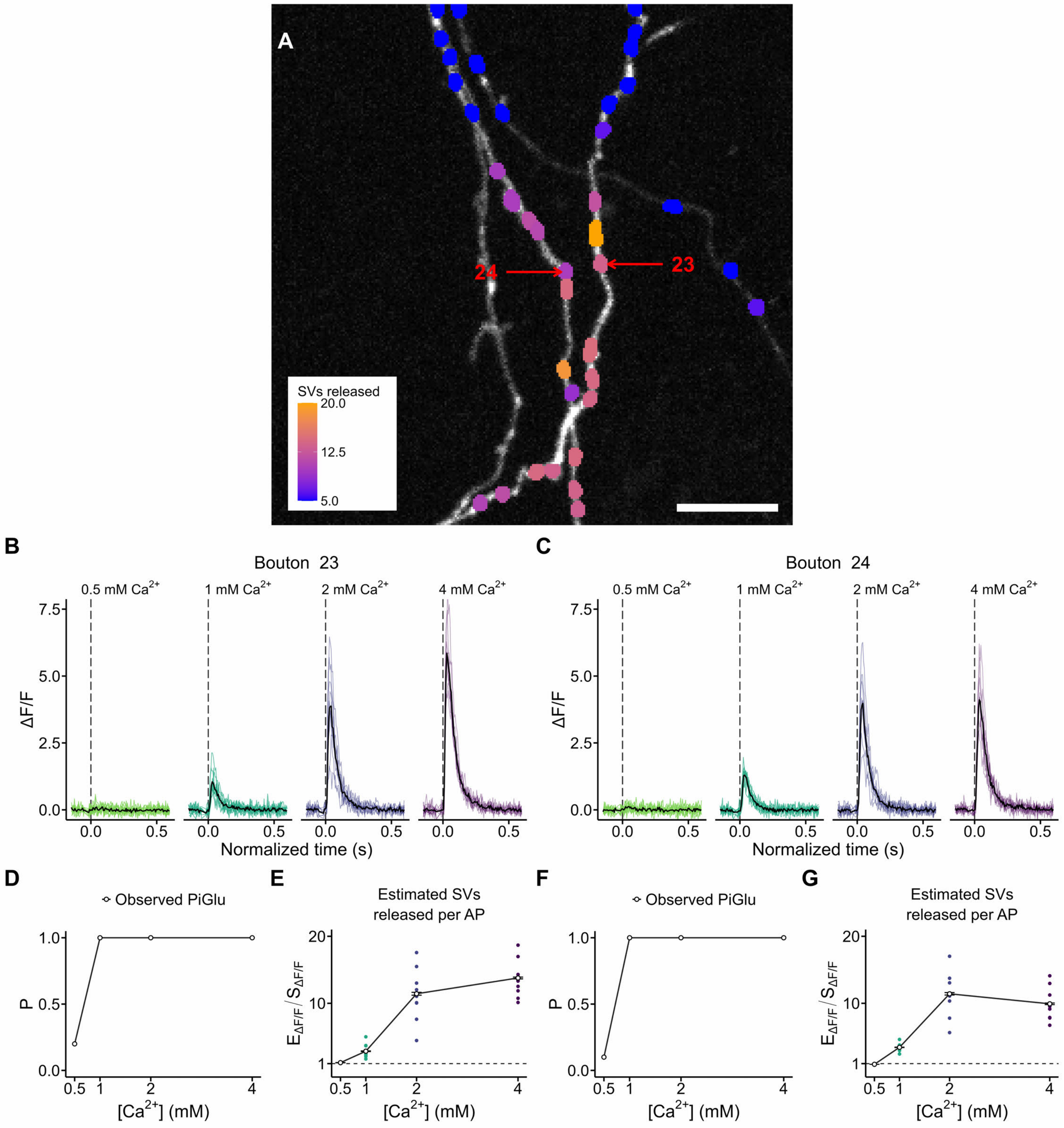


**Figure S2. Boutons separated by ~1.8 μm exhibit divergent iGluSnFR3 behavior. A)** The imaging region from **Figure 4I**, color-coded according to number of SVs released at 4 mM $Ca^{2+}$. Boutons 23 and 24 are indicated for further analysis. Scale bar = 5 μm. **B, C)** The iGluSnFR3 responses at all $[Ca^{2+}]_{bath}$ tested. Though the boutons are only separated by 1.8 μm on separate branches of the same axon, they exhibit divergent iGluSnFR3 behavior. **D,F)** $P_{iGlu}$ vs. $[Ca^{2+}]_{bath}$. **E,G)** Estimated SVs released per AP ($E_{\Delta F/F}$/ $S_{\Delta F/F}$) vs. $[Ca^{2+}]_{bath}$. Note that Bouton 24 appears to reaches its maximal response at 2 mM $Ca^{2+}$, with little change in behavior at 4 mM $Ca^{2+}$.

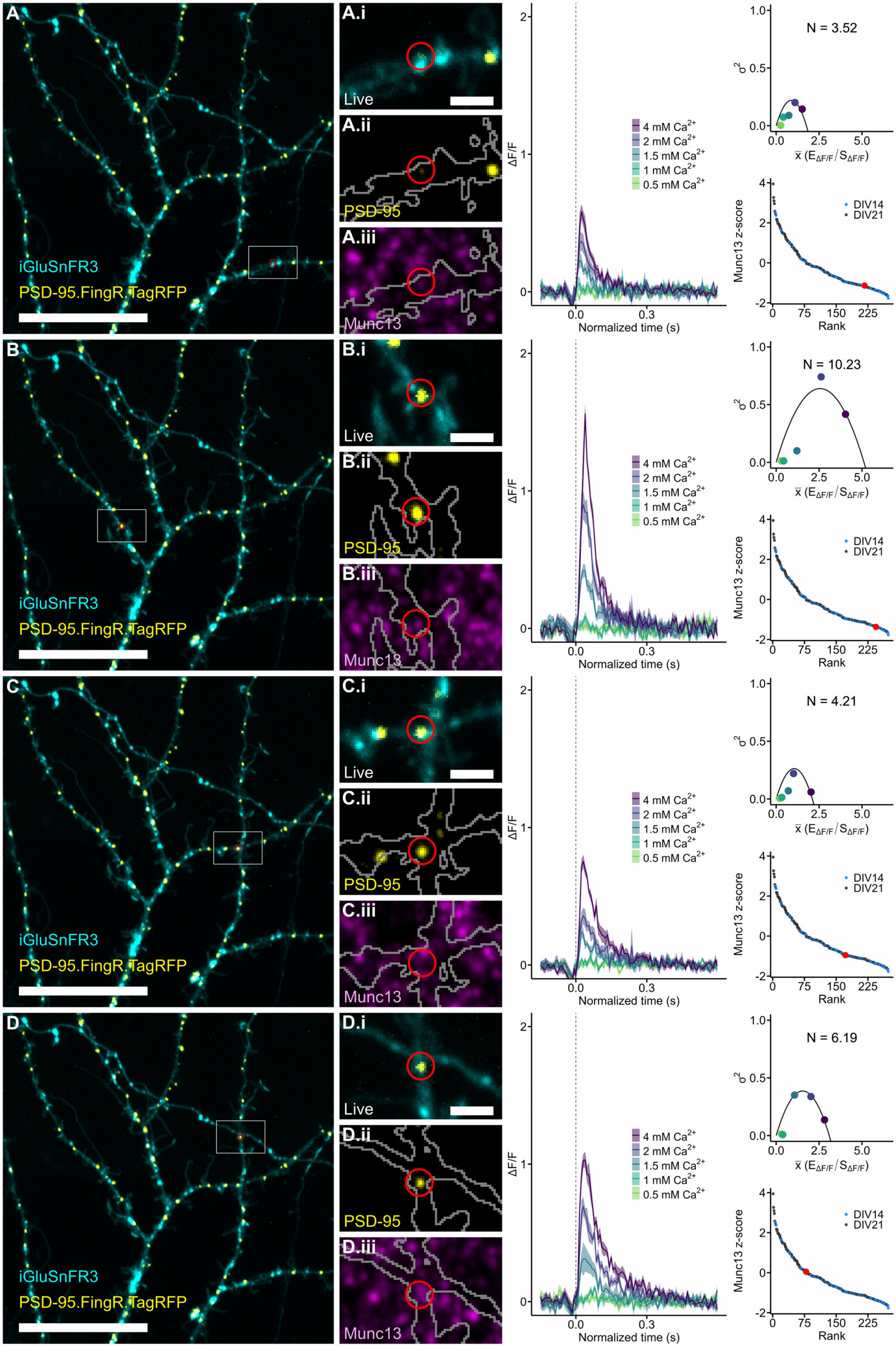

A
iGluSnFR3
PSD-95.FingR.TagRFP
A.i
Live
A.ii
PSD-95
A.iii
Munc13
4 mM Ca2+
2 mM Ca2+
1.5 mM Ca2+
1 mM Ca2+
0.5 mM Ca2+
ΔF/F
Normalized time (s)
N = 3.52
σ2
x̄ (EΔF/F/SΔF/F)
Munc13 z-score
Rank
DIV14
DIV21
B
B.i
B.ii
B.iii
N = 10.23
C
C.i
C.ii
C.iii
N = 4.21
D
D.i
D.ii
D.iii
N = 6.19

**Figure S3. Single-synapse reports for selected DIV14 synapses (main-text Figure 5). A)** Full imaging region of DIV14 dendrite expressing iGluSnFR3 (cyan) and PSD-95.FingR.TagRFP (yellow). Scale bar = 20 μm. **(i)** Inset of the imaging region shown in **A**. Red circle indicates the synapse selected for analysis. Scale bar = 2 μm. **(ii)** Post-hoc imaging of PSD-95.FingR.TagRFP at higher illumination intensity. Grey outline indicates a mask of the live iGluSnFR3 signal. **(iii)** Post-hoc imaging of Munc13 immunofluorescence. **Right**, the single synapse iGluSnFR3 response to single APs at each $[Ca^{2+}]_{bath}$ tested, mean-variance analysis and estimated $N_{sites}$, and Munc13 expression rank. **B-D)** The same as A, but for other synapses in the current imaging region. Note the variability in the goodness-of-fit for the uniform release probability binomial model. The synapse in panel **D** possesses excellent agreement with the binomial model.

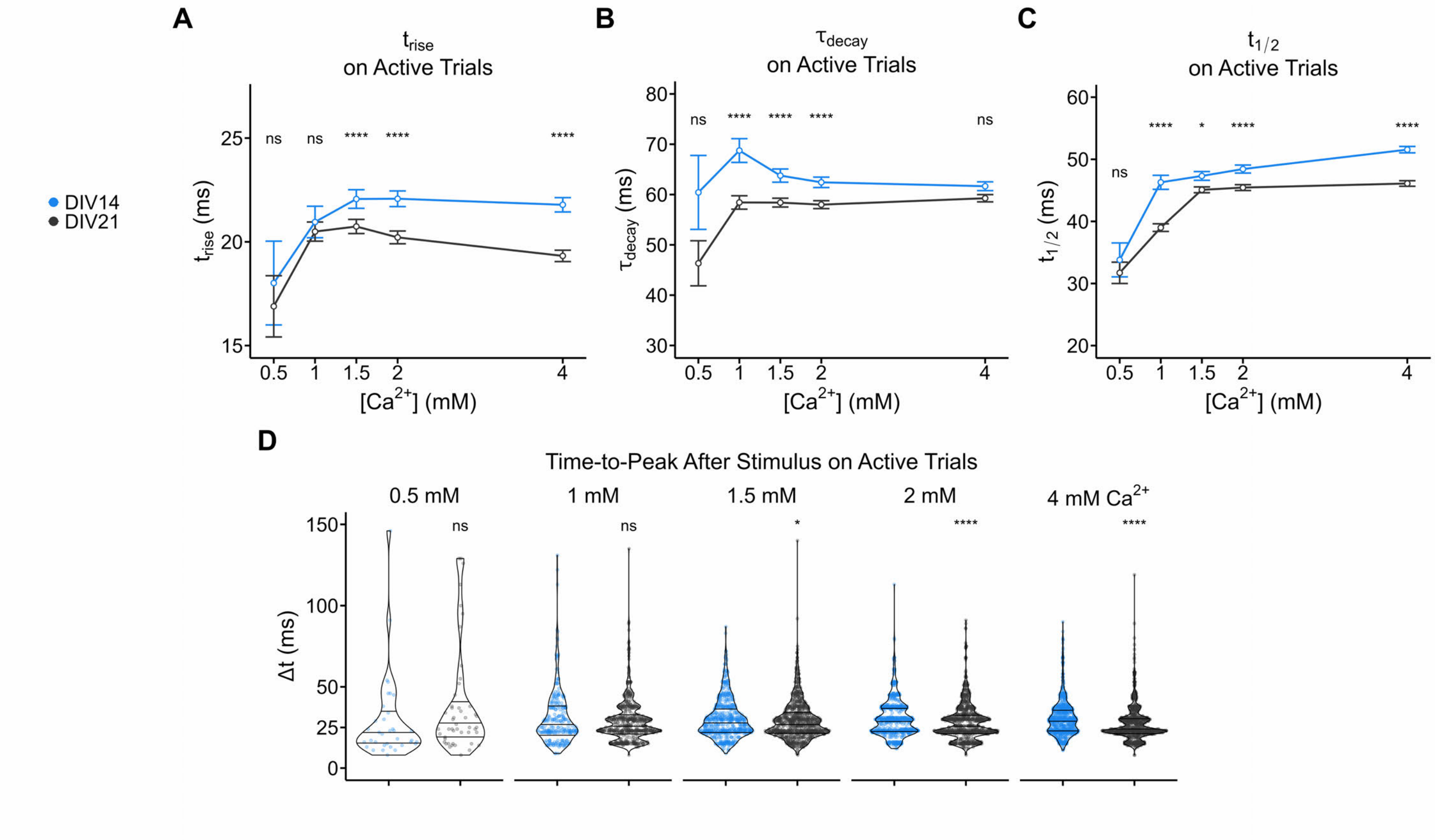


**Figure S4. DIV14 synapses exhibit slower glutamate release and clearance kinetics than their DIV21 counterparts. A)** $t_{rise}$ (ms) on active single AP trials for DIV14 and DIV21 synapses. iGluSnFR3 transients had longer rise times for DIV14 synapses at all $[Ca^{2+}]_{bath}$ tested. B) $\tau_{decay}$ (ms) on active single AP trials for DIV14 and DIV21 synapses. iGluSnFR3 transients decayed more slowly back to baseline in DIV14 synapses, suggesting that glutamate clearance post-stimulus is more rapid in DIV21 synapses. **C)** Full-width at half-maximum ($t_{1/2}$, ms) on active single AP trials for DIV14 and DIV21 synapses. iGluSnFR3 transients stimulated by single APs were longer in duration at DIV14 synapses than at DIV21 synapses, consistent with slower rise and decay kinetics at DIV14. **D)** Violin plots of time-to-peak after single AP stimulus (Δt, ms) for DIV14 and DIV21 synapses. Note that this data is identical to **Figure 5F**, but presented here as a violin plot to show the distribution at each $[Ca^{2+}]_{bath}$. Here, one can readily observe the quantization of the data driven by our imaging framerate (~7.8 ms per frame, ~128 Hz). As such, the distribution of measured Δt has multiple modes. Still, iGluSnFR3 transients more rapidly reached their peak fluorescence in DIV21 synapses, suggesting kinetic differences between DIV14 and DIV21 synapses that are partly explained by differences in Munc13 expression.

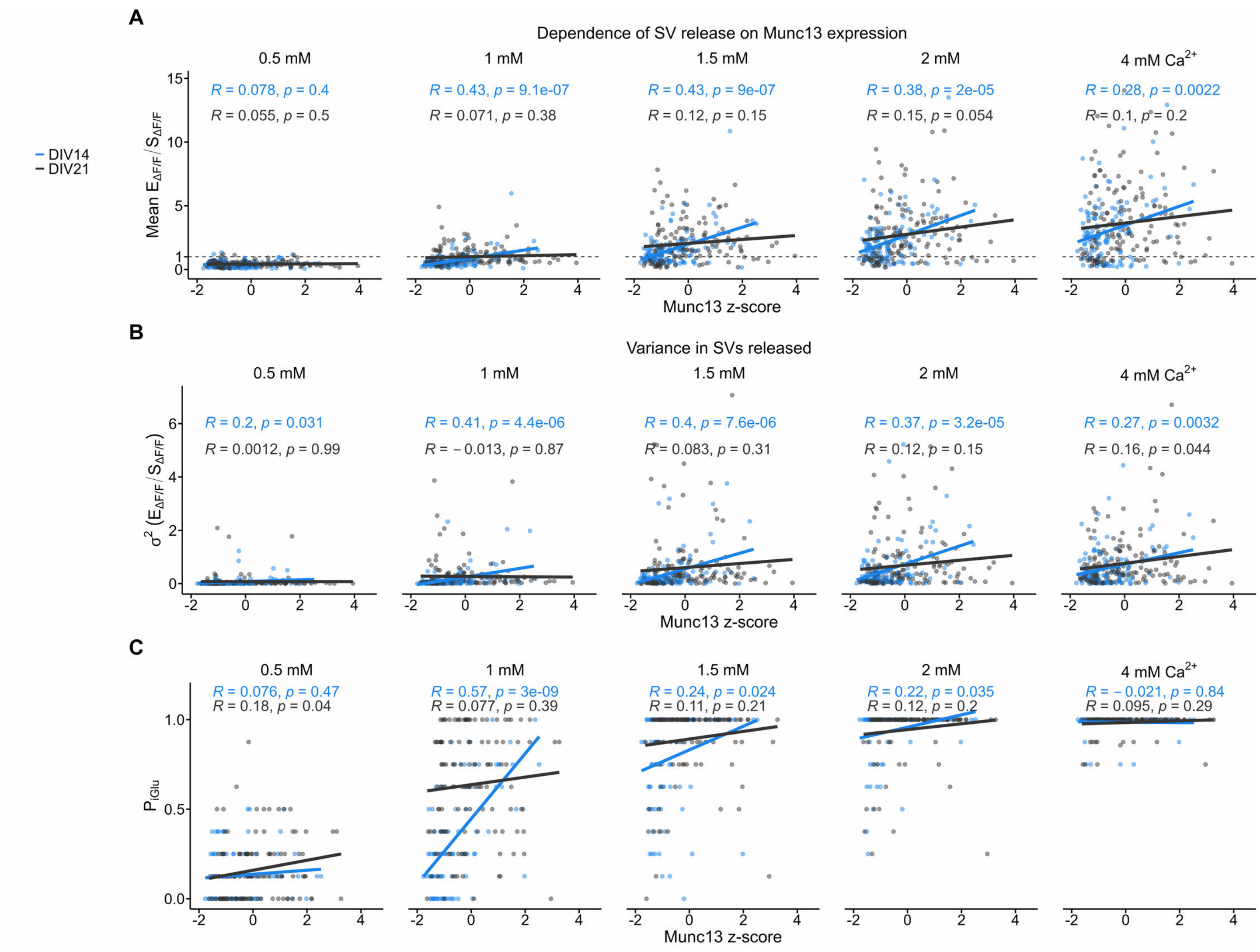


**Figure S5. Munc13 expression strongly predicts total SV release per AP, variance in SVs released, and probability of observing a glutamate transient ($P_{iGlu}$) at DIV14 synapses. A)** Average iGluSnFR3 transient amplitude in terms of ($E_{\Delta F/F}/S_{\Delta F/F}$) to estimate the number of SVs released per AP vs. Munc13 expression (z-score) for DIV14 and DIV21 synapses across $[Ca^{2+}]_{bath}$. Each point is a single synapse. Dashed line indicates the iGluSnFR3 response to a single SV. Shown are the R values and statistical significance of the Pearson correlation. **B)** Variance in SVs released ($\sigma^2$, $E_{\Delta F/F}/S_{\Delta F/F}$) vs. Munc13 expression (z-score) for DIV14 and DIV21 synapses across $[Ca^{2+}]_{bath}$. Shown are the R values and statistical significance of the Pearson correlation. **C)** Probability of observing an iGluSnFR3 transient per single AP trial vs. Munc13 expression (z-score) for DIV14 and DIV21 synapses across $[Ca^{2+}]_{bath}$. Shown are the R values and statistical significance of the Pearson correlation. Note that the data is highly discretized because each synapse possessed 8 trials at each $[Ca^{2+}]_{bath}$.

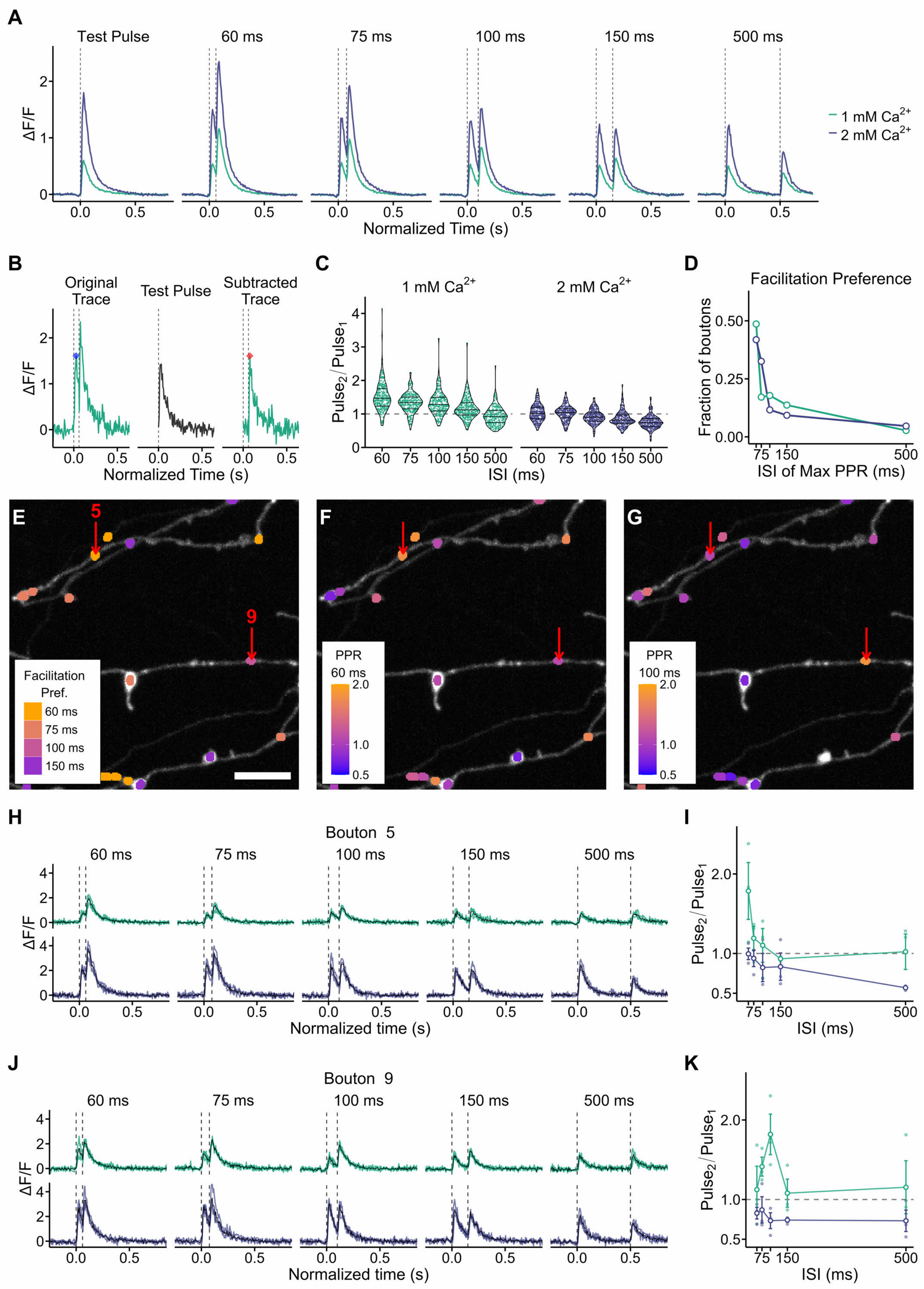
A
Test Pulse
60 ms
75 ms
100 ms
150 ms
500 ms
ΔF/F
Normalized Time (s)
1 mM Ca2+
2 mM Ca2+
B
Original Trace
Test Pulse
Subtracted Trace
ΔF/F
Normalized Time (s)
C
1 mM Ca2+
2 mM Ca2+
Pulse2/Pulse1
ISI (ms)
D
Facilitation Preference
Fraction of boutons
ISI of Max PPR (ms)
E
5
9
Facilitation Pref.
60 ms
75 ms
100 ms
150 ms
F
PPR 60 ms
G
PPR 100 ms
H
Bouton 5
60 ms
75 ms
100 ms
150 ms
500 ms
ΔF/F
Normalized time (s)
I
Pulse2/Pulse1
ISI (ms)
J
Bouton 9
60 ms
75 ms
100 ms
150 ms
500 ms
ΔF/F
Normalized time (s)
K
Pulse2/Pulse1
ISI (ms)

**Figure S6. iGluSnFR3 responses evoked by paired stimuli reveal heterogeneous short-term plasticity dynamics along the axonal arbor (Synaptophysin-mRuby segmentation). A)** The averaged iGluSnFR3 response for each stimulus protocol administered in this experiment across $[Ca^{2+}]_{bath}$ = 1, 2 mM. Dashed vertical lines indicate the stimulus. **B)** Subtraction routine for determining paired-pulse ratios (PPRs) at individual boutons. *Left:* The amplitude of $Pulse_1$ (blue diamond) was the maximum ΔF/F observed within the first stimulus epoch in the unmodified, Original Trace for each trial. *Middle:* For each bouton, we determined the average iGluSnFR3 response to a single stimulus (Test Pulse) from three trials. *Right:* The amplitude of $Pulse_2$ (red diamond) was the maximum ΔF/F observed within the second stimulus epoch after subtracting the Test Pulse from the Original Trace for each trial. **C)** Violin plots showing the value of PPR as a function of $[Ca^{2+}]_{bath}$, interstimulus interval (ISI). Each colored dot indicates the mean value of PPR for each bouton. The dashed line indicates PPR = 1, the threshold between facilitation (PPR > 1) and depression (PPR < 1). Boutons largely depressed at all ISIs at $[Ca^{2+}]_{bath}$ = 2 mM. **D)** We categorized boutons according to which ISI produced the maximum value of PPR for boutons, the "Facilitation Bias". Of the facilitating boutons (PPR ≥ 1.1), 48.6% facilitated most strongly at ISI = 60 ms and $[Ca^{2+}]_{bath}$ = 1 mM. **E)** Facilitation bias mapped to an exemplar axonal arbor for $[Ca^{2+}]_{bath}$ = 1 mM. In this imaging region, no boutons facilitated most strongly at ISI = 500 ms. Scale bar = 5 μm. **F,G)** PPR mapped to the same axonal arbor as in **E**. Warmer colors indicate facilitation, while cooler colors indicate depression. **F)** PPRs for ISI = 60 ms, $[Ca^{2+}]_{bath}$ = 1 mM. **G)** PPRs for ISI = 100 ms, $[Ca^{2+}]_{bath}$ = 1 mM. In **E**,**F**,**G**, red arrows indicate boutons targeted for functional analysis. **H, I)** Bouton report for Bouton #5. **H)** The individual trial responses are color-coded by $[Ca^{2+}]_{bath}$ with the averaged response in black. **I)** PPR curves across ISIs for 1 and 2 mM $Ca^{2+}$ for Bouton #5. **J,K)** Bouton report for Bouton #9 as in **H,I**..

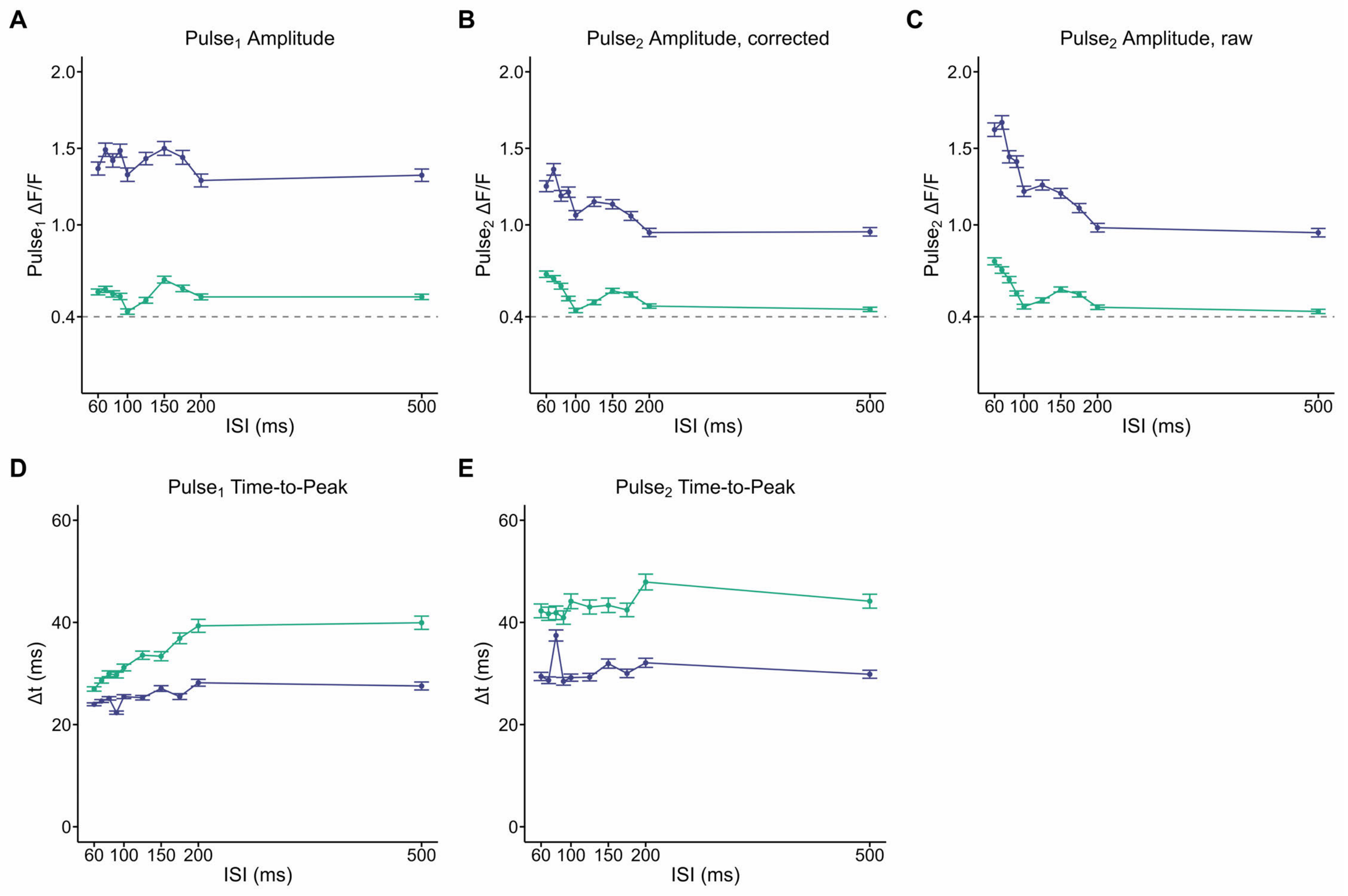


**Figure S7. Basic peak properties for paired-pulse experiment from main-text Figure 6. A)** $Pulse_1$ amplitudes as a function of ISI. There is little jitter across ISI, consistent with a stable initial $P_r$ across the experiment. Dashed line indicates the amplitude of the average iGluSnFR3 response to a putative single SV. **B)** $Pulse_2$ amplitudes as a function of ISI, corrected by subtracting the average iGluSnFR3 response to a single action potential for each synapse from its corresponding paired-pulse traces. The shape of the $Pulse_2$ amplitude curve implies short-term plasticity mechanisms are engaged in a frequency-dependent manner. **C)** Uncorrected $Pulse_2$ amplitudes as a function of ISI. Note that compared to *B*, $Pulse_2$ amplitudes at longer ISIs are unaffected. **D)** $Pulse_1$ time-to-peak. Consistent with observations from main-text **Figure 5**, iGluSnFR3 transients peak more rapidly following stimulus at 2 mM $Ca^{2+}$ than at 1 mM $Ca^{2+}$. Note that the shorter time-to-peak at short ISIs is likely an artifact of the way iGluSnFR3 were identified (e.g., for ISI = 60 ms, the maximum possible Δt is 60 ms, while for ISI ≥ 200 ms, the maximum possible Δt is 200 ms). **E)** $Pulse_2$ time-to-peak. Consistent with observations from main-text **Figure 5**, iGluSnFR3 transients peak more rapidly following stimulus at 2 mM $Ca^{2+}$ than at 1 mM $Ca^{2+}$.

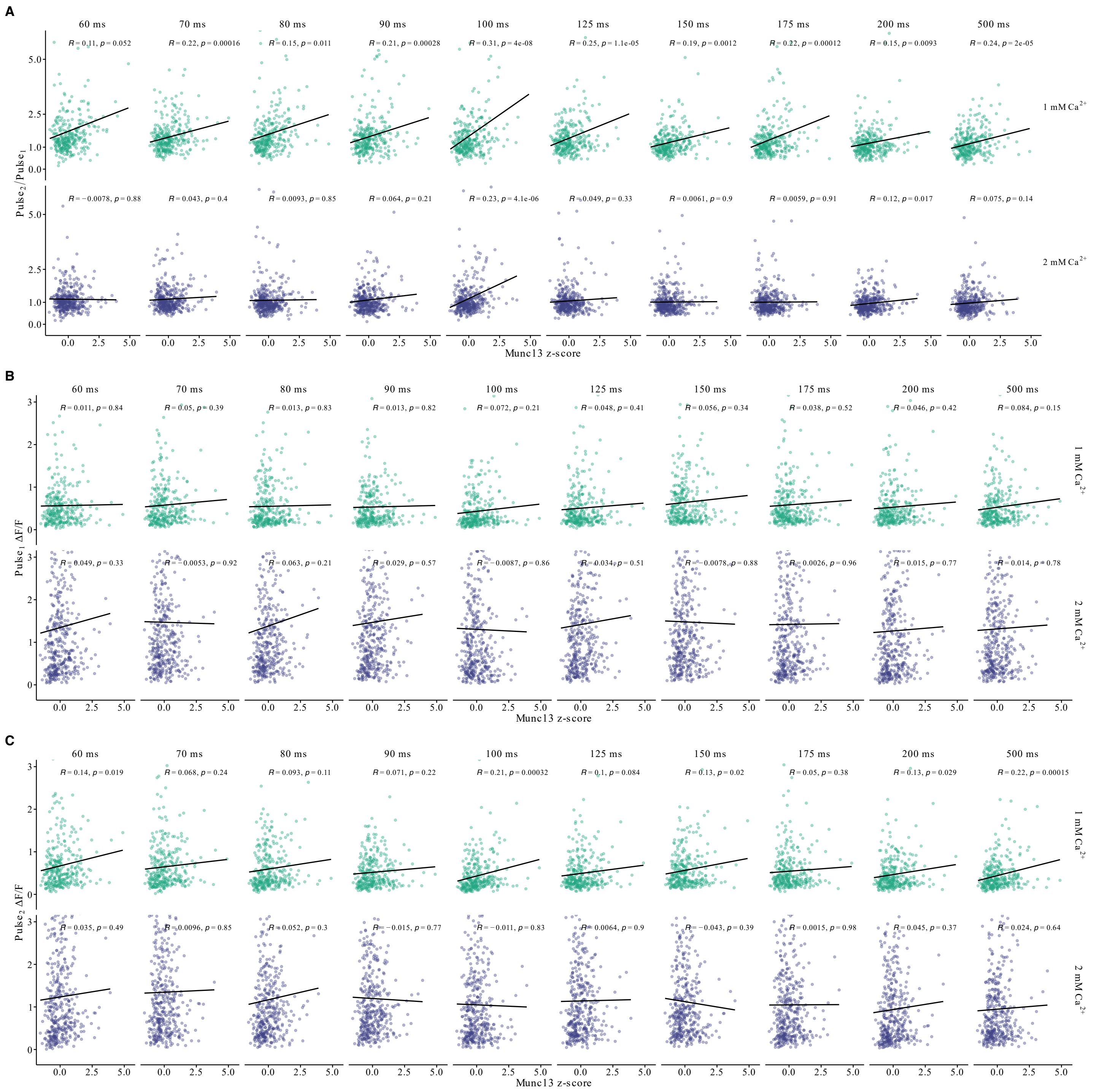


**Figure S8. Dependence of short-term plasticity dynamics on synaptic Munc13 expression for all ISIs. A)** At 1 mM $Ca^{2+}$, PPR positively correlates with Munc13 expression for all ISIs, with strength of Pearson correlation (R value) and statistical significance (p value) varying according to ISI. At 2 mM $Ca^{2+}$, PPR does not correlate with Munc13 expression with ISI = 100 ms as the notable exception. **B)** The amplitude (ΔF/F) of the iGluSnFR3 response to the first stimulus showed little to dependence on Munc13 expression at either $[Ca^{2+}]_{bath}$ tested. **C)** The amplitude (ΔF/F) of the iGluSnFR3 response to the $2^{nd}$ stimulus showed little to no dependence on Munc13 expression at either $[Ca^{2+}]_{bath}$ tested.

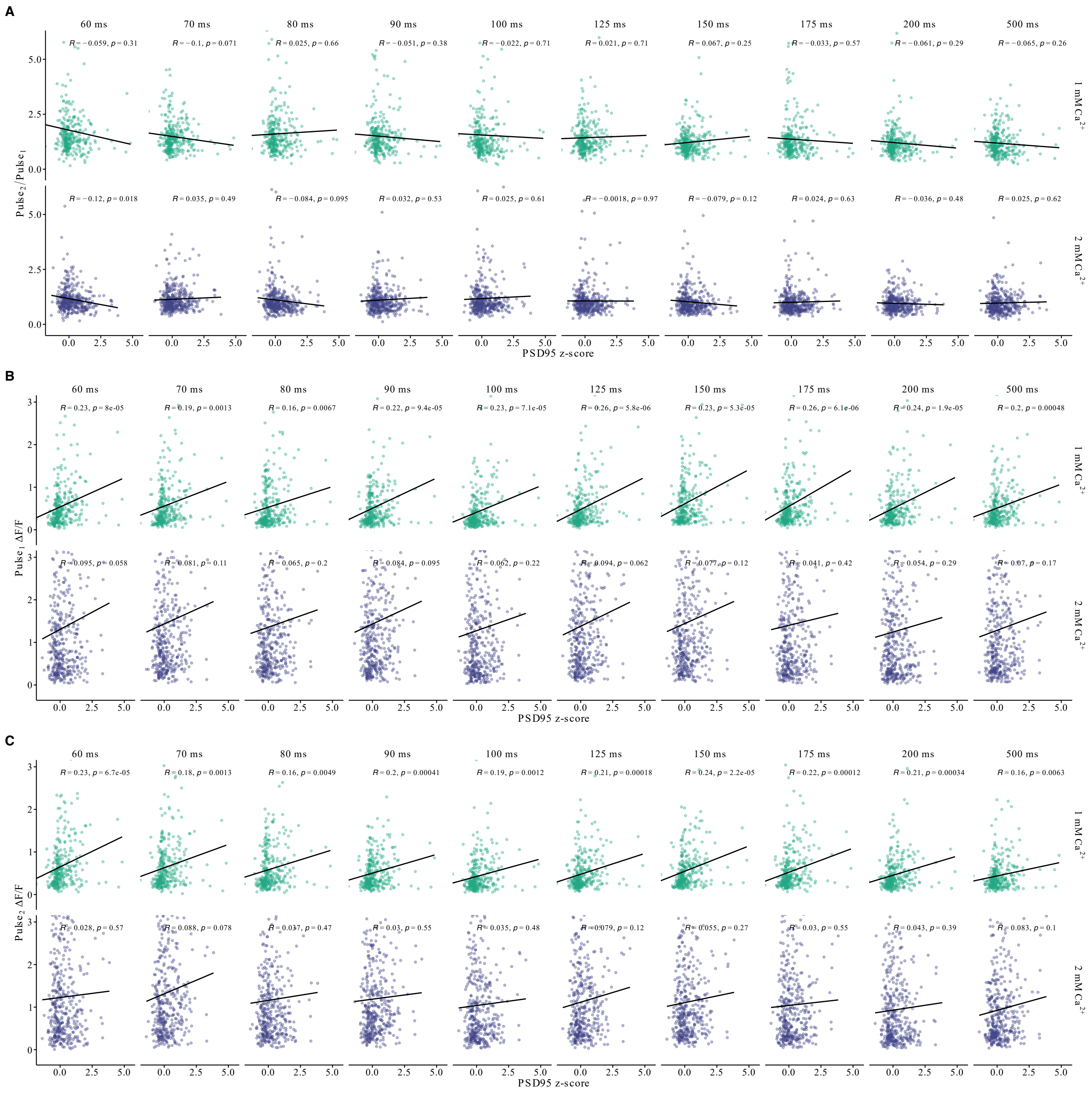


**Figure S9. Dependence of short-term plasticity dynamics on PSD95.FingR.TagRFP expression for all ISIs. A)** PPR did not correlate with PSD95.FingR.TagRFP expression for any ISI at either $[Ca^{2+}]_{bath}$ tested, with strength of Pearson correlation (R value) and statistical significance (p value) varying according to ISI. **B)** The amplitude (ΔF/F) of the iGluSnFR3 response to the first stimulus showed a dependence on PSD95.FingR.TagRFP expression at both $[Ca^{2+}]_{bath}$ tested. **C)** The amplitude (ΔF/F) of the iGluSnFR3 response to the 2nd stimulus showed substantial dependence on PSD95.FingR.TagRFP expression, but only at 1 mM $Ca^{2+}$.

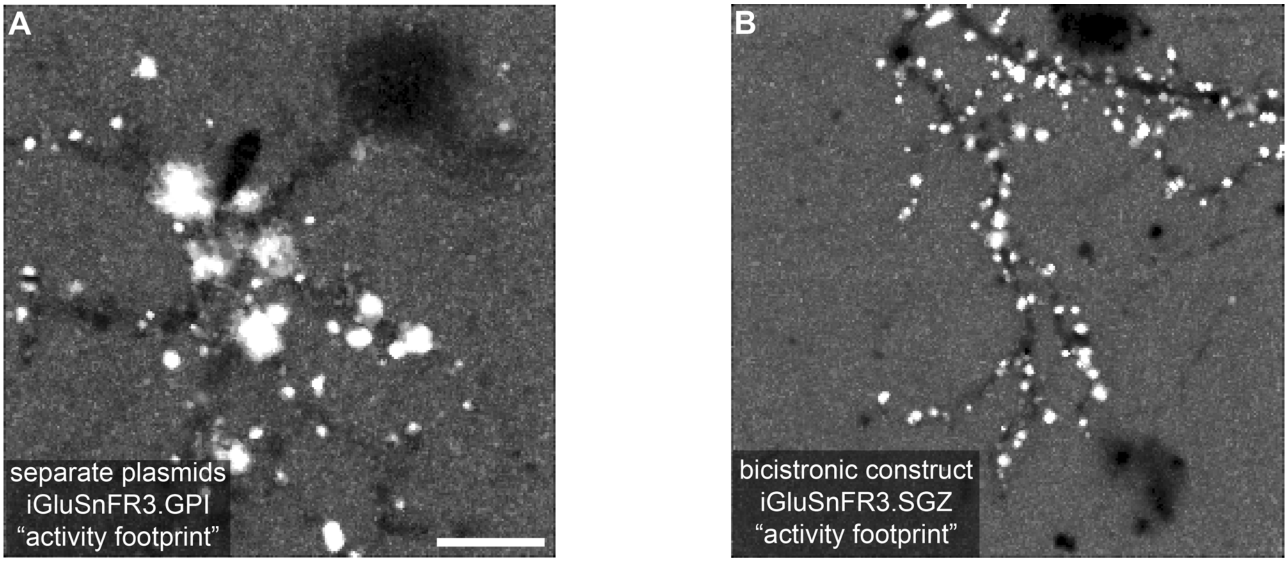


**Figure S10. Improved trafficking of iGluSnFR3.SGZ into the synapse improves activity-segmentation of single spines versus iGluSnFR3.GPI. A)** Activity-based segmentation of iGluSnFR3 signal collected from a neuron expressing spine-HaloTag and the GPI-anchored version of iGluSnFR3 (iGluSnFR3.GPI) from separate plasmids. While some individual spines are extracted for analysis, GPI-anchored iGluSnFR3 trafficks poorly into the synapse, limiting synaptic resolution and decreasing throughput. Scale bar = 20 μm. **B)** Activity-based segmentation of the stargazin(SGZ)-anchored version of iGluSnFR3 signal collected from a neuron expressing a single, bicistronic construct: spine-HaloTag-P2A-iGluSnFR3.SGZ. SGZ-anchored iGluSnFR3 avidly trafficks into the synapse, increasing synaptic resolution and throughput.

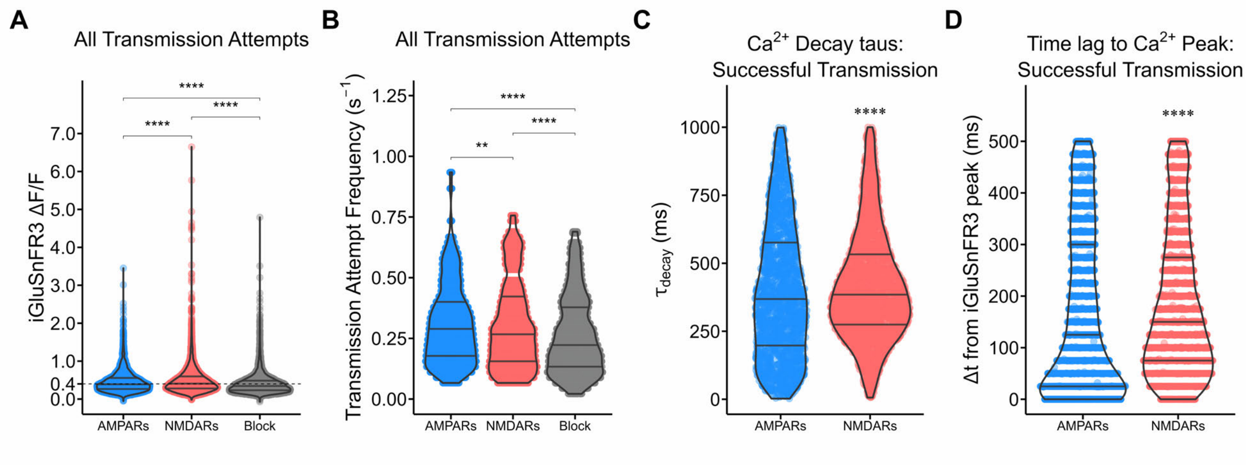


**Figure S11. Statistical properties of all-optical synaptic transmission recordings (main-text Figure 8,9). A)** iGluSnFR3 amplitude across all transmission attempts. Main-text **Figure 8** shows that iGluSnFR3 amplitudes for successful synaptic transmission events were statistically indistinguishable between the "AMPARs open" and "NMDARs open" stages of the experiment. However, when all synaptic transmission attempts are considered (inclusive of failures and successes), we do resolve differences in iGluSnFR3 amplitude. It remains unclear why iGluSnFR3 event amplitude increases slightly following APV washout (note the shift in the top quartile), and then decreases following iGluR blockade. **B)** Transmission attempt frequency by imaging phase. While some synapses appear to increase their transmission frequency following APV washout (red), most synapses experienced a rundown in spontaneous release frequency over the course of the experiment. **C)** $Ca^{2+}$ event $\tau_{decay}$ from single-exponential decay fitting among successful transmission events. **D)** Time delay to $Ca^{2+}$ peak from preceding iGluSnFR3 event among successful transmission events. On average, AMPARs quickly reached their peak amplitude following iGluSnFR3 events, indicating fast receptor kinetics upon successful receptor activation. By contrast, NMDARs took longer to peak, possibly due to saturation of $JF_{646}$-BAPTA dye or more permissive receptor activation dynamics due to their higher affinity for glutamate. Note that the quantization of the Δt values is an inherent property of our sampling frequency (~40 Hz).
